# Phagocytic activity of perivascular cell promotes CNS injury pathology

**DOI:** 10.64898/2026.08.03.742479

**Authors:** Tian Zhou, Zhenwei Zhang, Fanzhuo Zeng, Hao Song, Zhen Xu, Zhenyu Hu, Luming Yao, Wanying Wang, Tianlei Zhang, Xiaoyan Du, Kexiang Li, Zaichao Xie, Yu Sun, Baiqing Ren, Benji Fan, Shaoda Qi, Ying Li, Yifan Hu, Miao Huang, Yutong Chen, Qing Wang, Na Zhao, Maryam Ayazi, Shoujun Yu, Nengyuan Hu, Haitao Sun, Lisen Sui, Kaibin Huang, Qingming Qu, Qiang Liu, Frank W. Pfrieger, Xiong Cao, Chen-Song Zhang, Kairui Mao, Bo Wang, Zuliang Jie, Guojun Bu, Feng Mei, Timothy Megraw, Liping Wang, Yi Ren, Yiming Zheng

## Abstract

Injury, stroke, and neurological diseases cause persistent accumulation of cellular debris that deteriorates lesion microenvironment and impedes central nervous system (CNS) repair. Debris clearance in the injured CNS has long been attributed primarily to microglia and infiltrating macrophages. Here, we identify perivascular cells as previously unrecognized phagocytes that expand after injury and exhibit robust phagocytic activity. Perivascular cell phagocytosis is conserved across multiple mouse models of CNS injury and human stroke lesions. These cells exhibit key hallmarks of phagocytosis, including LC3-associated phagocytosis for efficient lysosomal degradation. Mechanistically, phosphatidylserine serves as the ‘eat-me’ signal and Axl mediates myelin debris uptake. Myelin phagocytosis drives perivascular cell proliferation, fibrosis and lesion progression. Genetic deletion of Axl in perivascular cells or pharmacological inhibition with the FDA-approved Axl inhibitor Gilteritinib reduces pathology and improves functional recovery after spinal cord injury. Together, these findings establish Axl-dependent perivascular cell phagocytosis as a therapeutic target for CNS repair.

## Introduction

Traumatic injury, ischemic stroke and neurological diseases damage the central nervous system (CNS) by creating a hostile microenvironment that initiates secondary pathological processes leading to irreversible neuronal dysfunction and death and impaired CNS repair ^1^. While the immune responses of resident microglia and infiltrating macrophages after CNS injury have been well-characterized ^2–5^, the cellular landscape within CNS lesions is likely far more intricate. Beyond microglia and macrophages, early morphometric studies ^6–10^ and recent high-throughput cellular heterogeneity analyses ^11–13^ have revealed rapid and substantial changes in blood vessels and vasculature-related cell types following CNS injury. However, the mechanisms that initiate and coordinate this neglected response are not yet fully understood.

A particularly intriguing subset of vasculature-related cells in this context are the perivascular cells, which are positioned between endothelial cells, astrocytes, and neurons within the neurovascular unit ^14–16^. These perivascular cells or those of perivascular origin exhibit dramatic increase in conditions associated with CNS fibrosis, including spinal cord injury (SCI), stroke, traumatic brain injury and autoimmune disease ^17–22^. In the healthy CNS, perivascular cells are associated with blood vessels of diverse diameters and they populate all vascular and microvascular beds from arterioles/venules, precapillary arterioles/venules to capillaries ^23–25^. Nevertheless, the precise distribution, identity and functions of distinct perivascular cell populations and subtypes are not fully understood and subject of ongoing debate. Research has largely focused on the physiological roles of perivascular cells in maintaining vascular integrity, including blood-brain barrier function, angiogenesis, and cerebral blood flow regulation^16,26–30^. Despite their strategic localization within neurovascular units, it remains unknown whether and how perivascular cells actively respond to pathologic conditions due to CNS injury, ischemia and disease.

In this study, we aimed to elucidate the contribution of perivascular cells to noxious microenvironments that arise following CNS injury. Using various animal models of CNS demyelinating pathologies and primary cell co-culture models, we uncovered a previously unappreciated role for perivascular cells in directly sensing and reacting to CNS injury. These cells acquire robust phagocytic and degradative capacity, enabling efficient engulfment of myelin debris, a major pathological component of CNS demyelination and injury ^31–34^. The engulfment of myelin debris by these cells has a negative impact as it induces pro-proliferative and pro-fibrotic states in perivascular cells that impede functional recovery, reminiscent of the diseased-associated states described in macrophages, microglia and astrocytes as recently demonstrated ^35–37^. These findings challenge the long-standing view that innate immune responses after CNS injury are primarily mediated by resident microglia and infiltrated macrophages. This study thus provides new insights into perivascular-immune mechanisms in CNS injury, highlighting the therapeutic potential of targeting perivascular cell function to promote functional recovery following injury, stroke and disease.

## Results

### Perivascular cells expand within demyelinating lesions after spinal cord injury

To investigate perivascular cell responses to CNS injury, we employed two complementary genetic labeling strategies based on transgenic mice expressing constitutive PDGFRβ-Cre and tamoxifen-inducible GLAST-CreER^T2^, which target perivascular cell populations including pericytes, perivascular fibroblasts, and vascular smooth muscle cells ^22,23,38^ (**Fig. 1A**). Crossing PDGFRβ-Cre mice with the Ai14 reporter line expressing tdTomato from the *ROSA26* locus revealed efficient and highly specific labeling of spinal cord perivascular cells, with negligible off-target labeling of microglia, astrocytes, or neurons (**Fig. S1A-I**). Following spinal cord injury (SCI), *Pdgfrb* expression remained largely restricted to perivascular cells throughout the SCI course, as confirmed by analysis of published single-cell RNA-sequencing datasets ^12,13^ and immunohistochemical staining of spinal cord sections from PDGFRβ-Cre::Ai14 mice with cell-specific markers (**Fig. S2A–G**). These findings establish PDGFRβ as a reliable marker for tracking perivascular cells in the injured spinal cord.

**Figure 1.**
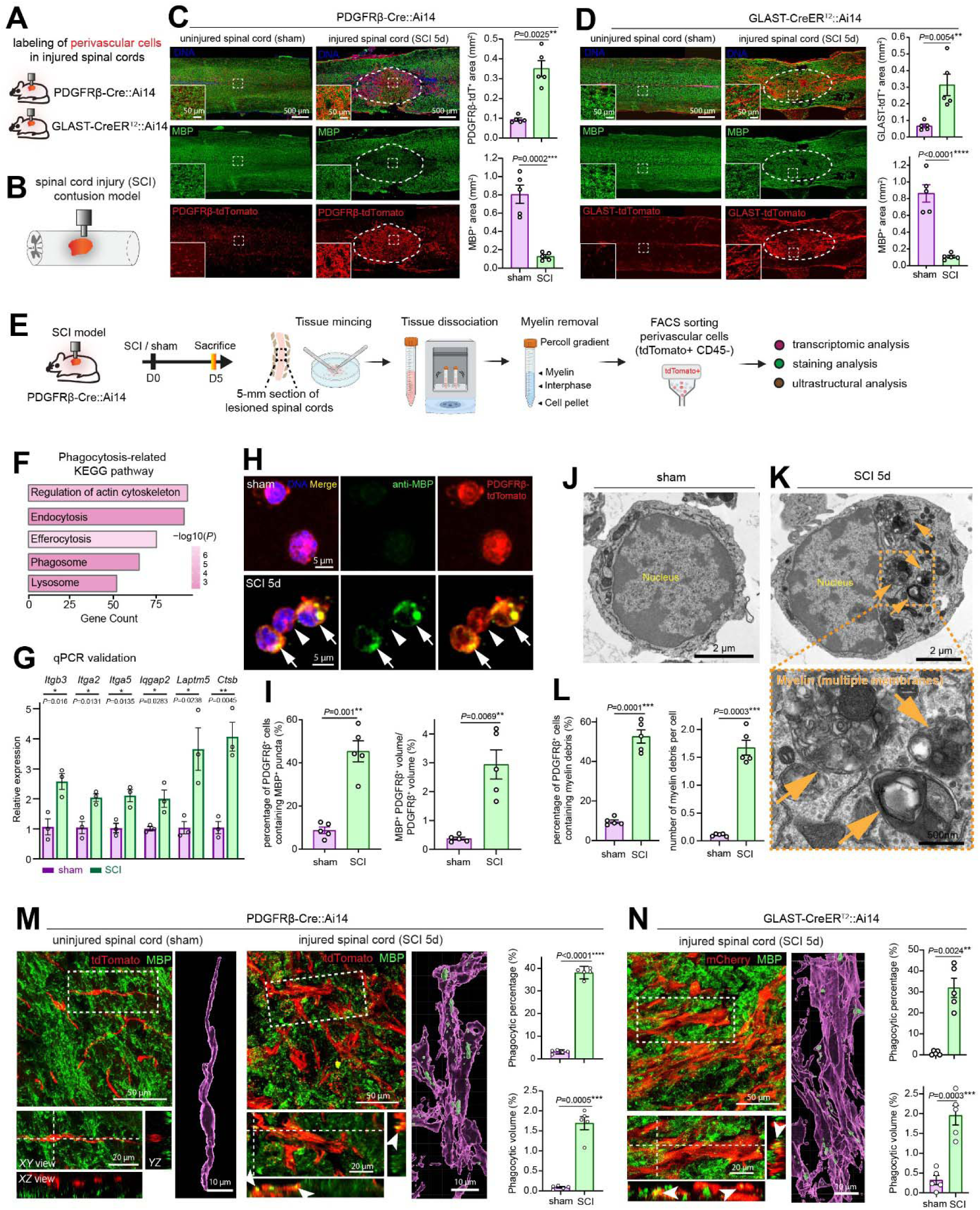
Identification of perivascular cells as phagocytes after spinal cord injury. **(A)** Schematic illustrating genetic labeling of perivascular cells in the injured CNS using PDGFRβ-Cre or tamoxifen-inducible GLAST-CreER^T2^ mice crossed with the R26R-loxP-STOP-loxP-tdTomato (Ai14) reporter strain. **(B)** Diagram of the mouse contusive spinal cord injury (SCI) model. The lesion core at thoracic level T9-10 is indicated in orange. **(C)** Representative confocal images and quantification showing PDGFRβ-tdTomato-labeled perivascular cells and myelin (MBP) in spinal cords 5 days after SCI. Dashed outlines indicate the lesion core, characterized by accumulation of PDGFRβ-tdTomato-positive cells and loss of MBP immunoreactivity. Insets show enlarged views of the indicated regions. Scale bar, 500 μm. Quantification shows the PDGFRβ-tdTomato-positive area and MBP-positive area. Data are presented as mean ± SEM (*n* = 5 mice). Statistical significance was determined by an unpaired two-tailed *t* test (for MBP-positive area) or an unpaired two-tailed *t* test with Welch’s correction (for PDGFRβ-tdTomato-positive area). **(D)** Representative confocal images and quantification showing GLAST-tdTomato-labeled perivascular cells and myelin (MBP) in spinal cords 5 days post SCI. Dashed outlines indicate the lesion core, characterized by accumulation of GLAST-tdTomato-positive cells and loss of MBP immunoreactivity. Insets show enlarged views of the indicated regions. Scale bar, 500 μm. Quantification shows the GLAST-tdTomato-positive area and MBP-positive area. Data are presented as mean ± SEM (*n* = 5 mice). Statistical significance was determined by an unpaired two-tailed *t* test. **(E)** Workflow for fluorescence-activated cell sorting (FACS) isolation of perivascular cells (PDGFRβ-tdTomato^+^CD45^−^) from injured spinal cords followed by transcriptomic profiling, immunocytochemistry, and transmission electron microscopy. **(F)** KEGG pathway analysis of RNA-sequencing data showing enrichment of phagocytosis-related pathways in perivascular cells isolated from SCI mice. **(G)** qRT-PCR validation of phagocytosis-related genes upregulated in perivascular cells isolated from SCI mice. Gene expression was normalized to *Gapdh*. Data are presented as mean ± SEM (*n* = 3 biological replicates). Statistical significance was determined by an unpaired two-tailed *t* test. **(H)** Representative confocal images and quantification showing MBP-positive myelin particles within PDGFRβ-tdTomato-positive perivascular cells isolated from SCI mice 5 days post injury. Arrow and arrowhead indicate cells containing or lacking intracellular myelin particles, respectively. Scale bar, 5 μm. **(I)** Quantification showing the percentage of perivascular cells containing myelin particles (left) and the relative volume of intracellular myelin particles (right). Data are presented as mean ± SEM (*n* = 5 biological replicates). Statistical significance was determined by unpaired two-tailed *t* test with Welch’s correction. **(J–L)** Representative transmission electron micrographs and quantification showing intracellular myelin structures (arrows) in perivascular cells isolated from sham-operated mice and SCI mice 5 days post injury **(J, K)**. Scale bar, 2 μm; 500 nm (enlarged views). **(L)** Quantification showing the percentage of cells containing myelin structures and the number of intracellular myelin structures per cell. Data are presented as mean ± SEM (*n* = 5 biological replicates; 178 cells from sham mice and 196 cells from SCI mice). Statistical significance was determined by an unpaired two-tailed *t* test with Welch’s correction. **(M)** Representative confocal images and quantification showing myelin particle uptake by PDGFRβ-tdTomato-labeled perivascular cells *in vivo*. In sham-operated spinal cords, perivascular cells and myelin sheaths were spatially separated. Enlarged views show orthogonal XY, XZ, and YZ projections demonstrating intracellular localization of MBP-positive myelin particles. Right, Imaris 3D reconstructions of intracellular myelin particles. PDGFRβ-tdTomato is pseudo-colored in the 3D reconstructions. Scale bar, 50 μm (overview), 20 μm (orthogonal views), and 10 μm (3D reconstructions). Quantification shows phagocytic percentage, or the percentage of phagocytic perivascular cells (MBP^+^ PDGFRβ^+^ cells/ total PDGFRβ^+^ cells) and phagocytic volume, or the relative intracellular myelin volume (MBP^+^ PDGFRβ^+^volume/ PDGFRβ^+^ volume). Data are presented as mean ± SEM (*n* = 5 mice). Statistical significance was determined by an unpaired two-tailed *t* test (for phagocytic percentage) or an unpaired two-tailed *t* test with Welch’s correction (for phagocytic volume). **(N)** Representative confocal images and quantification showing myelin particle uptake by GLAST-tdTomato-labeled perivascular cells in vivo. Boxed regions are enlarged to show orthogonal XY, XZ, and YZ projections and Imaris 3D reconstructions. GLAST-tdTomato is pseudo-colored in the 3D reconstructions. Scale bar, 50 μm (overview), 20 μm (orthogonal views), and 10 μm (3D reconstructions). Quantification shows phagocytic percentage (MBP^+^ GLAST^+^ cells/ total GLAST^+^ cells) and phagocytic volume (MBP^+^ GLAST^+^ volume/ GLAST^+^ volume). Data are presented as mean ± SEM (*n* = 5 mice). Statistical significance was determined by an unpaired two-tailed *t* test correction (for phagocytic volume) or an unpaired two-tailed *t* test with Welch’s correction (for phagocytic percentage).

We next examined the spatiotemporal dynamics of perivascular cells following SCI. Consistent with previous reports ^17,20^, the number of PDGFRβ⁺ cells increased markedly by 5 days post injury compared to control mice, which underwent sham operation without injury (**Fig. 1C**). Notably, unlike reactive astrocytes and microglia, which predominantly localize to the lesion border ^31,39,40^, the expanded PDGFRβ⁺ cells accumulated within the demyelinated lesion core, defined by the diminished myelin basic protein (MBP) staining (**Fig. 1B, C**). The preferential enrichment of perivascular cells within demyelinated regions suggest a previously unappreciated association between these cells and myelin debris after SCI.

To independently validate these observations, we employed the GLAST-CreER^T2^ lineage tracing system ^41^, which has been extensively used as a reliable tool to label injury-responsive perivascular cells after CNS injury ^17,20,22^. Previous lineage-tracing and transcriptomic analyses demonstrated that GLAST-expressing cells comprise the majority of PDGFRβ⁺ pericytes and perivascular fibroblasts in injured spinal cord tissue, with minimal expression of endothelial, glial, neural precursor, or myeloid lineage markers (**Fig. S3A**). Consistent with these reports, tamoxifen-induced labeling using GLAST-CreER^T2^::Ai14 mice selectively marked PDGFRβ⁺ and α-SMA⁺ perivascular cells in the spinal cord, representing 80–90% of the total perivascular cell population after SCI, while excluding major glial populations (**Fig. S3B–G**).

Importantly, GLAST-expressing perivascular cells recapitulated the response observed with PDGFRβ lineage tracing, undergoing robust expansion within the demyelinated lesion core by 5 days post injury (**Fig. 1D**). Thus, two independent genetic labeling strategies converge on the same conclusion that perivascular cells undergo marked expansion after SCI and preferentially accumulate within regions where myelin debris accumulates.

### Identification of perivascular cells as phagocytes after spinal cord injury

The preferential accumulation of perivascular cells within demyelinated lesions prompted us to ask whether and how these cells respond to myelin debris after SCI. To define their injury-induced molecular features, we performed RNA sequencing of perivascular cells that were acutely isolated from injured and sham control spinal cords by fluorescence activated cell sorting (FACS) (**Fig. 1E, Fig. S4A**). Differential expression analysis revealed broad transcriptional remodeling of these cells after SCI, including enrichment of pathways associated with phagocytosis, cell proliferation, inflammation, and extracellular matrix remodeling (**Fig. S4B–D**). Notably, genes involved in actin cytoskeletal dynamics, endocytosis, efferocytosis, phagosome formation, and lysosomal function were among those prominently upregulated and were independently validated by qPCR (**Fig. 1F, G; Fig. S4B, C**). These findings indicate that perivascular cells display a phagocytic transcriptional program following CNS injury.

To determine whether perivascular cells actively engulf myelin debris, we first examined perivascular cells that were acutely sorted from spinal cord tissues of PDGFRβ-Cre::Ai14 mice by FACS. Whereas perivascular cells from uninjured mice contained little to no myelin basic protein (MBP), cells isolated from injured spinal cords harbored abundant intracellular MBP-positive material (**Fig. 1H, I**). Ultrastructural analysis by transmission electron microscopy further revealed intracellular multilamellar myelin structures within perivascular cells isolated from the injured spinal cords (**Fig. 1J–L**; **Fig. S5**). These findings provide direct evidence that perivascular cells internalize myelin after SCI.

We next assessed myelin uptake *in vivo*. In the uninjured spinal cord, PDGFRβ⁺ perivascular cells remained spatially segregated from intact myelin and lacked detectable intracellular MBP (**Fig. 1M, Movie 1**). Following SCI, however, PDGFRβ⁺ cells contained abundant myelin debris, as confirmed by orthogonal imaging and three-dimensional (3D) reconstruction of engulfed material (**Fig. 1M, Fig. S6, Movie 1**). Similar myelin uptake was observed in GLAST-lineage-traced perivascular cells using the GLAST-CreER^T2^::Ai14 line (**Fig. 1N**), indicating that this phenomenon is not restricted to a single genetic labeling strategy. Together, these findings identify perivascular cells as previously unrecognized phagocytes that actively engulf myelin debris after SCI.

### Perivascular cell phagocytosis is conserved across CNS demyelinating conditions in mice and humans

Because demyelination is a common feature of diverse pathologic conditions ^37,42–46^, we next asked whether perivascular cell phagocytosis extends beyond SCI. To address this question, we examined a transient middle cerebral artery occlusion (MCAO) model of ischemic stroke in mice (**Fig. 2A, B**). Similar to SCI, ischemic lesions exhibited marked expansion of PDGFRβ⁺ perivascular cells within demyelinated regions (**Fig. 2C**). Notably, perivascular cells in infarcted brain tissue contained abundant intracellular myelin debris, whereas little to no myelin uptake was detected in uninjured controls (**Fig. 2D–F**; **Movie 2**). These findings indicate that perivascular cell phagocytosis is not restricted to traumatic injury but is also a prominent feature of ischemic demyelination.

**Figure 2.**
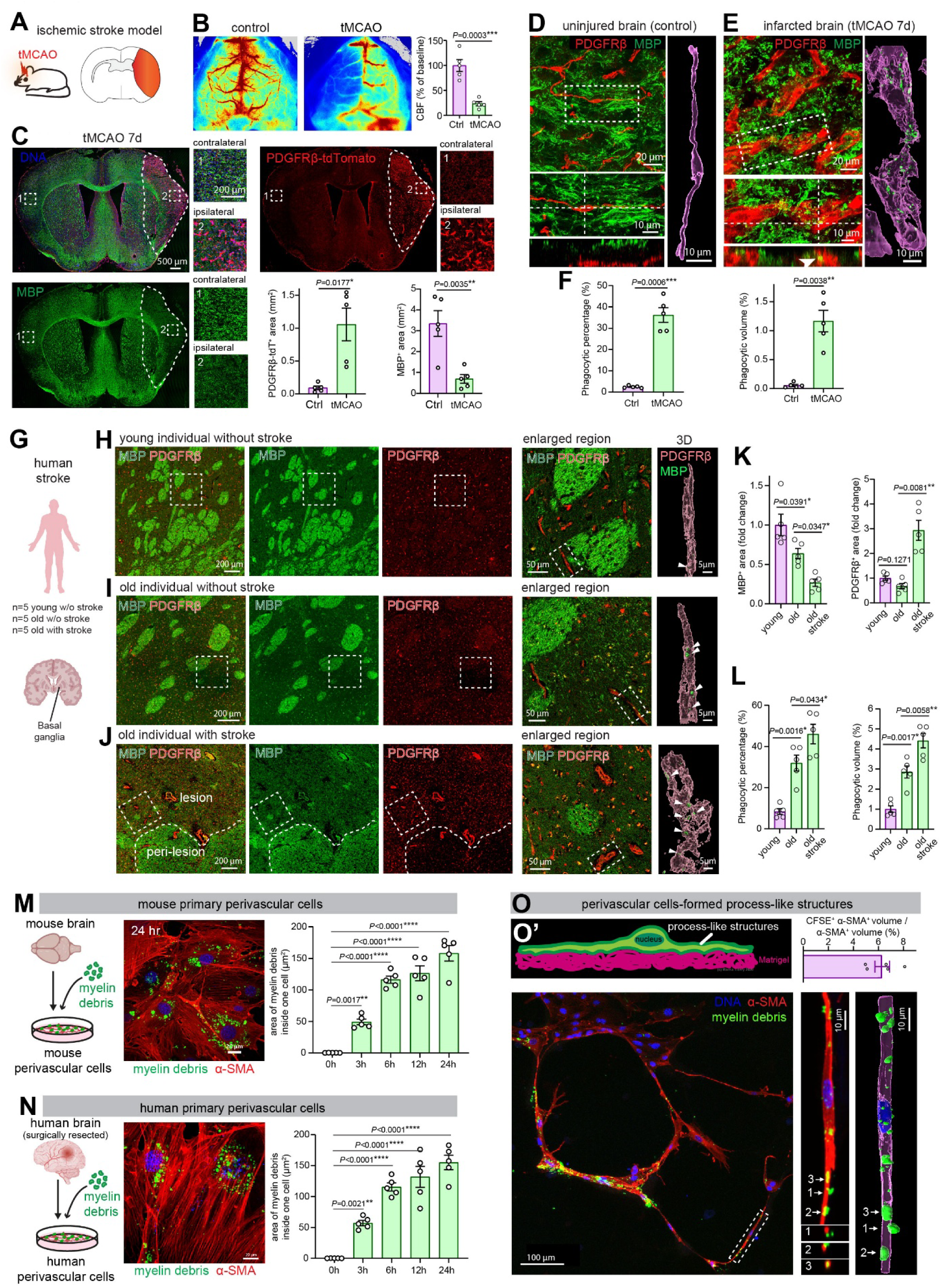
Perivascular cell phagocytosis is a common feature across CNS injury diseases and conserved in mouse and human. **(A)** Schematic of the transient middle cerebral artery occlusion (tMCAO) model of ischemic stroke. The infarcted brain region is indicated in orange. **(B)** Representative laser speckle contrast images showing cerebral blood flow in sham-operated and tMCAO mice. Quantification shows cerebral blood flow quantification, confirming successful induction of ischemic stroke. Data are presented as mean ± SEM (*n* = 5 mice). Statistical significance was determined by an unpaired two-tailed *t* test. **(C)** Representative confocal images showing PDGFRβ-tdTomato-labeled perivascular cells and myelin (MBP) 7 days after tMCAO. The infarcted region (dashed outline) exhibits accumulation of PDGFRβ-tdTomato-positive cells and reduction of MBP immunoreactivity. Enlarged views of the contralateral (**#1**) and ipsilateral (**#2**) regions are shown. Scale bar, 500 μm; 200 μm (enlarged views). Quantification shows the PDGFRβ-tdTomato-positive area and MBP-positive area. Data are presented as mean ± SEM (*n* = 5 mice). Statistical significance was determined by an unpaired two-tailed *t* test (MBP-positive area) or an unpaired two-tailed *t* test with Welch’s correction (PDGFRβ-tdTomato-positive area). **(D, E)** Representative confocal images showing perivascular cells (PDGFRβ, red) and myelin (MBP, green) in uninjured **(D)** and tMCAO **(E)** brains. In uninjured brains, perivascular cells and myelin sheaths are spatially separated, whereas PDGFRβ-positive cells contain intracellular MBP-positive myelin particles (arrowheads) 7 days after tMCAO. Boxed regions are enlarged to show orthogonal XY projections (bottom) and Imaris 3D reconstructions (right). PDGFRβ is pseudo-colored in the 3D reconstructions. Scale bar, 20 μm; 10 μm (orthogonal views and 3D reconstructions). **(F)** Quantification showing phagocytic percentage (MBP^+^ PDGFRβ^+^ cells/ total PDGFRβ^+^ cells) and phagocytic volume (MBP^+^ PDGFRβ^+^ volumes/ PDGFRβ^+^ volumes). Data are presented as mean ± SEM (*n* = 5 mice). Statistical significance was determined by an unpaired two-tailed *t* test with Welch’s correction. **(G)** Schematic illustrating the human brain samples and regions analyzed. **(H–J)** Representative confocal images showing perivascular cells (PDGFRβ, red) and myelin (MBP, green) in brain samples from young individuals without stroke **(H)**, aged individuals without stroke **(I)**, and aged individuals with ischemic stroke **(J)**. Stroke lesions exhibit increased PDGFRβ-positive cell accumulation and reduced MBP immunoreactivity. Dashed lines delineate the infarct core from the peri-lesion region. Enlarged views and Imaris 3D reconstructions demonstrate intracellular MBP-positive myelin particles within perivascular cells (arrowheads). Scale bar, 200 μm (overview), 50 μm (enlarged views), and 5 μm (3D reconstructions). **(K)** Quantification showing the relative MBP-positive area and PDGFRβ-positive area in human brain samples, which was normalized to young individual group. Data are presented as mean ± SEM (*n* = 5 human brain samples). Statistical significance was determined by one-way ANOVA with Tukey’s post hoc test (for MBP-positive area) or Welch’s ANOVA with Dunnett’s post hoc test (PDGFRβ-positive area). **(L)** Quantification showing phagocytic percentage (MBP^+^ PDGFRβ^+^ cells/ total PDGFRβ^+^ cells) and phagocytic volume (MBP^+^ PDGFRβ^+^volumes/ PDGFRβ^+^ volumes). Data are presented as mean ± SEM (*n* = 5 human brain samples). Statistical significance was determined by one-way ANOVA with Tukey’s post hoc test. **(M, N)** Schematics (left) and representative confocal images (right) showing uptake of CFSE-labeled myelin debris by primary mouse **(M)** and human **(N)** brain perivascular cells. Human perivascular cells were isolated from surgically resected epileptic brain tissue. Cells were stained for α-SMA and analyzed at the indicated time points following myelin exposure. Scale bar, 20 μm. Quantification shows the area of engulfed myelin debris over time. Data are presented as mean ± SEM (*n* = 5 biological replicates). Statistical significance was determined by one-way ANOVA with Dunnett’s post hoc test. **(O)** Top, schematic illustrating the process-like morphology of perivascular cells cultured on Matrigel (**O**′). Bottom, representative confocal images showing uptake of CFSE-labeled myelin debris by thin cellular processes of α-SMA-positive perivascular cells. Enlarged regions (1–3) are shown as orthogonal XY and XZ projections together with Imaris 3D reconstructions. Scale bar, 100 μm (left) and 10 μm (right). Quantification shows the relative intracellular myelin volume normalized to total α-SMA-positive cell volume. Data are presented as mean ± SEM (*n* = 5 biological replicates).

To determine whether this response is conserved in humans, we analyzed postmortem basal ganglia samples from stroke patients together with age-matched and young control individuals (**Fig. 2G**; **Fig. S7A**). Consistent with our previous observations ^47^, stroke lesions exhibited evident myelin loss, which was accompanied by a marked increase in perivascular cell density (**Fig. 2H–K**). Importantly, perivascular cells within these lesions contained a considerable amount of intracellular myelin debris, closely mirroring the phenotype observed in the mouse MCAO model (**Fig. 2L**). Aging is one of the key risk factors for CNS demyelination ^45,46,48^. Aged individuals without stroke also exhibited modest myelin degeneration and detectable myelin uptake by perivascular cells, although to a lesser extent than stroke patients (**Fig. 2L**). Similarly, PDGFRβ⁺ perivascular cells in aged mouse brains actively engulfed myelin debris (**Fig. S7B–D**; **Movie 3**). Together, these findings establish that perivascular cell phagocytosis is conserved in both mice and humans and is shared across injury-induced and age-associated demyelination.

We next asked whether perivascular cell phagocytosis also occurs in autoimmune demyelinating disease. In experimental autoimmune encephalomyelitis (EAE), a mouse model of multiple sclerosis, demyelinating lesions were evident by 14 days after immunization, as indicated by reduced MBP signal (**Fig. S8B, C**) and presence of degraded MBP (dMBP) (**Fig. S8D, E**), which was detected by an antibody that recognizes specific epitope in degenerated myelin ^49^. Demyelinating lesions were associated with infiltration of inflammatory cells and accumulation of perivascular cells (**Fig. S8A–E**). Despite their distinct etiology and lesion architecture, EAE lesions likewise accumulated PDGFRβ⁺ perivascular cells that exhibited evident engulfment of myelin debris (**Fig. S8F, G**; **Movie 4**). Thus, perivascular cell phagocytosis emerges as a shared feature of traumatic, ischemic, age-related, and autoimmune demyelination.

To further examine the conservation and dynamics of perivascular cell phagocytosis, we established primary mouse and human perivascular cell cultures using a high-yield enzymatic isolation strategy and challenged them with purified myelin debris (**Fig. S9A**). The isolated perivascular cells expressed canonical perivascular markers, including PDGFRβ, α-SMA, NG2, and Desmin, while lacking endothelial, astrocytic, and microglial markers (**Fig. S9B–E**). Both primary mouse and human perivascular cells efficiently engulfed carboxyfluorescein succinimidyl ester (CFSE)-labeled myelin debris *in vitro* in a time-dependent manner (**Fig. 2M, N**). To better recapitulate the elongated morphology of perivascular cells *in vivo*, we further optimized culture conditions on Matrigel to promote the formation of thin process-like extensions (**Fig. 2O**). After coculture with myelin debris, these remodeled process-like structures demonstrate dynamic and robust uptake of myelin debris, with myelin particles observed at various stages of internalization (**Fig. 2O**; **Movie 5**).

Collectively, these findings establish perivascular cells as a previously unrecognized phagocytic population that is broadly engaged across diverse demyelinating conditions and conserved between mice and humans. The remarkable consistency of this perivascular response across multiple experimental models, human pathology, and primary cell cultures indicates that perivascular cell-mediated phagocytosis of myelin debris is a common feature of CNS response to demyelination.

### Perivascular cells exhibit potent and broad phagocytic capacity

To benchmark the phagocytic capacity of perivascular cells, we compared them with macrophages and microglia, the best known phagocytes of the injured CNS. Following either SCI or MCAO, both PDGFRβ⁺ perivascular cells and Iba1⁺ macrophages/microglia actively engulfed myelin debris (**Fig. S10A, D**). Although a smaller fraction of perivascular cells engaged in phagocytosis of myelin debris, their myelin uptake capacity was comparable to that of macrophages/microglia in both injury models (**Fig. S10B–F**). Consistent with these *in vivo* observations, primary perivascular cells efficiently engulfed purified myelin debris *in vitro*. While macrophages displayed more rapid uptake during the initial phase, the phagocytic capacity of perivascular cells progressively increased and became indistinguishable from that of macrophages by 24 hr after myelin addition (**Fig. S10G, H**). Thus, perivascular cells exhibit a phagocytic capacity comparable to that of professional phagocytes albeit with slow kinetics.

We next asked whether perivascular cells are specialized for myelin clearance or whether they engulf a broader range of cargos. To address this question, we examined the uptake of degenerating neurons following ischemic stroke. In our mouse MCAO stroke model, there was significant neuronal degeneration and death as marked by reduced NeuN staining (**Fig. S11A**). Notably, 3D reconstruction from confocal images revealed that approximately 90% of PDGFRβ^+^perivascular cells engulfed NeuN^+^ neuronal bodies, a reactive response that was long thought to occur primarily in macrophages/microglia after stroke (**Fig. S11B**). Remarkably, both the frequency and efficiency of neuronal uptake were comparable between perivascular cells and macrophages (**Fig. S11C, D**). Similar results were obtained *in vitro*, where primary perivascular cells engulfed fluorescently labeled neuronal bodies as efficiently as bone marrow-derived macrophages (**Fig. S11E, F**). Furthermore, perivascular cells readily internalized dead yeast-derived zymosan particles, a structurally distinct phagocytic substrate, at levels comparable to macrophages (**Fig. S11G, H**).

Together, these findings demonstrate that perivascular cells are not merely lesion-associated bystanders but highly potent phagocytes with broad cargo recognition. Their ability to efficiently engulf myelin debris, degenerating neurons, and microbial particles places them among the major phagocytic populations of the injured CNS and suggests a previously unrecognized role in orchestrating tissue remodeling following neurological injury.

### Injury-associated mechanisms underlying perivascular cell phagocytosis

Perivascular cells exhibit little or no detectable phagocytic activity under physiological conditions but readily display phagocytic function following CNS injury. We reasoned that this apparent transition may reflect the exposure of a phagocytic potential that is normally restrained by the neurovascular architecture. Under healthy conditions, perivascular cells are structurally enwrapped by astrocytic end-feet, a key structural component of the blood-brain barrier (BBB) ^50^ (**Fig. S12A, D**), which prevents perivascular cells from accessing neural debris. Following either SCI or MCAO, however, this structural organization was profoundly disrupted ^50^, and astrocytic end-feet coverage was largely absent within the lesion core, leaving perivascular cells directly exposed to cellular debris (**Fig. S12B–G**). These observations suggest that injury-induced disruption of the perivascular niche creates anatomical conditions that expose perivascular cells to myelin debris and other injury-associated substrates.

We next investigated the mechanisms that initiate perivascular cell phagocytosis. To capture the early cellular response to demyelination cues, we performed RNA sequencing of cultured perivascular cells following exposure to myelin debris for 6 hr (**Fig. 3A**). Myelin-engulfing perivascular cells rapidly activated transcriptional programs associated with phagocytosis, including actin cytoskeletal remodeling, efferocytosis, and phagosome formation, together with inflammatory and fibrotic pathways (**Fig. 3B**; **Fig. S13A, B**). Among the most prominently induced genes were members of the integrin family, key regulators of cytoskeletal dynamics and phagocytosis (**Fig. 3C, D**). Thus, myelin debris exposure is sufficient to initiate a phagocytic transcriptional program in perivascular cells.

**Figure 3.**
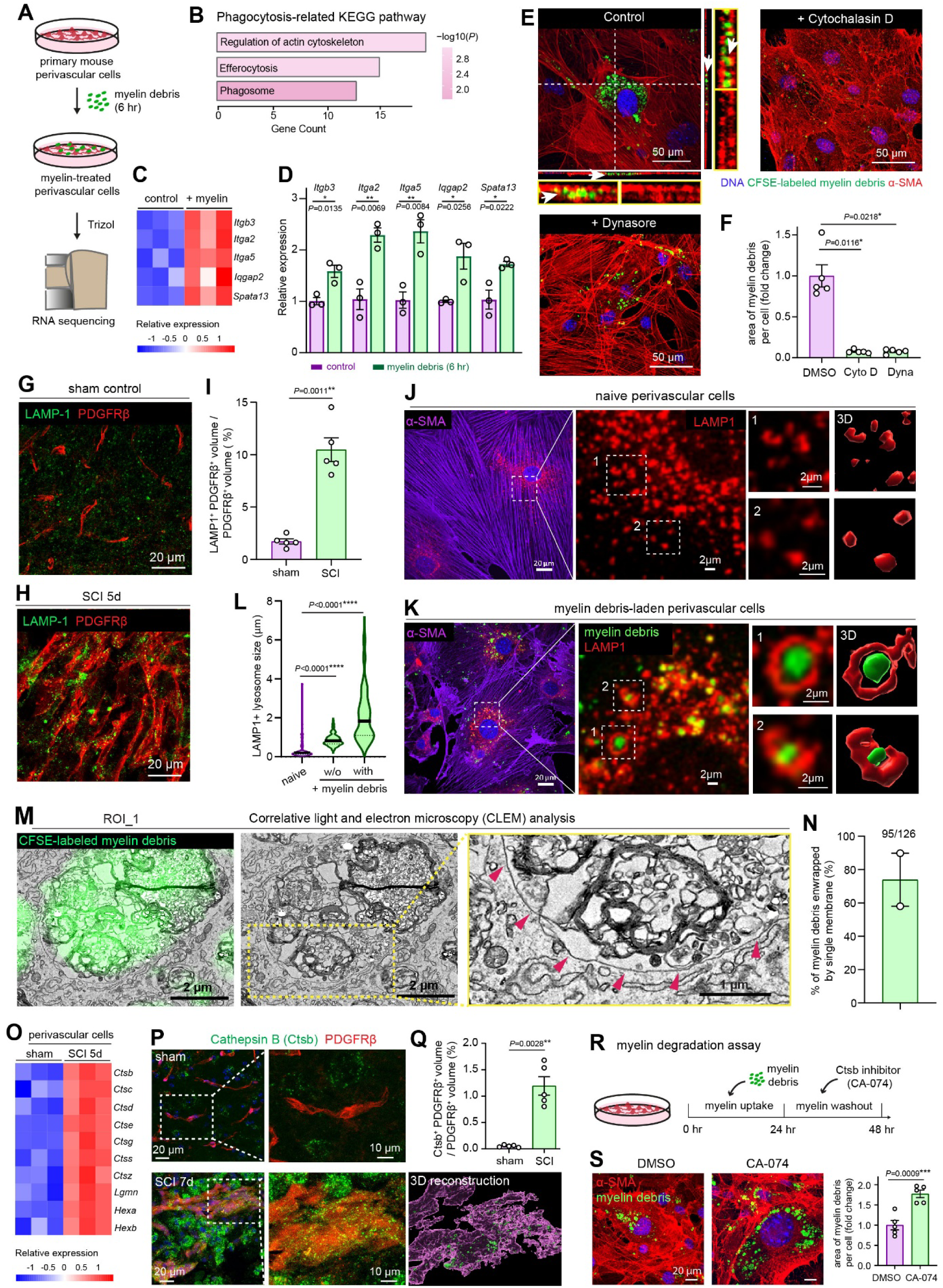
Perivascular cells exhibit the key hallmarks of phagocytosis. **(A)** Schematci illustrating bulk RNA-sequencing analysis of primary perivascular cells following engulfment of myelin debris for 6 hr. **(B)** KEGG pathway analysis of RNA-sequencing data showing enrichment of phagocytosis-related pathways in myelin debris-laden perivascular cells. **(C)** Heatmap showing phagocytosis-related genes upregulated in myelin debris-laden perivascular cells. **(D)** qRT-PCR validation of phagocytosis-related genes upregulated in myelin debris-laden perivascular cells. Gene expression was normalized to *Gapdh*. Data are presented as mean ± SEM (*n* = 3 biological replicates). Statistical significance was determined by an unpaired two-tailed *t* test. **(E, F)** Representative confocal images showing uptake of CFSE-labeled myelin debris by primary perivascular cells in the presence of the actin polymerization inhibitor cytochalasin D (0.5 μM, 24 hr) or the dynamin inhibitor Dynasore (80 μM, 24 hr). Orthogonal XZ and YZ projections demonstrate enclosure of myelin debris by α-SMA-positive cellular structures (arrows). Scale bar, 50 μm. **(F)** Quantification showing the relative intracellular myelin area normalized to DMSO-treated controls. Data are presented as mean ± SEM (*n* = 5 biological replicates). Statistical significance was determined by the Kruskal–Wallis test with Dunn’s post hoc test. **(G, H)** Representative confocal images showing LAMP-1 immunostaining in PDGFRβ-positive perivascular cells from sham-operated and SCI mice 5 days post injury. Scale bar, 20 μm. **(I)** Quantification showing the relative volume of LAMP-1-positive lysosomal compartments normalized to total PDGFRβ-positive cell volume. Data are presented as mean ± SEM (*n* = 5 mice). Statistical significance was determined by an unpaired two-tailed *t* test with Welch’s correction. **(J, K)** Representative confocal images showing LAMP-1-positive lysosomal compartments in untreated (naïve) and myelin debris-laden primary perivascular cells. Enlarged views demonstrate enlarged lysosomal compartments following myelin uptake. Two representative regions are further magnified and reconstructed in 3D to illustrate lysosomal compartments surrounding intracellular myelin debris. Perivascular cells were identified by α-SMA immunostaining. Scale bar, 20 μm (left) and 2 μm (right). **(L)** Quantification showing the diameter of LAMP-1-positive lysosomal compartments associated with or without (w/o) intracellular myelin debris. Data are presented as mean ± SEM (*n* = 5 biological replicates). Statistical significance was determined by the Kruskal–Wallis test with Dunn’s post hoc test. **(M)** Representative correlative light and electron microscopy (CLEM) images showing CFSE-labeled intracellular myelin debris and the surrounding vesicular ultrastructure in primary perivascular cells. Arrows indicate single-membrane vesicles enclosing myelin debris. Scale bar, 2 μm (left and middle) and 1 μm (right). **(N)** Quantification showing the percentage of intracellular myelin debris enclosed by single-membrane vesicles. Data are presented as mean ± SEM (95 of 126 myelin particles from *n* = 2 CLEM cell samples). **(O)** Heatmap showing lysosomal enzyme genes upregulated in FACS-isolated perivascular cells from SCI mice 5 days post injury. **(P, Q)** Representative confocal images showing Cathepsin B (Ctsb) immunostaining in PDGFRβ-positive perivascular cells from sham-operated mice and SCI mice 7 days post injury **(P)**. Enlarged views and Imaris 3D reconstructions demonstrate increased Ctsb expression following SCI. Scale bar, 20 μm (left) and 10 μm (right). **(Q)** Quantification showing the relative Ctsb volume normalized to total PDGFRβ-positive cell volume. Data are presented as mean ± SEM (*n* = 5 mice). Statistical significance was determined by an unpaired two-tailed *t* test with Welch’s correction. **(R)** Schematic illustrating the *in vitro* myelin degradation assay. Following myelin uptake, extracellular myelin debris was removed, and degradation of intracellular myelin was assessed in the presence or absence of the Cathepsin B inhibitor CA-074. **(S)** Representative confocal images showing impaired degradation of CFSE-labeled myelin debris following inhibition of Cathepsin B with CA-074 (4 μM, 24 hr). Scale bar, 20 μm. Quantification shows the relative intracellular myelin area normalized to DMSO-treated controls. Data are presented as mean ± SEM (*n* = 5 biological replicates). Statistical significance was determined by an unpaired two-tailed *t* test.

Phagocytosis in professional phagocytes begins with plasma membrane invagination and formation of actin-rich structures, namely phagocytic cups to encapsulate target particles ^51^. Consistent with this mechanism, perivascular cells formed prominent actin-rich structures surrounding myelin debris, neuronal bodies, and zymosan particles during uptake (**Fig. 3E, Fig. S13C, D**). Pharmacological disruption of actin polymerization with cytochalasin D markedly inhibited engulfment, as did inhibition of the membrane scission GTPase dynamin using Dynasore (**Fig. 3E, F**). These findings establish that perivascular cells employ the canonical cellular machinery of phagocytosis to internalize extracellular substrates.

Phagocytosis is a multistep process that involves phagosome formation, maturation, and fusion with lysosomes to form phagolysosomes for cargo degradation ^52^. We therefore examined whether perivascular cells mobilize lysosomal pathways following engulfment. *In vivo*, SCI induced a significant increase in LAMP1-positive lysosomal structures within perivascular cells (**Fig. 3G–I**). In cultured cells, internalized myelin debris was frequently enclosed within LAMP1-positive compartments that enlarged following debris engulfment (**Fig. 3J–L**), indicative of substantial lysosomal engagement. To more precisely track the intracellular fate of myelin debris in perivascular cells at high resolution, we performed correlative light and electron microscopy (CLEM) (**Fig. S14A, B**). CLEM analysis confirmed that CFSE fluorophore-labeled myelin debris corresponded to ultrastructurally defined intracellular myelin-containing compartments within perivascular cells (**Fig. 3M, Fig. S14C, D**). Notably, approximately 70% of myelin debris were enclosed by single membranes (**Fig. 3N**). These findings indicate that engulfed myelin debris is efficiently delivered to lysosome-associated degradative compartment in perivascular cells.

Consistent with active lysosomal processing, transcriptomic analysis of perivascular cells sorted by FACS from SCI 5d mice revealed upregulation of genes encoding lysosomal enzymes, including cathepsin B (Ctsb) (**Fig. 3O**). Elevated *Ctsb* expression in perivascular cells was independently confirmed in published single-cell datasets from both SCI ^13^ and MCAO ^53^ models and validated at the protein level *in vivo* (**Fig. 3P, Q**; **Fig. S13E–H**). Exposure to myelin debris similarly induced Ctsb expression in cultured perivascular cells, where the enzyme localized to cargo-containing compartments (**Fig. S13I, J**). Functionally, inhibition of Ctsb with CA-074Me impaired degradation of engulfed material (**Fig. 3R, S**), indicating that lysosomal enzyme Ctsb is required for efficient processing of internalized myelin by perivascular cells.

Together, these findings define a multistep program that activates perivascular cell phagocytosis following CNS injury. Injury-induced disruption of astrocytic barriers exposes perivascular cells to tissue debris, which in turn activates a canonical engulfment program characterized by cytoskeletal remodeling, phagosome formation, engagement of lysosome-associated degradative pathways, and enzymatic degradation of internalized myelin debris.

### Non-canonical degradative pathway facilitates degradation of engulfed cargos by perivascular cells

Having established that perivascular cells internalize myelin debris, we next asked how engulfed cargo is trafficked to lysosomes. Canonical autophagy has been implicated in myelin clearance by Schwann cells and endothelial cells ^34,54,55^, raising the possibility that a similar mechanism operates in perivascular cells. Following engulfment of myelin debris or zymosan, both perivascular cells and macrophages generated large LAMP1⁺ vesicular structures that were decorated by LC3 (**Fig. 4A**), a hallmark autophagy protein. Although cargo uptake and rapamycin treatment each increased the abundance of LC3⁺LAMP1⁺ vesicles (**Fig. 4B**), cargo-associated vesicles were substantially larger than rapamycin-induced autophagosomes (**Fig. 4C**). Moreover, myelin debris preferentially localized to these enlarged LC3⁺LAMP1⁺ compartments after the co-treatment of rapamycin and myelin debris (**Fig. 4D**), suggesting engagement of a LC3-dependent degradative pathway distinct from canonical autophagy.

**Figure 4.**
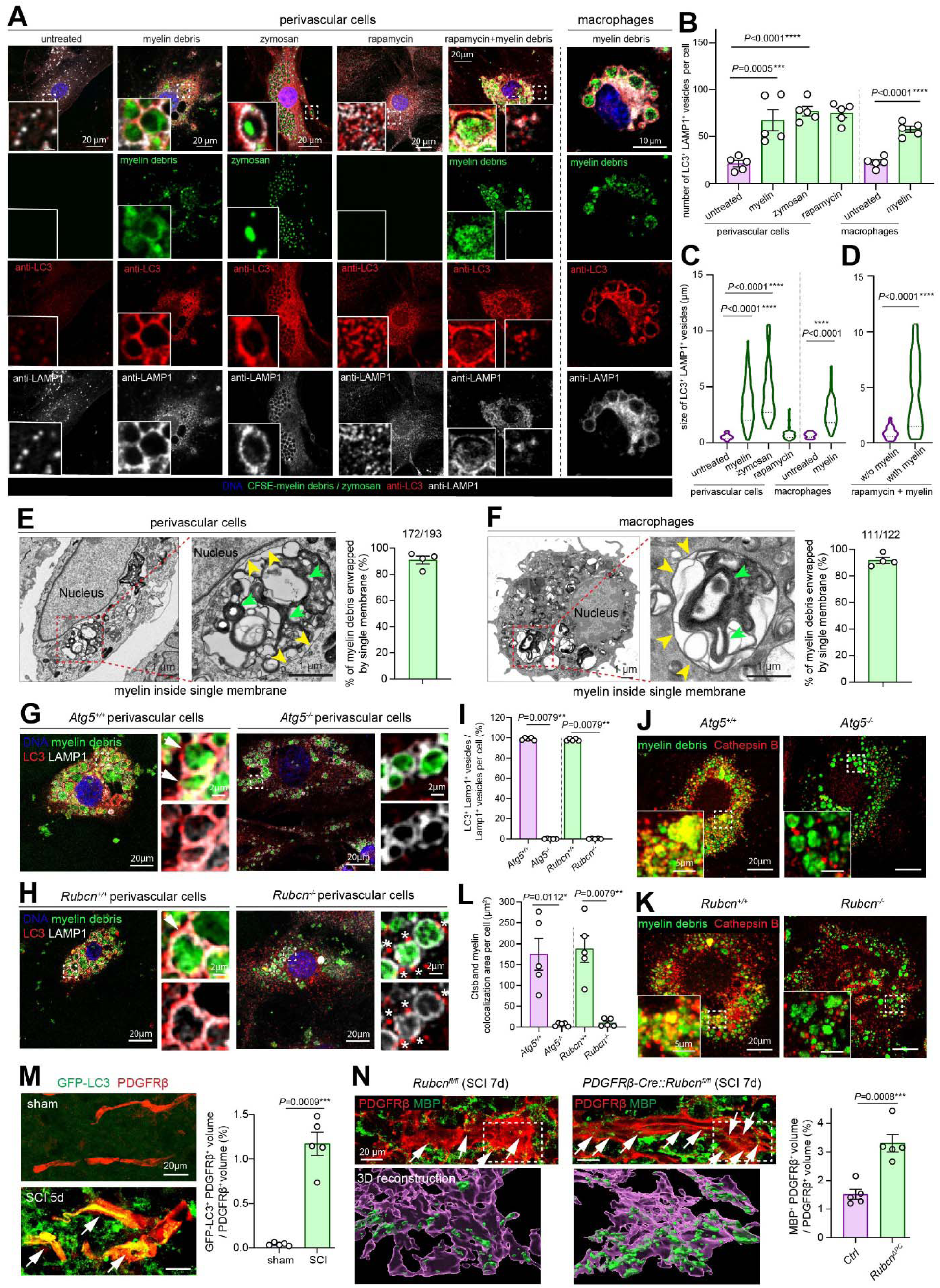
Non-canonical degradative pathway facilitates degradation of engulfed cargo by perivascular cells. **(A)** Representative confocal images showing LC3 and LAMP1 immunostaining in primary wild-type perivascular cells and bone marrow-derived macrophages following treatment with CFSE-labeled myelin debris, CFSE-labeled zymosan, or rapamycin (200 nM, 24 hr). Insets show representative LC3 and LAMP1 staining. Rapamycin induced small LC3^+^ LAMP1^+^ vesicles characteristic of canonical autophagy, whereas myelin debris and zymosan induced large LC3^+^ LAMP1^+^ vesicles surrounding engulfed cargo. Co-treatment with rapamycin and myelin debris showed that intracellular myelin debris was predominantly associated with large LC3^+^ LAMP1^+^ vesicles. Scale bar, 20 μm (perivascular cells), 2 μm (insets), and 10 μm (macrophages). **(B)** Quantification showing the number of LC3^+^ LAMP1^+^ vesicles per perivascular cell or bone marrow-derived macrophage under the indicated treatments. Data are presented as mean ± SEM (*n* = 5 biological replicates). Statistical significance was determined by one-way Welch ANOVA with Dunnett’s post hoc test. **(C)** Quantification showing the size of LC3^+^ LAMP1^+^ vesicles under the indicated treatments. Data are presented as mean ± SEM (*n* = 5 biological replicates). Statistical significance was determined by the Kruskal–Wallis test with Dunn’s post hoc test. **(D)** Quantification comparing the size of LC3^+^ LAMP1^+^ vesicles associated with intracellular myelin debris or without (w/o) myelin association in perivascular cells co-treated with rapamycin and myelin debris. Vesicles associated with myelin debris were significantly larger than those without myelin association. Data are presented as mean ± SEM (*n* = 5 biological replicates). Statistical significance was determined by the Mann–Whitney test. **(E, F)** Representative transmission electron micrographs showing membrane ultrastructure in myelin-laden perivascular cells **(E)** and bone marrow-derived macrophages **(F)**. Myelin debris, identified by its characteristic multilamellar membrane structure (green arrowheads), is enclosed by single-membrane vesicles (yellow arrowheads). Additional single-membrane vesicles containing degrading myelin are also visible. Scale bar, 1 μm. Quantification shows the percentage of intracellular myelin debris enclosed by single-membrane vesicles. Data are presented as mean ± SEM (172 of 193 myelin particles in **E**, and 111 of 122 myelin particles in **F**; *n* = 4 biological replicates). **(G, H)** Representative confocal images showing LC3 (red) and LAMP1 (white) immunostaining in *Atg5^+/+^*, *Atg5^−/−^*, *Rubcn^+/+^* and *Rubcn^−/−^* primary perivascular cells following treatment with CFSE-labeled myelin debris (green). Control cells exhibited LC3 recruitment to myelin-containing LAMP1-positive vesicles (arrows), whereas *Atg5^−/−^*cells or *Rubcn^−/−^* cells lacked LC3 recruitment despite retaining LAMP1^+^ vesicles. Scale bar, 20 μm and 2 μm (insets). **(I)** Quantification showing the percentage of LC3^+^ LAMP1^+^ vesicles relative to total LAMP1^+^ vesicles in the indicated perivascular cells. Data are presented as mean ± SEM (*n* = 5 biological replicates). Statistical significance was determined by the Mann–Whitney test. **(J, K)** Representative confocal images showing Cathepsin B (Ctsb; red) immunostaining in *Atg5^+/+^*, *Atg5^−/−^*, *Rubcn^+/+^* and *Rubcn^−/−^* primary perivascular cells following treatment with CFSE-labeled myelin debris (green) for 24 hr. Scale bar, 20 μm and 5 μm (enlarged views). **(L)** Quantification showing the colocalization area between Cathepsin B and intracellular myelin debris in the indicated perivascular cells. Data are presented as mean ± SEM (*n* = 5 biological replicates). Statistical significance was determined by an unpaired two-tailed *t* test with Welch’s correction (Atg5) or the Mann–Whitney test (Rubcn). **(M)** Representative confocal images showing GFP-LC3 localization in PDGFRβ-positive perivascular cells from sham-operated mice and SCI mice 5 days post injury. Perivascular cells displayed diffuse GFP-LC3 fluorescence in sham spinal cords but prominent GFP-LC3 puncta (arrows) following SCI. Scale bar, 20 μm. Quantification shows the percentage of GFP-LC3 volume relative to total PDGFRβ-positive cell volume. Data are presented as mean ± SEM (*n* = 5 mice). Statistical significance was determined by an unpaired two-tailed *t* test with Welch’s correction. **(N)** Representative confocal images (top) and 3D reconstructions (bottom) showing intracellular myelin debris (MBP, green) in PDGFRβ-positive perivascular cells from control and perivascular cell-specific *Rubcn* knockout (*Rubcn*^Δ*PC*^) mice 7 days after SCI. *Rubcn*^Δ*PC*^ perivascular cells accumulated more intracellular MBP-positive myelin particles (arrows) than control cells. Scale bar, 20 μm. Quantification shows the relative intracellular myelin volume (MBP-positive volume normalized to total PDGFRβ-positive cell volume). Data are presented as mean ± SEM (*n* = 5 mice). Statistical significance was determined by an unpaired two-tailed *t* test.

To define the nature of these structures, we inspected myelin-laden perivascular cells and macrophages by transmission electron microscopy (TEM). In both cell types, engulfed myelin was enclosed by single-membrane compartments (**Fig. 4E, F**), unlike the double-membraned autophagosomes reported during Schwann cell-mediated myelin clearance and characteristic of canonical autophagy ^54^. Instead, together with the LC3 recruitment observed above, the ultrastructural features closely resemble LC3-associated phagocytosis (LAP), a specific degradative pathway in which LC3 is recruited to single-membrane phagosomes to promote cargo degradation ^56^.

To directly test whether LAP mediates myelin degradation, we first deleted *Atg5* in perivascular cells, a core component required for LC3 lipidation during both canonical autophagy and LAP ^56^. To circumvent the developmental lethality associated with germline *Atg5* deletion ^57^, we generated inducible *Atg5*-deficient perivascular cells using *CAG-CreER::Atg5*^fl/fl^ mice (**Fig. S15A–C**). 4-hydroxy tamoxifen-mediated *Atg5* deletion in primary perivascular cells abolished LC3 recruitment to LAMP1⁺ cargo-containing compartments (**Fig. 4G, I**), indicating defective formation of LAP-associated phagosomes, or LAPosomes. In line with this, Ctsb localization to myelin-containing vesicles was significantly reduced (**Fig. 4J–L**), consistent with impaired lysosomal processing. Functional analysis further demonstrated that *Atg5*-deficient perivascular cells failed to efficiently degrade engulfed myelin, resulting in reduced amounts of neutral lipid degradation products, and accumulation of undegraded myelin material (**Fig. S15G–K**). These findings established a critical role for LC3-dependent machinery in cargo degradation.

A defining molecular feature that distinguishes LAP from canonical autophagy is its dependence on Rubicon, which is essential for LAP but dispensable for—and inhibitory to—canonical autophagy ^58^. We therefore generated global *Rubcn* knockout mice and isolated primary *Rubcn*-deficient perivascular cells for mechanistic analysis (**Fig. S15D–F**). Similar to *Atg5* deficiency, *Rubcn* loss abolished LAPosome formation around engulfed myelin, reduced Ctsb recruitment to cargo-containing compartments, and impaired myelin degradation (**Fig. 4H–L**; **Fig. S15I–K**). These results identify Rubicon-dependent LAP, rather than canonical autophagy, as the principal pathway mediating myelin degradation in perivascular cells.

We next investigated whether LAP operates *in vivo* after SCI. In GFP-LC3 reporter mice, LC3 puncta were scarce in perivascular cells under normal conditions but became significantly induced following SCI (**Fig. 4M**). To determine whether LAP is required for myelin degradation *in vivo*, we generated perivascular cell-specific *Rubcn* knockout (*Rubcn*^Δ*PC*^) mice by crossing a newly generated *Rubcn*^fl/fl^ allele with PDGFRβ-Cre mice (**Fig. S15L**). Following SCI, *Rubcn*-deficient perivascular cells accumulated substantially more intracellular myelin debris than their control counterparts (**Fig. 4N**). Together with the *in vitro Rubcn* KO data, these findings indicate that cargo degradation in perivascular cells requires LAP.

Collectively, these findings demonstrate that injury-associated perivascular cells employ a Rubicon-dependent LAP pathway to process engulfed cargo. Thus, injury-associated perivascular cells engage a non-canonical LC3-associated degradative mechanism classically associated with professional phagocytes, enabling efficient phagocytosis of myelin debris following CNS injury.

### Axl receptor mediates perivascular cell phagocytosis of myelin debris

To identify the receptor responsible for myelin debris recognition, we first tested whether perivascular cells utilize phagocytic pathways, including LRP1, Fc receptor and complement receptor, which have been previously implicated in myelin clearance by macrophages, microglia, astrocytes, or oligodendrocytes ^59–61^. Genetic disruption of LRP1 did not affect myelin uptake by perivascular cells (**Fig. S16A–E**). Likewise, manipulation of immunoglobulin-dependent opsonization or deletion of the high-affinity Fcγ receptor Fcgr1 failed to alter myelin uptake (**Fig. S16F–J**). In addition, loss of the complement receptor CR3 had no detectable effect on myelin uptake (**Fig. S16K, L**). Thus, perivascular cells do not rely on several established myelin recognition pathways that mediate debris clearance by other cell types.

To identify the putative receptor, we performed an unbiased survey of phagocytic receptor expression by reanalyzing multiple published single-cell RNA sequencing datasets from mouse and human CNS vasculature and spinal cord injury ^13,24,62–64^. Most canonical phagocytic receptors that are highly expressed by macrophages, microglia, or astrocytes, including *Trem2*, *Megf10*, and *Cd36* were absent or expressed at barely undetectable levels in perivascular cells (**Fig. S17A**). In contrast, Axl emerged as the only well-established phagocytic receptor consistently enriched in perivascular cell populations across both mouse and human CNS datasets (**Fig. S17B–D**). Axl belongs to the TAM receptor family, together with MerTK and Tyro3, which are well known for mediating phosphatidylserine (PtdSer)-dependent phagocytosis of apoptotic cells by professional phagocytes ^65^. However, unlike Axl, Tyro3 was barely detectable in perivascular cells and Mertk was predominantly expressed by microglia (**Fig. S17B**). Under normal conditions, *Axl* expression was highly restricted to perivascular cells, with little or no expression detected in microglia, astrocytes, neurons, or other neural cell types (**Fig. S17B–E**). Following SCI, perivascular cells remained the predominant *Axl*-expressing population throughout the injury response, whereas infiltrating myeloid cells only gradually upregulated *Axl* (**Fig. S17E**). Thus, re-analysis of published data point to Axl as a plausible candidate receptor initiating uptake and phagocytosis in perivascular cells.

We next validated the cellular expression pattern of Axl *in vivo*. Hybridization chain reaction-based fluorescence in situ hybridization (HCR-FISH) demonstrated that *Axl* transcripts were predominantly localized to spinal cord perivascular cells, with substantially lower expression in other CNS cell types (**Fig. S17F, F’**). Consistent with these observations, immunostaining revealed robust Axl protein expression in PDGFRβ^+^ perivascular cells, weaker expression in astrocytes, and little or no detectable expression in neurons, microglia, or oligodendrocytes (**Fig. 5A–C**). Notably, Axl expression remained high in perivascular cells before and after SCI (**Fig. S17G–H’**). These findings indicate that perivascular cells constitutively express Axl and likely utilize a pre-existing phagocytic receptor program, rather than inducing Axl expression *de novo* in response to injury.

**Figure 5.**
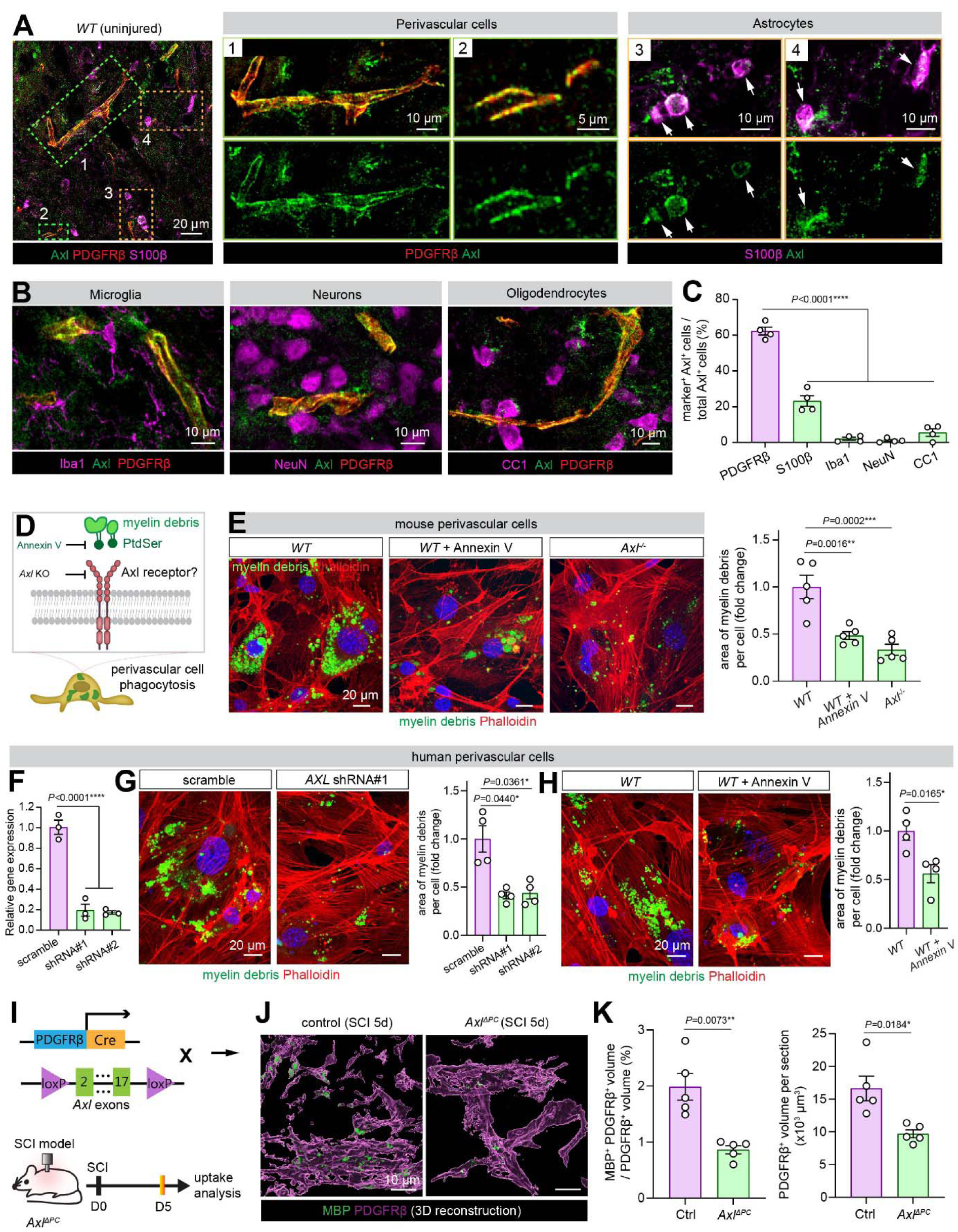
Axl is the receptor for myelin debris phagocytosis by perivascular cells. **(A)** Representative confocal images showing Axl protein (green) localization in PDGFRβ^+^ perivascular cells (red) and S100β^+^ astrocytes (magenta) in the spinal cords of uninjured mice. Boxed regions (#1–4) are shown at higher magnification on the right. Note the preferential localization of Axl in perivascular cells, with weak localization in astrocytes. Scale bar: 20 μm (left), 10 μm (enlarged views #1, #3, and #4), 5 μm (enlarged view #2). **(B)** Representative confocal images showing prominent Axl protein expression (green) in PDGFRβ^+^ perivascular cells (red), but little or no expression in Iba1^+^ microglia, NeuN^+^ neurons, or CC1^+^ oligodendrocytes (magenta). Scale bar: 10 μm. **(C)** Quantification showing the percentage of Axl^+^ cells co-expressing the indicated cell-type markers. Data are shown as mean ± SEM (n=4 mice). Statistical significance was determined by one-way ANOVA with Dunnett’s post hoc test. **(D)** Schematic illustrating the proposed model in which Axl acts as the phagocytic receptor for perivascular cells by recognizing phosphatidylserine (PtdSer) exposed on myelin debris (bridging ligands not shown), together with the experimental strategies used to test this model, including masking PtdSer with recombinant Annexin V and genetic deletion of Axl. **(E)** Representative confocal images showing reduced uptake of myelin debris by primary mouse brain perivascular cells following PtdSer masking with recombinant Annexin V (4 μg) or genetic deletion of Axl. Quantification shows the relative area of engulfed myelin debris normalized to untreated wild-type controls. Data are shown as mean ± SEM (n=5 biological replicates). Statistical significance was determined by one-way ANOVA with Dunnett’s post hoc test. **(F)** qRT-PCR validation of *AXL* knockdown in primary human brain perivascular cells using two independent shRNAs. Data are shown as mean ± SEM (n=3 biological replicates). Statistical significance was determined by one-way ANOVA with Dunnett’s post hoc test. **(G)** Representative confocal images showing reduced uptake of myelin debris (green) by primary human brain perivascular cells (red) following shRNA-mediated *AXL* knockdown. Scale bar: 20 μm. Quantification shows the relative area of engulfed myelin debris normalized to scramble shRNA controls. Data are shown as mean ± SEM (n=4 biological replicates). Statistical significance was determined by one-way ANOVA with Dunnett’s post hoc test. **(H)** Representative confocal images showing reduced uptake of myelin debris by primary human brain perivascular cells following PtdSer masking with recombinant Annexin V (4 μg). Scale bar: 20 μm. Quantification shows the relative area of engulfed myelin debris normalized to untreated controls. Data are shown as mean ± SEM (n=4 biological replicates). Statistical significance was determined by unpaired two-tailed *t* test. **(I)** Schematic illustrating the generation of perivascular cell-specific *Axl* knockout mice (*Axl*^Δ*PC*^) using the Cre-loxP system. A newly generated Axl floxed allele, in which exons 2–17 are flanked by loxP sites, was crossed with the PDGFRβ-Cre line. Control and *Axl*^Δ*PC*^ mice were subjected to SCI and analyzed 5 days post injury for myelin debris uptake by perivascular cells. **(J)** Representative 3D reconstruction images showing reduced uptake of myelin debris (green) by PDGFRβ^⁺^ perivascular cells (magenta) in *Axl*^Δ*PC*^ mice compared with control mice 5 days after SCI. Scale bar: 10 μm. **(K)** Quantification showing the relative volume of engulfed myelin debris within PDGFRβ^+^ perivascular cells and the total PDGFRβ^⁺^ perivascular cell volume in control and *Axl*^Δ*PC*^ mice following SCI. Data are shown as mean ± SEM (n=5 mice). Statistical significance was determined by unpaired two-tailed *t* test with Welch’s correction.

To determine whether Axl is functionally required for myelin debris phagocytosis, we isolated perivascular cells from global *Axl* knockout mice and assessed cargo uptake *in vitro* (**Fig. 5D**). Loss of *Axl* markedly impaired engulfment of myelin debris (**Fig. 5E**). Axl mediates phagocytosis of phosphatidylserine (PtdSer)-exposed targets through bridging ligands such as Gas6 or Protein S (**Fig. 5D**), a mechanism well characterized in apoptotic cells ^66^. Given that myelin is highly rich in lipids (70-85%), including PtdSer ^67^, we hypothesized that PtdSer-dependent TAM receptor signaling contributes to perivascular cell-mediated myelin phagocytosis. Consistent with this idea, masking PtdSer with Annexin V significantly reduced myelin uptake by mouse perivascular cells (**Fig. 5E**), phenocopying the *Axl* knockout. Similar reduction of myelin debris engulfment was observed in primary human perivascular cells following lentiviral knockdown of human *AXL* or blockade of PtdSer recognition (**Fig. 5F–H**). These findings identify Axl as a conserved receptor mediating PtdSer-dependent myelin debris recognition and uptake in both mouse and human perivascular cells *in vitro*.

We next examined the requirements for Axl *in vivo*. Given that Axl transcripts and proteins were also detected in other cell types, albeit at a much lower level (**Fig. 5A-C, Fig. S17E, E’**), we generated perivascular cell-specific *Axl* knockout (*Axl*^Δ*PC*^) mice by crossing PDGFRβ-Cre mice with an *Axl* floxed strain (**Fig. 5I, Fig. S18A, B**). In *Axl*^Δ*PC*^ animals, Axl protein was significantly reduced in PDGFRβ⁺ perivascular cells but remained unchanged in PDGFRβ negative cells (**Fig. S18C**). Following SCI, *Axl*-deficient perivascular cells significantly reduced myelin debris uptake compared with cells from control littermates (**Fig. 5J, K**), confirming that Axl is required for myelin uptake by perivascular cells *in vivo*.

Pericytes, a major population of CNS perivascular cells are critical for endothelial transcytosis and blood-brain barrier (BBB) integrity ^28,30,68^. To exclude possible confounding effects of perivascular cell loss of Axl on vascular physiology, we evaluated BBB integrity in *Axl*^Δ*PC*^ mice by three complementary BBB assays, including permeability to the very small tracer Sulfo-NHS-Biotin (443Da), distribution of tight junction marker ZO-1 and transcytosis marker Mfsd2a. *Axl*^Δ*PC*^ mice showed neither leakage of Sulfo-NHS-Biotin, nor did they exhibit signs of defective tight junctions or increased transcytosis (**Fig. S18D–F**), indicating that Axl depletion in perivascular cells does not cause baseline vascular abnormalities. Therefore, the reduced myelin uptake after SCI is unlikely to be an indirect consequence of baseline vascular abnormalities following *Axl* deletion.

Given that perivascular cells also engulf NeuN⁺ neuronal material (**Fig. S11**), we next asked whether this process similarly depends on Axl. Interestingly, unlike the Axl-dependent phagocytosis of myelin debris, Axl deficiency did not affect the engulfment of NeuN⁺ neuronal material by perivascular cells following MCAO (**Fig. S19A–C**). Consistently, *in vitro* phagocytosis assays showed that neither Axl depletion nor PtdSer masking with Annexin V altered neuronal debris uptake by cultured perivascular cells (**Fig. S19D, E**). These findings indicate that, unlike myelin debris uptake, perivascular cell phagocytosis of neuronal material occurs independently of Axl and PtdSer recognition, suggesting the involvement of distinct, yet-to-be-identified phagocytic receptor(s).

Together, these *in vivo* and *in vitro* data demonstrate that Axl is selectively required for perivascular cell phagocytosis of myelin debris, consistent with a critical role for PtdSer as an ‘eat-me’ signal in this context.

### Perivascular cell phagocytosis of myelin debris drives lesion expansion after SCI

Prior studies suggested that while timely and efficient clearance of myelin debris by macrophages and glial cells is beneficial, inefficient degradation of engulfed myelin debris results in persistent intracellular accumulation of myelin-derived lipids, particularly cholesterol, driving phagocytes towards disease-promoting states that impairs tissue repair ^31–33,45,69^. To determine the pathophysiological consequences of Axl-mediated perivascular cell phagocytosis of myelin debris, we subjected *Axl*^Δ*PC*^ mice to SCI and evaluated perivascular cell proliferation and related pathological changes by tissue histology and behavioral analysis (**Fig. 6A**). In control mice subjected to SCI, perivascular cells in the injured spinal cord exhibited high proliferative activity, as demonstrated by immunostaining for cell proliferation marker Ki67. By contrast, *Axl*^Δ*PC*^ mice showed a significant reduction in both proliferating and total perivascular cells following SCI (**Fig. 6B–D**), suggesting that Axl-dependent phagocytosis of myelin debris drives perivascular cell proliferation following SCI.

**Figure 6.**
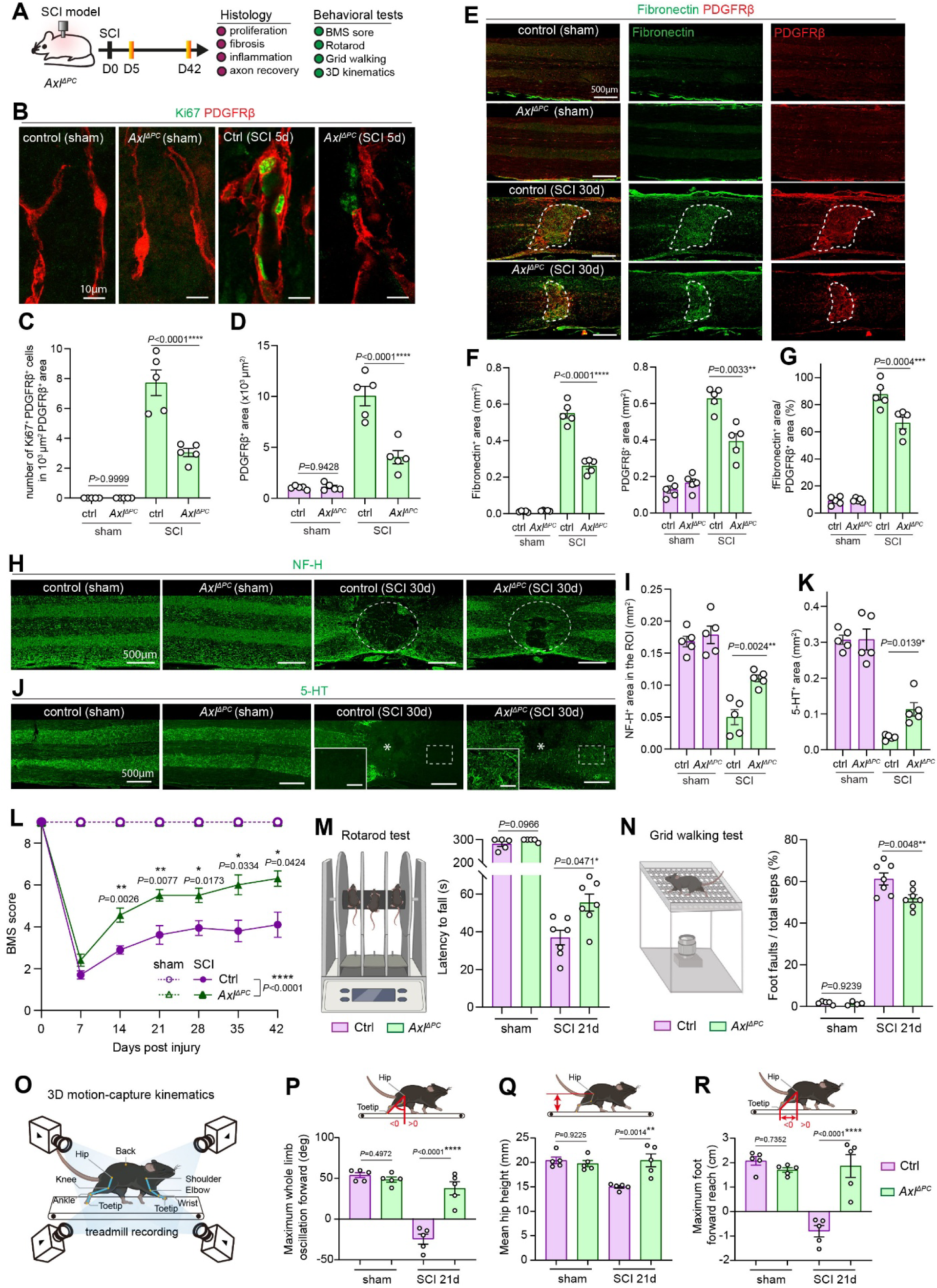
Genetic blockade of Axl-dependent phagocytosis in perivascular cells ameliorates CNS injury pathology. **(A)** Schematic illustrating the experimental design for histological and behavioral analyses of control and *Axl*^Δ*PC*^ mice following SCI. **(B)** Representative confocal images showing proliferation marker Ki67 (green) in PDGFRβ^+^ perivascular cells (red) in spinal cords from control and *Axl*^Δ*PC*^ mice 5 days after sham operation or SCI. Scale bar: 10 μm. **(C, D)** Quantification of Ki67^+^ PDGFRβ^+^ cells normalized to PDGFRβ^+^ area **(C)** and PDGFRβ^+^ cell area **(D)** in spinal cords from the indicated groups. Data are shown as mean ± SEM (n=5 mice). Statistical significance was determined by two-way ANOVA with Holm–Šídák’s post hoc test. **(E)** Representative confocal images showing fibronectin (green) and PDGFRβ^+^ perivascular cells (red) in spinal cords from control and *Axl*^Δ*PC*^ mice 30 days after SCI. Dashed outlines indicate the lesion area. Scale bar: 500 μm. **(F, G)** Quantification of fibronectin^+^ area and PDGFRβ^+^ area **(F),** and fibronectin^+^ area normalized to PDGFRβ^+^ area **(G).** Data are shown as mean ± SEM (n=5 mice). Statistical significance was determined by two-way ANOVA with Holm–Šídák’s post hoc test. **(H, I)** Representative confocal images **(H)** and quantification **(I)** of NF-H^+^ axons (green) in spinal cords from control and *Axl*^Δ*PC*^ mice 30 days after sham operation or SCI. Dashed outlines indicate the region used for quantification. Scale bar: 500 μm. Data are shown as mean ± SEM (n=5 mice). Statistical significance was determined by two-way ANOVA with Holm–Šídák’s post hoc test. **(J, K)** Representative confocal images **(J)** and quantification **(K)** of 5-HT^+^ axon fibers (green) in spinal cords from control and *Axl*^Δ*PC*^ mice 30 days after SCI. Insets show higher-magnification views of the caudal spinal cord. Dashed outlines indicate the region used for quantification. Scale bar: 500 μm; 100 μm (insets). Data are shown as mean ± SEM (n=5 mice). Statistical significance was determined by two-way ANOVA with Holm–Šídák’s post hoc test. **(L)** Basso Mouse Scale (BMS) scores over 42 days following sham operation or SCI in control and *Axl*^Δ*PC*^ mice. Data are shown as mean ± SEM (n=5–10 mice). Statistical significance was determined by two-way ANOVA (mixed-effects model) with Geisser–Greenhouse correction followed by Dunnett’s post hoc test. **(M)** Rotarod performance, measured as latency to fall, in the indicated groups. Data are shown as mean ± SEM (n=5–7 mice). Statistical significance was determined by two-way ANOVA with Holm–Šídák’s post hoc test. **(N)** Grid walking performance, quantified as the percentage of faulty foot placements, in the indicated groups. Data are shown as mean ± SEM (n=4–7 mice). Statistical significance was determined by two-way ANOVA with Holm–Šídák’s post hoc test. **(O)** Schematic of the markerless 3D motion-capture system used for hindlimb kinematic analysis. **(P–R)** Schematic and quantifications of hindlimb kinematic parameters, including maximum whole-limb forward oscillation **(P),** mean hip height **(Q),** and maximum forward foot reach **(R),** in control and *Axl*^Δ*PC*^ mice 21 days after sham operation or SCI. Data are shown as mean ± SEM (n=5 mice). Statistical significance was determined by two-way ANOVA with Holm–Šídák’s post hoc test.

To directly assess whether Axl-dependent engulfment of myelin debris promotes perivascular cell proliferation, we performed three complementary assays. First, we measured cell proliferation in cultured perivascular cells, and observed that myelin debris markedly increased proliferation in wildtype cells, whereas *Axl* deficiency significantly reduced this response (**Fig. S20A–B’**). Second, we transplanted primary perivascular cells that had been preloaded with or without myelin debris into Matrigel plugs and quantified their proliferative activity *in vivo* (**Fig. S20C**). Whereas untreated cells exhibited only basal proliferation, myelin debris-laden cells showed robust expansion within the plugs, accompanied by enlarged morphology and increased proliferative activity (**Fig. S20D, D’**). These effects were markedly attenuated in Matrigel plugs of Axl-deficient perivascular cells (**Fig. S20D, D’**). Third, we directly microinjected myelin debris into uninjured spinal cords of control and *Axl*^Δ*PC*^ mice to examine the local perivascular cell response (**Fig. S20E**). Compared to PBS injection, injected myelin debris markedly increased perivascular cell proliferation and PDGFRβ⁺ area. Both effects were substantially reduced in *Axl*^Δ*PC*^ mice (**Fig. S20F, F’**).

Together, these results establish Axl-mediated uptake of myelin debris as a key driver of pathological perivascular cell proliferation after SCI.

### Perivascular cell phagocytosis of myelin debris contributes to CNS fibrosis

Fibrosis is a defining feature of CNS injury that hinders axonal regeneration through excessive extracellular matrix (ECM) deposition ^19,70^. Because perivascular cells are a major source of fibrotic scar tissue^17–22^ and Axl-mediated myelin uptake promotes their proliferation, we hypothesized that myelin debris engulfment by perivascular cells promotes CNS fibrosis. Indeed, *Axl*^Δ*PC*^ mice exhibited markedly reduced fibrotic scar formation after SCI, with significantly decreased deposition of fibronectin and collagen I and a smaller perivascular cell-covered lesion area compared to control mice (**Fig. 6E–G, Fig. S21A, B**).

Because TGF-β signaling is a master regulator of fibrosis ^71^, we asked whether Axl-dependent myelin debris engulfment activates this pathway in perivascular cells. Following SCI, perivascular cells displayed a significant increase in nuclear phosphorylated Smad3 (p-Smad3), indicative of TGF-β pathway activation, whereas this response was significantly reduced in *Axl*^Δ*PC*^ mice (**Fig. S21C–E**). Strikingly, myelin debris alone induced robust p-Smad3 nuclear translocation in cultured perivascular cells, reaching levels comparable to recombinant TGF-β1 stimulation, and this effect was significantly reduced by *Axl* deficiency (**Fig. S21F, G**). These findings identify myelin debris as a previously unrecognized trigger of TGF-β signaling in perivascular cells, providing a mechanistic link between Axl-mediated myelin phagocytosis and fibrotic scar formation after SCI.

To test whether myelin debris directly stimulates ECM production, we measured fibronectin and collagen I expression by cultured perivascular cells following myelin uptake. Whereas untreated cells expressed little fibronectin or collagen I, myelin debris robustly induced expression of both proteins to levels comparable to recombinant TGF-β1 stimulation (**Fig. S21H–K**). Notably, this induction was significantly blunted in *Axl*-deficient cells (**Fig. S21H–K**), establishing Axl-dependent myelin debris engulfment as a key driver of ECM protein expression. Collectively, these findings suggest that myelin debris activates Axl-dependent TGF-β signaling, leading to increased ECM protein expression in perivascular cells.

### Genetic blockade of Axl-dependent phagocytosis in perivascular cells ameliorates injury pathology and improves functional recovery after SCI

Fibrotic scar formation stabilizes injured CNS tissue but ultimately hampers repair by generating a dense ECM barrier that restricts axonal regeneration and functional recovery ^70^. Indeed, genetic ablation of proliferating perivascular cells reduces fibrosis and improves functional recovery after SCI ^19^. Because Axl-dependent phagocytosis promoted perivascular cell expansion and fibrosis (**Fig. 6B–G**), lesion progression and macrophage infiltration (**Fig. S22A–C**), we investigated whether targeted inhibition of this pathway enhances neural repair and functional recovery after SCI.

To assess neuroaxonal repair after SCI, we first quantified neurofilament heavy chain-positive (NF-H⁺) axons 30 days after injury. Whereas NF-H⁺ axons were largely absent from the lesion region of control mice following SCI, *Axl*^Δ*PC*^ mice exhibited a significant increase in axonal density within the injured region (**Fig. 6H, I**). We next examined serotonergic (5-HT⁺) fibers of the raphe-spinal tract, which play essential roles in sensory and motor function ^72^. 5-HT⁺ fibers were nearly completely lost in control mice at 30 days post-injury (**Fig. 6J, K**). In contrast, *Axl*^Δ*PC*^ mice displayed substantially more 5-HT⁺ axons within and around the lesion (**Fig. 6J, K**). Together, these findings indicate that inhibition of perivascular cell phagocytosis creates a more permissive environment for axonal preservation and/or regrowth after SCI.

We next assessed whether inhibition of perivascular cell phagocytosis improves functional recovery after SCI. Consistent with the enhanced axonal preservation and/or regrowth observed in *Axl*^Δ*PC*^ mice compared to control animals, these animals exhibited significantly improved locomotor recovery, as measured by the Basso Mouse Scale (BMS) (**Fig. 6L**). Improved motor performance was confirmed by the Rotarod assay, in which *Axl*^Δ*PC*^ mice remained on the rotating rod significantly longer than SCI-treated control mice (**Fig. 6M**; **Fig. S22D**). Likewise, in the grid-walking assay, *Axl*^Δ*PC*^ mice exhibited fewer foot-placement errors than control mice, indicating improved sensorimotor coordination (**Fig. 6N**; **Fig. S22E**). To further quantify locomotor function, we employed a 3D motion-capture system that combines live-animal multi-view imaging with markerless 3D skeleton reconstruction and kinematic analysis ^73^(**Fig. 6O**). Whereas SCI caused severe impairment of hindlimb kinematics in control mice, several parameters related to hindlimb stepping were significantly improved in *Axl*^Δ*PC*^ mice (**Fig. 6P–R**; **Fig. S22F**). Collectively, these findings demonstrate that Axl-dependent phagocytosis by perivascular cells contributes to neurological dysfunction and that its inhibition promotes functional recovery after SCI.

### Gilteritinib, an Axl-targeting drug, selectively inhibits perivascular cell phagocytosis without affecting phagocytosis by other cells

Given the beneficial effects of perivascular cell-specific *Axl* deletion after SCI, we next explored the therapeutic potential of pharmacological approach. We focused on Gilteritinib, an FDA-approved tyrosine kinase inhibitor with potent activity against Axl that is currently used to treat FLT3-mutant acute myeloid leukemia ^74^. Gilteritinib robustly inhibited myelin debris uptake by both mouse and human perivascular cells *in vitro* for 24 hr (**Fig. 7A–C**). Moreover, oral administration of Gilteritinib (5 mg/kg), beginning 1 day before SCI and continuing for 7 days after injury, significantly reduced myelin debris engulfment by perivascular cells *in vivo* (**Fig. 7D, E**). These findings establish perivascular cell phagocytosis as a pharmacologically tractable target and support the translational potential of Axl inhibition for SCI.

**Figure 7.**
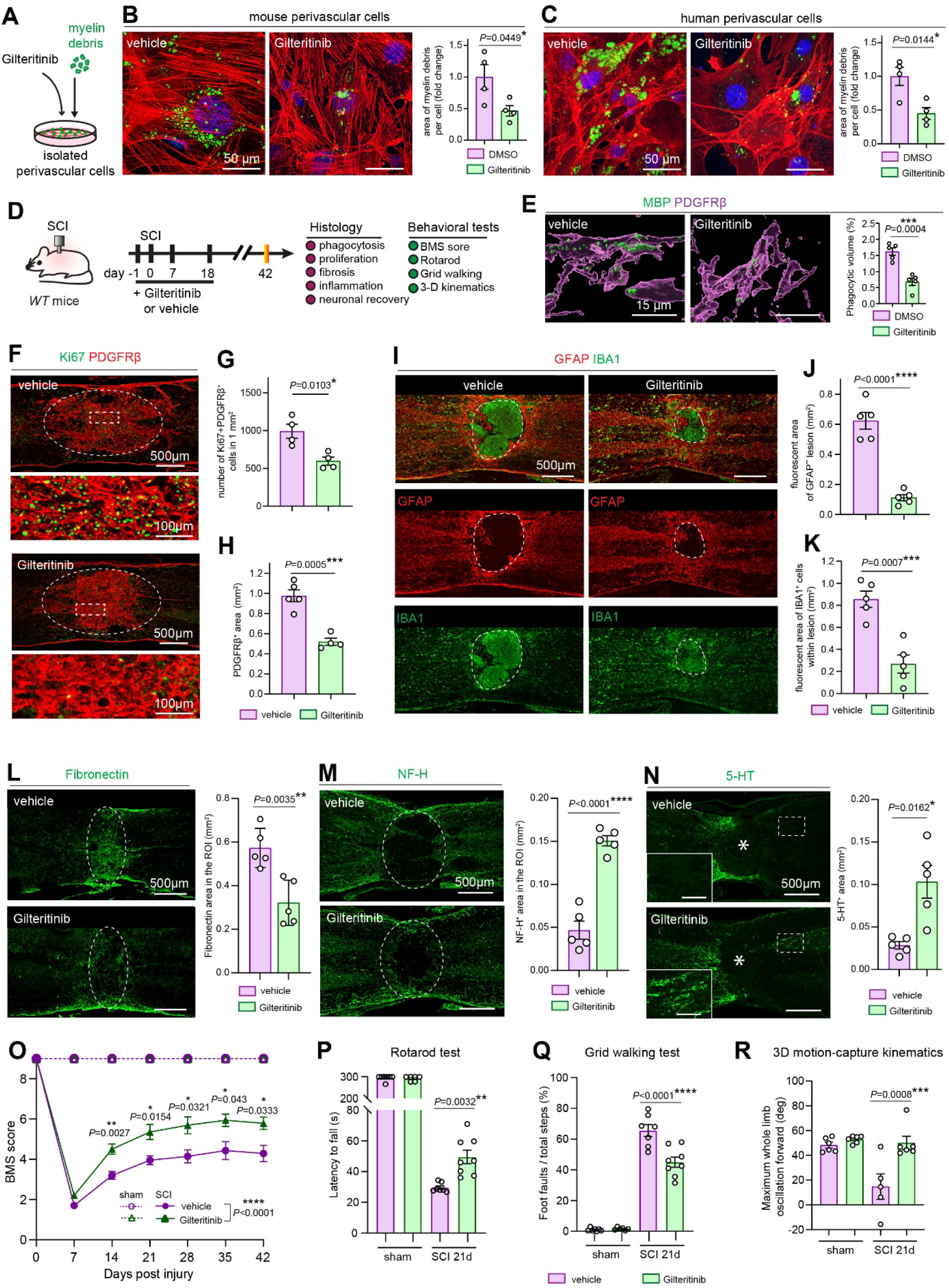
Gilteritinib inhibits Axl-dependent phagocytosis by perivascular cells and ameliorates CNS injury pathology. **(A)** Schematic illustrating the experimental design for evaluating the effect of Gilteritinib on myelin debris phagocytosis by primary brain perivascular cells *in vitro*. **(B, C)** Representative confocal images and Quantification showsing reduced uptake of CFSE-labeled myelin debris (green) by primary mouse **(B)** and human **(C)** brain perivascular cells (α-SMA staining, red) following Gilteritinib treatment (1 μM, 24 hr). Scale bar: 50 μm. Quantification shows the relative area of engulfed myelin debris normalized to DMSO vehicle-treated controls. Data are shown as mean ± SEM (*n*=4 biological replicates). Statistical significance was determined by unpaired two-tailed *t* test. **(D)** Schematic illustrating the experimental design for evaluating the effect of Gilteritinib treatment in wild-type mice following SCI. **(E)** Representative 3D reconstruction images showing reduced uptake of MBP^+^ myelin debris (green) by PDGFRβ^+^ perivascular cells (magenta) in Gilteritinib-treated mice compared with vehicle-treated controls 5 days after SCI. Scale bar: 15 μm. Quantification shows the relative volume of engulfed myelin debris normalized to total PDGFRβ^+^ perivascular cell volume. Data are shown as mean ± SEM (*n*=5 mice). Statistical significance was determined by unpaired two-tailed *t* test. **(F–H)** Representative confocal images **(F)** and quantification **(G, H)** showing reduced proliferation of Ki67^+^ PDGFRβ^+^ perivascular cells in spinal cords from vehicle- and Gilteritinib-treated mice 7 days after SCI. Dashed outlines indicate the regions used for quantification. Scale bar: 500 μm; 100 μm (enlarged view). Quantification shows Ki67^+^ PDGFRβ^+^ cell density **(G)** and total PDGFRβ^+^ area **(H).** Data are shown as mean ± SEM (*n*=4 mice). Statistical significance was determined by unpaired two-tailed *t* test. **(I–K)** Representative confocal images **(I)** and quantifications **(J, K)** showing reduced lesion area and macrophage/microglia accumulation in spinal cords from Gilteritinib-treated mice 42 days after SCI. GFAP-negative lesion area and IBA1^+^ cells are shown. Dashed outlines indicate the regions used for quantification. Scale bar: 500 μm. Data are shown as mean ± SEM (*n*=5 mice). Statistical significance was determined by unpaired two-tailed *t* test. **(L–N)** Representative confocal images and Quantification showsing fibronectin production at 21 days after SCI **(L),** NF-H^+^ neurofilaments at 42 days after SCI **(M),** and 5-HT^+^ axon fibers at 42 days after SCI **(N)** in vehicle- and Gilteritinib-treated mice. Dashed outlines indicate the regions used for quantification. Insets in **(N)** show higher-magnification views of the caudal spinal cord. Scale bar: 500 μm. Quantification shows the area occupied by the indicated markers. Data are shown as mean ± SEM (*n*=5 mice). Statistical significance was determined by unpaired two-tailed *t* test, except for **(N)** by unpaired two-tailed *t* test with Welch’s correction. **(O)** Basso Mouse Scale (BMS) scores over 42 days following sham operation or SCI in vehicle- and Gilteritinib-treated mice. Data are shown as mean ± SEM (n=6–10 mice). Statistical significance was determined by two-way ANOVA (mixed-effects model) with Geisser–Greenhouse correction followed by Dunnett’s post hoc test. **(P–R)** Quantification of motor function 21 days after sham operation or SCI, including Rotarod performance measured as latency to fall **(P),** grid walking performance measured as the percentage of faulty foot placements **(Q),** and maximum whole-limb forward oscillation measured by 3D motion-capture kinematic analysis **(R).** Data are shown as mean ± SEM. Rotarod and grid walking: *n*=6–8 mice. 3D motion-capture kinematic analysis: *n*=5–6 mice. Statistical significance was determined by two-way ANOVA with Holm–Šídák’s post hoc test.

Although Axl expression was enriched in perivascular cells compared to other cell types (**Fig. 5A–C**), we could not exclude that Axl inhibition affects phagocytotic activity in other CNS cell types. Neither Gilteritinib treatment nor genetic Axl deletion impaired myelin debris engulfment by primary astrocytes or bone marrow-derived macrophages *in vitro* (**Fig. S23A–H**) or macrophage phagocytosis *in vivo* after SCI (**Fig. S24A, B, E–F’**). In fact, astrocytes contributed negligibly to debris clearance within the lesion core because they were largely absent from this region after injury (**Fig. S24C–D’**). Thus, pharmacologic or genetic Axl inhibition selectively suppresses myelin debris uptake by perivascular cells while sparing astrocytes and macrophages from inhibition.

### Gilteritinib ameliorates injury pathology and improves functional recovery after SCI

To evaluate the therapeutic efficacy of pharmacological Axl inhibition on functional recovery after SCI, mice received oral Gilteritinib (5 mg/kg) beginning 1 day before injury and continuing through 18 days post-injury, followed by histological and behavioral analyses at the indicated time points (**Fig. 7D**). Consistent with the effects of genetic *Axl* deletion, Gilteritinib significantly reduced perivascular cell proliferation and accumulation within the lesion (**Fig. 7F–H**). This was accompanied by attenuation of the injury response, including reduced size of the GFAP-negative lesion and diminished infiltration of the IBA-1 postive myeloid cells into the lesion core (**Fig. 7I–K**). Transcriptomic profiling further revealed suppression of phagocytic and NF-κB signaling pathways in cultured perivascular cells exposed to myelin debris with Gilteritinib co-treatment (**Fig. S25A–C**).

In parallel, Gilteritinib substantially reduced fibrotic scar formation after SCI (**Fig. 7L**) and suppressed ECM-receptor interaction and TGF-β signaling pathways in perivascular cells (**Fig. S25D, E**). Consistent with these findings, Gilteritinib-treated mice had significantly increased NF-H⁺ axonal and 5-HT⁺ areas at 42 days post-injury (**Fig. 7M, N**).

These anatomical improvements translated into functional recovery. Gilteritinib-treated mice displayed significantly higher BMS scores throughout the recovery period (**Fig. 7O**), accompanied by improved motor coordination, sensorimotor performance and hindlimb kinematics in Rotarod, grid walking assays and 3D motion-capture kinematics analysis (**Fig. 7P–R**; **Fig. S25F–J**). Together, these findings demonstrate that pharmacological inhibition of Axl phenocopies genetic blockade of perivascular cell phagocytosis, remodeling the lesion environment to favor axonal preservation and functional recovery after SCI. Given its clinical approval and robust efficacy *in vivo*, Gilteritinib provides a direct translational path toward targeting detrimental perivascular cell responses after CNS injury

## Discussion

This study identifies perivascular cells as a previously unrecognized phagocytic cell population that emerges across multiple CNS injuries and disease conditions, including SCI, ischemic stroke and EAE. While microglia and infiltrating macrophages have long been considered the principal phagocytes under pathogenic conditions, we show that perivascular cells rapidly expand and engage in robust phagocytosis across multiple demyelinating conditions (**Fig. S26**). Similar responses observed in human stroke lesions further suggest that this represents a conserved injury-associated program. Mechanistically, disease-associated perivascular cells respond to myelin debris through the constitutively expressed Axl receptor and mobilize key components of the phagocytic machinery, including actin-dependent engulfment, lysosomal degradation, and LC3-associated phagocytosis (LAP), a non-canonical degradative pathway. Blocking Axl ameliorated the pathophysiological responses to CNS injury and promoted functional recovery. Together, these findings expand the cellular repertoire of phagocytic cells in the CNS and establish perivascular cells as active participants that shape lesion pathology.

The identification of perivascular cells as phagocytes has important implications for understanding debris clearance within CNS lesions. Although microglia, astrocytes, and infiltrating macrophages possess established phagocytic functions, their spatial and temporal distributions can not fully account for debris handling within lesion cores. Reactive glial cells are largely confined to lesion borders ^75^, whereas macrophage infiltration is delayed and their prolonged lipid uptake can induce dysfunctional foamy states ^33,45,76 69^. By contrast, blood vessels remain abundant within lesion cores, and perivascular cells rapidly expand within these regions after injury. Together with recent studies demonstrating phagocytic functions in vascular endothelial cells ^34,77–80^, our findings reveal vascular-associated cells as an underappreciated component of the CNS phagocytic network. Notably, perivascular cells exhibit robust phagocytic capacity and are capable of engulfing diverse cargos, supporting their role as *bona fide* phagocytes during CNS injury.

A central finding of this study is the identification of Axl as a key regulator governing myelin debris recognition by perivascular cells. The selective requirement for Axl in perivascular cell phagocytosis of myelin debris likely reflects fundamental differences in phagocytic receptor utilization among different cell types. Macrophages and astrocytes express diverse repertoires of phagocytic receptors that may compensate for Axl loss ^81,82^, whereas few phagocytic receptors have been identified in perivascular cells. As the first phagocytic receptor defined in this population, Axl is required for myelin debris engulfment by perivascular cells. Consequently, both genetic and pharmacological inhibition of Axl selectively disrupt perivascular cell phagocytosis without affecting myelin debris uptake by astrocytes or macrophages.

Importantly, Axl-dependent phagocytosis is not simply a debris-clearing process. Instead, engulfment of lipid-rich myelin debris promotes perivascular cell proliferation, activates profibrotic programs, and drives lesion expansion. These observations parallel emerging studies showing that excessive lipid accumulation can convert macrophages, microglia, and astrocytes into disease-associated states ^35–37^, suggesting that maladaptive responses to lipid-rich debris may represent a common mechanism contributing to CNS pathology. Importantly, the identification of Axl as a central regulator of perivascular cell phagocytosis provides a rational therapeutic target for modulating perivascular cell responses after CNS injury. Genetic deletion of Axl in perivascular cells or pharmacological inhibition with the FDA-approved drug Gilteritinib attenuated fibrosis, neuroinflammation, and lesion progression while improving functional recovery after SCI. Because Gilteritinib was administered systemically, we cannot exclude additional biological effects on other Axl-expressing cell populations, including activated macrophages and microglia in CNS as well as peripheral macrophages. Although Gilteritinib did not impair macrophage myelin uptake in our experimental systems, its effects on other macrophage functions remain to be determined.

Our results also raise several new questions. Perivascular cells are a heterogeneous population comprising pericytes, fibroblasts, and vascular smooth muscle cells, and the relative contribution of each subtype to phagocytosis remains unclear. In addition, Axl is required for myelin debris engulfment but dispensable for uptake of neuronal corpses, indicating that distinct recognition pathways exist for different cargos. More broadly, given the widespread distribution of perivascular cells throughout vascularized tissues, it will be important to determine whether similar phagocytic functions operate in other organs following injury. Addressing these questions may reveal previously unappreciated roles for perivascular cells in immunity, tissue remodeling, and regenerative responses.

## Methods and Materials

### Mice

All mice were maintained in specific pathogen-free animal facility at Xiamen University School of Life Sciences, Southern Medical University and Florida State University College of Medicine. Mice were housed and bred on a 12/12 hr light/dark cycle, with ad libitum access to food and water. The use of mice in this study was approved by the Institutional Animal Care and Use Committee of Xiamen University, Southern Medical University and Florida State University College of Medicine.

*Rosa26-LSL-tdTomato* Cre reporter line Ai14 (B6. Cg-Gt(ROSA)26Sortm14(CAG-tdTomato)Hze/J) was from Jackson lab (Bar Harbor, ME) (Stock No. 007914) ^83^. *PDGFR*β*-Cre* mice and *GLAST-CreER^T^*^2^ mice were described previously ^41,84^. To label *PDGFR*β*-*expressing perivascular cells, *PDGFR*β*-Cre* mice were crossed with *Ai14* mice to obtain *PDGFR*β*-Cre*::*Ai14* mice. To avoid the unwanted expression of Cre recombinase and unexpected recombination in the *PDGFR*β*-Cre*::*Ai14* mice, we performed excision detection of *loxP-STOP-loxP* in the *PDGFR*β*-Cre/+*::*Ai14/+* mouse tails using primers flanking the two *loxP* sites (see Genomic PCR below). Only the reporter mice without excision bands were used. To label *GLAST-*expressing perivascular cells, *GLAST-Cre^ERT^*^2^ mice were crossed with *Ai14* mice to obtain *GLAST-CreER^T^*^2^::*Ai14* mice, followed by tamoxifen-mediated Cre recombination at 4 weeks of age. Tamoxifen (Aladdin, Cat# T137974-5g) was dissolved in corn oil (Aladdin, Cat#C116023-500mL) at concentration of 20 mg/ml, and was administered (0.1 mg/g body weight per day for 5 days) by oral gavage. *CAG-CreER^TM^* (B6.Cg-Tg(CAG-cre/Esr1*)5Amc/J) mice were from Jackson lab (Stock No. 004682). *GFP-LC3* BAC transgenic mice were previously described ^85^.

*Mbp^shi^* or *Mbp^−/−^* mice (C57BL/6J, C3Fe.SWV-Mbp^shi^/J), also known as Shiverer mice were from Jackson lab (Stock No. 001428). The shiverer allele arose from a spontaneous mutation resulting in a large deletion from intron 1 to exon 6 in the *Mbp* gene and a lack of functional MBP protein ^86^. *CR3*^−/−^ or *Itgam*^−/−^ mice (B6.129S4-*Itgam^tm1Myd^*/J), disrupting gene region encoding the translational initiation codon and 15 amino acids of the signal peptide, were from Jackson lab (Stock No. 003991) ^87^. *Axl*^−/−^mice (*Axl^tm1Grl^*/J), as previously described ^88^ were from Jackson lab (Stock No. 011121) and backcrossed onto C57BL/6 background. *Atg5^flox/flox^* mice, originally from Mizushima lab at the University of Tokyo, were generated by introducing two *loxP* sites that flanks exon 3 of *Atg5* gene as previously described ^89^.

*Rubcn*^−/−^ mice (C57BL/6JGpt-*Rubcn^em1Cd61^*^00^/Gpt), *Rubcn^flox/flox^* mice (C57BL/6JGpt-*Rubcn^em1Cflox^*/Gpt) and *Axl^flox/flox^* mice (C57BL/6JGpt-*Axl^em1Cflox^*/Gpt) were generated via CRISPR/Cas9 technology by GemPharmatech (Nanjing, China). *Fcgr1*^−/−^ mice (NM-KO-18033) were generated via CRISPR/Cas9 technology by Shanghai Model Organisms Center (Shanghai, China). Briefly, for *Rubcn*^−/−^ mice, two selected gRNAs (sgRNA1: 5’-CGTCTCTCAGTAGAGCTCCA-3’, sgRNA2: ATAGCAGCAGTTGCTTAATG), targeting all *Rubcn* isoforms were constructed and transcribed, then microinjected together with *Cas9* mRNA into cytoplasm of zygotes of C57BL/6 mice to generate *Rubcn*^−/−^ mice. Founder mice were screened for deletion alleles. One allele, which deleted 6100bp including exon 3-6 and caused premature stop at amino acid 115, was isolated and backcrossed with C57BL/6 mice. Homozygous *Rubcn*^−/−^ mice are viable and fertile. To generate *Fcgr1*^−/−^ mice, three selected gRNAs (sgRNA1: 5’-cagtaaatctgaggataacctgg-3’, sgRNA2: 5’-aacagtaccataatgcaagcagg-3’, sgRNA3: 5’-TTGTCGAATGTTTGACCTTATGG-3’) were *in vitro* transcribed and microinjected with *Cas9* mRNA. One allele, deleting all coding sequence was isolated and backcrossed with C57BL/6 mice. Homozygous *Fcgr1*^−/−^ mice are viable and fertile. For *Rubcn^flox/flox^* mice, two gRNAs same to *Rubcn*^−/−^ mice were used for *loxP* site engineering. Two *loxP* sites that flank exons 3 and 6 were inserted into the endogenous *Rubcn* locus via CRISPR/Cas9-mediated homologous recombination with donor template. For *Axl^flox/flox^*mice, two selected gRNAs (sgRNA1: 5’-GACAAGAAACAGTAACCGGG-3’, sgRNA2: 5’-CCGATGTGAACCGATGCTGT-3’) were used for engineering of *loxP* sites. Two *loxP* sites flanking exon 2 and 17 of *Axl* were inserted into *Axl* endogenous locus via CRISPR/Cas9-mediated homologous recombination.

### Genomic PCR

Two mm of mouse tails were cut and lysed in 150 µL of 50 mM NaOH. The mixture was heated at 95°C for 30 minutes and then cooled to room temperature, followed by addition of 12 µl of 1 M Tris-HCl (pH 6.8) for neutralization. After centrifugation for 30 seconds at 3000 rpm, the supernatant containing genomic DNA was collected for PCR genotyping or stored at −20°C. PCR reaction was performed with 20 µL of mixture containing 1 µL of the supernatant, 1 µL of 10 µM forward primer, 1µL of 10 µM reverse primer, 10 µL of 2X Flash HS PCR Master mix (Accurate Biology, #AG12301) and 7 µL of distilled water. After PCR, 7 µL of amplified PCR products were loaded on a 2% agarose gel to determine mouse genotypes. The genotyping primers for flox alleles were listed in the Table 1.

### Contusive spinal cord injury (SCI)

Spinal cord contusion injury was performed at thoracic level (T9-10). In brief, adult mice (8-10 weeks old) were anesthetized by intraperitoneal injection of 2.5% tribromoethanol (Avertin, Mecklin, Cat#75-80-9) at 0.1 ml/10 g body weight. Following the loss of pedal reflexes, a T8-T10 laminectomy was performed to completely expose the spinal cord. Contusion was delivered at T9 using a precision impactor (RWD 68099 II, RWD Life Science) fitted with a 1.3 mm diameter impact head. The impactor parameters were set at a speed of 1.5 m/s, a depth of 0.6 mm, and a dwell time of 0.5 s. After hemostasis, the muscles and skin were sutured hierarchically with a mouse wound suture device. Mice were warmed after surgery until they were fully awake. The bladder was auxiliary emptied twice per day until urination function was restored. Sham-operated mice were subjected to laminectomy without contusion.

### Transient middle cerebral artery occlusion (tMACO) mouse model

The middle cerebral artery occlusion (MCAO) was performed on adult mice (20-25g) by transient occlusion of the middle cerebral artery for 60 minutes, to induce focal cerebral ischemia. Briefly, mice were anesthetized with sodium pentobarbital (100 mg/kg, intraperitoneal) before surgery. First, the common carotid, internal carotid, and external carotid artery were separated, then a silicone-coated filament (MEYUE, Cat# M8502) was inserted through the external carotid artery, and into the internal carotid artery until resistance was felt where it reached the middle cerebral artery (approximately 9-10 mm from common carotid artery bifurcation). 60 minutes after MCAO, the filament was withdrawn to allow reperfusion. Body temperature was maintained at 37.0 ± 0.5 °C during the operation. Sham-operated mice underwent the same surgical procedures, including vessel exposure, but had no silicone filament inserted. At indicated time-points after ischemia-reperfusion, mice were sacrificed.

### Active induction of EAE

The active EAE model was induced by MOG_35-55_ peptide MEVGWYRSPFSRVVHLYRNGK (Sangon Biotech, Cat#T510219-0001) as previously described with minor modifications ^90^. Briefly, EAE induction was performed with 7- to 8-week-old female C57BL/6J mice. Mice were subcutaneously (in the back region) injected with 200 μg of MOG_35-55_ peptide in Complete Freund’s Adjuvant (CFA, Thermo Fisher, Cat#77145) containing 5 mg/ml of heat-killed *Mycobacterium tuberculosis* (H37 Ra strain, BD Biosciences, Cat#231141). On the day of immunization and 48, 72 hours later, the mice were injected intravenously with Pertussis toxin (200 ng/mouse, List Biologicals, Cat#181). Mice were examined and scored daily for EAE symptoms as follows: 0, no obvious changes in motor function compared to non-immunized mice; 0.5, limp tail tip; 1, limp tail; 1.5, limp tail and hind leg inhibition; 2, limp tail and weak hind legs; 2.5, limp tail and dragging of hind legs; 3, paraplegia (complete paralysis of two hind limbs); 3.5, paraplegia and hind legs are together on one side of body; 4, paraplegia and partial front leg paralysis; 4.5, paraplegia and partial front leg paralysis, no movement around the cage; 5, moribund state or death.

### Laser speckle contrast imaging

To confirm the occlusion of the distal MCA, cerebral blood flow of MCAO and sham-operated mice was monitored with laser speckle contrast imaging system (LSCI) (RWD Life Science Co., Ltd, China) before and after surgery. LSCI is based on the blurring of scattered laser interference mode by blood cell flow to show the blood circulation in real time. In brief, mice were anesthetized with sodium pentobarbital (100 mg/kg, intraperitoneal), cut along the midline to remove the skull skin. The acquisition parameters were 50 frames per second, recorded continuously for 5s, and a total of 250 frames were collected. In order to evaluate the change of speckle contrast over time, the region of interest (ROI) was selected and centered on the laser-irradiated cortical area.

### Gilteritinib administration

Wildtype mice were administrated with Gilteritinib (MedChemExpress, Cat#HY-12432) in corn oil via oral gavage one day prior to injury. Corn oil alone was used as vehicle control. Mice in both vehicle and Gilteritinib group underwent spinal cord injury as described above. These injured mice were administrated with 100µL of 5mg/kg Gilteritinib or corn oil once per day for 5 days a week by oral gavage till day 18. BMS score was daily assessed. The mice were sacrificed at the time points as indicated when spinal cord tissues were harvested for histology and immunostaining analysis. No overt adverse events, including death case, weight loss, bleeding, peripheral edema, dyspnea, vomiting and diarrhea were noticed during the 42-day recovery window in mice.

### Basso Mouse Scale (BMS)

The locomotor behavior of mice was assessed using the Basso Mouse Scale (BMS)^91^. Briefly, BMS scores ranging from 0 to 9 were assigned by two well-trained investigators who were blinded to the experimental groups. Each mouse was individually placed in an acrylic transparent enclosure (30 cm × 25 cm × 20 cm) lined with padding, which served as an open field. This setup prevented escape while allowing clear observation. Animals from different treatment groups were randomly assigned to cages prior to testing. BMS scoring was based on hindlimb joint movement, weight support, trunk position and stability, stepping coordination, paw placement, and tail control, with scores ranging from 0 (no movement) to 9 (normal locomotion). Before testing, mice were habituated to handling and to the open field environment. BMS evaluations were conducted at baseline (pre-injury) and daily for 6 weeks post-injury.

### Rotarod test

Motor coordination was assessed using an accelerated rotarod test (Med Associates Inc., model ENV-574M, VT, USA). During the training phase, mice were habituated to the apparatus by being placed on the stationary rod for 3 mins, followed by adaptation at a constant speed of 4 rpm for 1 min. On the following day, formal testing was conducted. Before testing, mice were again allowed to adapt to the rotarod at 3 rpm for 1 min, after which they were rested for 5 min. During the test session, the rotarod was accelerated from 4 to 40 rpm over a 300s period. Each mouse underwent three trials, and the latency to fall from the rod was recorded for each trial.

### Grid walking test

The grid walking test was used to assess motor coordination and balance by quantifying foot placement errors during locomotion. One day before the formal experiment, mice were placed on the grid apparatus for a 3-min habituation session. On the test day, mice were placed on an elevated wire grid platform (35 × 30 cm, grid size 12 × 12 mm, elevated 40 cm above the floor) and allowed to explore freely for 5 minutes. Behavior was recorded using a smartphone camera at a resolution of 1080p and a frame rate of 60 frames per second. An investigator blinded to the genotype or treatment counted the number of foot slips, which were defined as instances when either the left or right hindlimb slipped below the grid surface. Only hindlimb movements were analyzed, and the first 100 steps of each hindlimb were independently assessed. The percentage of slips (foot faults) was calculated by dividing the number of slips by the total number of hindlimb steps.

### 3D motion-capture kinematic analysis

3D hindlimb kinematics were quantified using a markerless motion-capture system as previously described ^73^, with minor modifications. Locomotor behavior was recorded using a 3D-AI Animal Behavior Analysis System (Shenzhen Bayone BioTech Co., Ltd., Shenzhen, China) equipped with five synchronized Intel RealSense D405 depth cameras. Four cameras were positioned around the treadmill, and one additional camera was mounted above the recording arena. All five cameras were jointly calibrated to establish a common 3D coordinate system. Following calibration, synchronized recordings from the peripheral cameras were processed using a multi-view geometric triangulation pipeline to reconstruct 3D coordinates of body landmarks for subsequent hindlimb kinematic analysis ^92^.

For gait recordings, mice were recorded while walking on a motorized treadmill moving at a constant speed of 1 cm s⁻¹. Video data were acquired from the synchronized peripheral cameras at 60 frames per second. Each animal was recorded continuously for 3 min to ensure the capture of a sufficient number of complete locomotor cycles for subsequent analysis.

For markerless tracking and 3D skeleton reconstruction, investigators blinded to experimental group allocation manually annotated 17 anatomical landmarks in approximately 1300 representative frames sampled from all recording, including: back, left_front_shoulder, left_front_elbow, left_front_wrist, left_front_toetip, right_front_shoulder, right_front_elbow, right_front_wrist, right_front_toetip, left_hind_hip, left_hind_knee, left_hind_ankle, left_hind_toetip, right_hind_hip, right_hind_knee, right_hind_ankle and right_hind_toetip. The annotated dataset was used to train a pose estimation model based on an HRNet-W48 backbone initialized with ImageNet-pretrained weights ^93^. Following automated pose estimation, 2D landmark detections from the synchronized peripheral cameras were reconstructed into 3D coordinates using a custom-developed multi-view reconstruction pipeline integrating camera calibration, multi-view landmark correspondence, and geometric triangulation. The resulting 3D skeletal trajectories enabled quantitative analysis of posture, locomotion, and hindlimb kinematics.

Reconstructed locomotor sequences were visually inspected, and frames containing tracking errors or reconstruction artifacts were excluded from subsequent analysis. Only accurately reconstructed gait cycles were included in the final dataset.

For kinematic analysis, reconstructed 3D coordinates were processed using custom-written Python scripts. The coordinates of the toe and hip were denoted as:

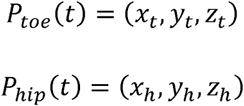

where *x*, *y*, and *z* correspond to the rostrocaudal, mediolateral, and vertical axes, respectively.

To account for body translation during locomotion, all landmarks were transformed into a body-centered coordinate system. For each frame, the projection of the back landmark onto the horizontal (XY) plane was defined as the origin of the body-centered coordinate system, while the global vertical (Z) axis was retained. Individual gait cycles were identified based on right hind paw trajectories and manually verified to exclude irregular or interrupted gait cycles. For each animal, 14–31 valid gait cycles were included for analysis.

All extracted kinematic parameters were subjected to quality assessment across gait cycles and animals to evaluate measurement reliability and their ability to capture distinct aspects of hindlimb locomotor function. Based on these criteria, three complementary parameters, including maximum foot forward reach, maximum hindlimb forward oscillation, and mean hip height, were selected as representative measures for subsequent statistical analyses.

Maximum hindlimb forward oscillation was defined as the maximal forward excursion angle of the hindlimb relative to the body vertical axis across all valid gait cycles. Positive values indicate forward swing. For each frame, the limb angle was calculated from the vector connecting the hip and toe landmarks. Results are expressed in degrees (°).

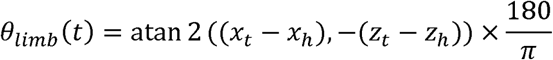

Maximum foot forward reach was defined as the maximal anterior displacement of the toe relative to the hip along the rostrocaudal axis over all frames from all valid gait cycles. Values are reported in centimeters (cm).

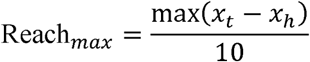

Mean hip height was defined as the average vertical position of the hip across all valid gait cycles. Values are reported in millimeters (mm).

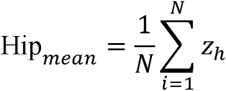

### Human brain samples

Postmortem human brain samples from individuals with or without stroke were obtained from the National Human Brain Bank for Development and Function (Chinese Academy of Medical Sciences and Peking Union Medical College, Beijing, China). This study was approved by Southern Medical University, and the China Human Brain Banking Consortium (Approval Number: 009-2014). All brain samples were collected following the standardized operational protocol established by the China Human Brain Bank Consortium. Blocks were selected from the basal ganglia, paraffin-embed sections were cut at the thickness of 5μm. Information on the donors, including group, age and gender, is summarized in Figure. S7A.

Fresh human brain tissue was obtained from one patient with epilepsy undergoing surgical resection of epileptogenic zones. The patient, a 28-year-old male, presented with symptomatic focal epilepsy and a surface electroencephalogram (EEG) pattern showing left temporal spikes and slow waves in the left medial temporal lobe, for which lesionectomy was performed with written and informed consent from patient’s family. The use of surgically resected tissues was approved by Southern Medical University. Before the surgery, an extensive pre-surgical workup was performed on the patient. Post-surgical histological examination of resected tissue was performed to confirm the nature of the tissue. A small part (∼2 cm^3^) of the planned surgical margin of resection was obtained for perivascular cell isolation.

### Reagents

Carboxyfluorescein succinimidyl ester (CFSE) (Cat#C1157) was purchased from Life Technologies (Carlsbad, CA). Matrigel matrix growth factor reduced (Cat#354230) was from BD Biosciences (San Jose, CA). 4-hydroxyl tamoxifen (4-OHT) (Cat#T176) and Green-conjugated Zymosan (Cat#IAK0110) were from Sigma. Rapamycin (Cat#553210) was purchased from EMD Millipore (Burlington, MA). DNase I (Cat#A610099-0001) from Sangon Biotech (Shanghai, China), fetal bovine serum (FBS) (Cat#A6904) was from Gibco (Waltham, MA). Recombinant mouse TGF-β1 (Cat#CK33) was from Novoprotein (Beijing, China). Gilteritinib (Cat#HY-12432) was from MedChemExpress (Monmouth Junction, NJ). Silicone-coated filaments were from Meyue BIO (Changsha, China). 2,3,5-triphenyltetrazolium chloride (TTC) (Cat#0765-1) was from Lablead (Beijing, China). Electron microscopy fixative (G1102) was purchased from Servicebio (Wuhan, China). Cytochalasin D (Cat#C11560) was from Acmec Biochemical (Shanghai, China), Dynasore (Cat#324410) was from Sigma, CA-074 methyl ester (CA-074me, Cat#A8239) was from APExBIO. Purified Recombinant Annexin V (Cat#556416) was from BD Pharmingen.

### Antibodies

Rabbit anti-MBP (#ab40390; IF 1:400) from Abcam (Cambridge, UK), rabbit anti-α-SMA (#ab124964; IF 1:400, WB 1:1000) from Abcam, and rabbit anti-PDGFRβ (clone 28E1, #3169; WB 1:1000) from Cell Signaling Technology (Danvers, MA). Rabbit anti-NG2 (#ab5320; IF 1:150) from Millipore (Billerica, MA). Goat anti-PDGFRβ (#AF385, #AF1042; IF 1:200) from R&D system (Minneapolis, MN), rat anti-PDGFRβ (#14-1402-81, IF 1:100) from Invitrogen (Carlsbad, CA). Goat anti-GFAP (#ab53554; IF1:400), rabbit anti-Ki67 (#ab15580; IF 1:400), chicken anti-mCherry (#ab205402; IF1:2000) and rabbit anti-GAPDH (#ab181602; WB 1:3000) from Abcam (Cambridge, MA). Rabbit anti-LC3 (#4108; IF 1:100; WB 1:1000) and rabbit anti-Atg5 (clone D5F5U, #12994; WB1:1000) from CST. Rat anti-Lamp1 (clone 1D4B; IF 1:100) from Developmental Studies Hybridoma Bank. Rat anti-CD31 (#550274; IF 1:10) from BD Biosciences (Franklin Lakes, NJ). Rabbit anti-Iba-1 (#019-19741; IF 1:400) from FUJIFILM Wako (Osaka, Japan). Rabbit anti-CC-1 (#OB-PRB070-01; IF 1:200) from Oasis Farm (Hangzhou, China), rabbit anti-NeuN (#Ab177487; IF 1:200) from Abcam, rabbit anti-NF-H/NF-200 (#N4142; IF 1:500) from Sigma, rabbit anti-AQP4 (#HPA014784; IF 1:200) from Atlas Antibodies (Stockholm, Sweden). Rabbit anti-Cathepsin B (#31718S; IF 1:500) from CST. Goat anti-Axl (#AF854; IF 1:200) from R&D system, rabbit anti-Rubicon (D9F7) (#8465; WB 1:1000) from CST, mouse anti-β-actin (#SC-47778; WB 1:3000) from Santa Cruz, rabbit anti-Desmin (#5332; IF 1:100) from CST, goat anti-5-HT (#20079; IF 1:3000) from ImmunoStar (Hudson, WI), rabbit anti-dMBP (#AB5864; IF 1:500) from Millipore, rabbit anti-P2RY12 from CST (#69766S, IF1:500), rat anti-F4/80 (#50-4801-82; IF 1:50) from Invitrogen, rabbit anti-S100β from Abcam (#ab52642; IF1:100), rabbit anti-Mfsd2a (#80302S, IF 1:500) from CST, rabbit anti-ZO-1 (#AF5145; IF 1:200) from Affinity Biosciences (Cincinnati, OH), rabbit anti-p-Smad3 (#AP0727 IF 1:150) from ABclonal (Woburn, MA), rabbit anti-Fibronectin (#ab23750; IF1:400) from Abcam, rabbit anti-Collagen I (#ab34710; IF1:400) from Abcam. Donkey-conjugated secondary antibodies (IF 1:1000) from Invitrogen. HRP-conjugated secondary antibodies (WB 1:3000) from Yeasen Biotech.

### Spinal cord single-cell preparation

Mice were anesthetized and perfused with ice-cold PBS containing 6 U/ml heparin sodium (BBI, Cat#A603251-0001) to remove circulating red blood cells. 5-mm sections of lesioned spinal cords from SCI mice or total spinal cords from sham control mice were collected rapidly and transferred to a tissue processing tube (RWD Life Science, Cat# SCT-25) containing 2 ml of digestion solution in the single-cell preparation kit (RWD Life Science, Cat#DHABE-5003). Spinal cords were enzymatically dissociated using a RWD TDS-5 Tissue Digestion System with SCT-25 tissue-processing tubes at 37℃ for 30 min. The suspension was sequentially filtered through 100 µm and 70 µm cell strainers, centrifuged (1200rpm, 4℃, 10 min) and the pellets were resuspended in 4 ml myelin-removal reagent. An equal volume of PBS was added to create a layered effect, followed by density-gradient centrifugation (3000g, 4℃, 10 min). After removing the upper solution and the myelin fragments in the middle layer, the remaining pellet was washed once with 4 ml PBS (1200 rpm, 4℃, 10 min) and resuspended in 1 ml ice-cold L15 medium. After addition of 1.7 ml of 22% BSA, a second density-gradient centrifugation was performed (2600 rpm, 4℃, 10 min) to remove residual myelin debris. The final pellet was resuspended in L15 medium for downstream assays.

### Fluorescence-activated cell sorting (FACS) of perivascular cells

Perivascular cells were isolated from PDGFRβ-Cre::Ai14 mice following preparation of single-cell suspensions from spinal cord tissues. Cells were resuspended in L-15 medium containing 0.5% bovine serum albumin (BSA) and incubated with fluorophore-conjugated antibodies on ice for 20 min. After two washes with FACS buffer (PBS containing 0.5% BSA), cells were labeled with APC-conjugated anti-CD45 antibody (1:200; BD Biosciences, Cat#559864) to exclude immune cells and with DAPI (0.5 μg/mL) to exclude dead cells. Perivascular cells were purified by sorting tdTomato^+^ CD45^−^ DAPI^−^ cells using a Beckman Coulter CytoFLEX SRT cell sorter (Beckman Coulter Life Sciences, Indianapolis, IN, USA) at the Southern Medical University Flow Cytometry Core Facility. Flow cytometry data were analyzed using FlowJo software (version 10.8.1; FlowJo, LLC, Ashland, OR, USA).

### Isolation and culture of primary mouse and human perivascular cells

Mouse primary perivascular cells were enzymatically isolated from mouse brains or spinal cords. 7-day old mice (n=6) were euthanized, and mouse brains or spinal cords were collected using sterile scissors. Human brain pericytes were isolated from surgically resected brain tissue from an epilepsy patient as described above. The post-surgical tissue was transported in a cold human pericyte medium (ScienCell Research Laboratories, Cat#1201), containing 1% pericyte growth supplement, 10% fetal bovine serum (FBS) and 1% penicillin/streptomycin. The procedures for isolating mice and human perivascular cells are similar. The collected CNS tissues were placed in 100 mm Petri dish with either cold Minimum Essential Medium (MEM, Corning, Cat#45000-384), or cold Ham’s Nutrient Mixture F12 (Sigma-Aldrich, Cat#51651C-1000ML) with sodium bicarbonate. Whole spinal cords were used for spinal cord perivascular cell isolation, whereas for brain perivascular cell isolation, the meninges, olfactory bulbs, cerebellum, medulla and large vessels were removed from the brains. The remaining brain tissues were minced into small pieces in another 100 mm Petri dish with cold Hams F12 medium. The minced brain pieces were washed once with Hams F12 medium and then centrifuged at 1200 rpm for 5 min at 4℃. Following wash, the tissues were incubated at 37 ℃ for 40 minutes in an enzymatic solution that contains 1 mg/ml papain (Molecular Probes, Cat#IC10256680) and 2000U of DNase I (Sangon Biotech, Cat#A610099-0001) in Hams F12 medium.

The digested brain tissues were homogenized by 5 passes through an 18-gauge needle and subsequently 5 passes through a 21-gauge needle. The microvessels will mostly survive trituration and be separated from the brain fragments. After mixing the homogenized brain parts with 1.7 volumes of 22% bovine serum albumin (BSA) in PBS, they were centrifuged for 10 minutes at 2600 rpm, followed by careful removal of the top layer of myelin and lipid layer. The microvessels and cell pellets on the bottom were resuspended in 6 ml of Hams F12. For microvessel collection, gradient centrifugation was performed, initially at 1200 rpm for 5 min, then at 1000 rpm 5min, 800 rpm 5min, 500 rpm 5min. Mouse pericyte medium (ScienCell Research Laboratories, Cat#1231), containing 1% pericyte growth supplement, 10% fetal bovine serum (FBS), 1% penicillin/streptomycin was used to resuspend the collected microvessels. The resuspended microvessels were next placed in two wells of a 6-well plate that were coated with 4.8 mg/ml gelatin (Sigma Cat#G1890) for 2 h at 37 °C. Perivascular cells sprout from the microvascular buds that attach effectively to plates. The cells usually reached 100% confluence after three days of culture. The confluent cultures were trypsinized using 0.025% trypsin and then passaged 1:4 onto 6-well plates that were precoated with fresh gelatin.

*Axl*^−/−^, *CR3*^−/−^ and *Rubcn*^−/−^ mouse perivascular cells were isolated similarly from *Axl*^−/−^, *CR3*^−/−^ and *Rubcn*^−/−^ mice, respectively. Perivascular cell sprouting and outgrowth were imaged using 20× or 40× objective with an inverted MF52-N fluorescence microscope (MSHOT, China). Following five passages, nearly all cells stained positive for common perivascular cell markers, including PDGFRβ, NG2, α-SMA and desmin as shown by laser scanning confocal microscopy (Fig. S9).

### Generation of *Atg5* knockout perivascular cells

*Atg5* knockout in perivascular cells was achieved by 4-hydroxy tamoxifen (4-OHT, Sigma, Cat#T176) treatment of primary perivascular cells that were isolated from the brains of *CAG-Cre^ERT^*::*Atg5^flox/flox^*littermates. Isolation, characterization and culture of *CAG-Cre^ERT^*::*Atg5^flox/flox^*perivascular cells were performed as described above. The isolated perivascular cells were treated with 4-OHT at 200nM in EtOH for 48 hr to induce *Atg5* gene excision, and perivascular cells without 4-OHT treatment (only EtOH) were used as control. *Atg5* knockout cells were analyzed by Western blot detection of Atg5 protein and LC3-I/II conversion.

### Bone marrow-derived macrophage (BMDM) isolation

Bone marrow-derived macrophages (BMDMs) were isolated from C57BL/6J mouse bone marrow. Briefly, first femurs from 8-12 weeks old mice were dissected. Then, the marrow cavity of femurs was flushed with DMEM and bone marrow cells were seeded into DMEM (Gibco, Cat# C11995500BT) containing 15% medium supernatant from L929 cells (a source of M-CSF) after filtering through a 70-μm mesh. BMDMs were cultured in medium supplemented with 10% serum. The medium was replaced 2 days after plating, changed again on day 4, and the culture was maintained until day 7. BMDMs were maintained at 37℃, with 5% CO_2_.

### Primary astrocyte isolation

Primary astrocyte cultures were established from the whole brain of P1-P2 neonatal C57BL/6J mice. Briefly, meninges and blood vessels were carefully removed in pre-cooled DMEM (Gibco, Cat# C11995500BT). The whole brain was then transferred to a tube containing 0.025% trypsin (Viva cell, Cat# C3530-0500) and triturated for 5 minutes. The cell suspension was quenched with 100% FBS, centrifuged at 1000 rpm for 8 minutes, and the supernatant was discarded. The cell pellet was resuspended in astrocyte culture medium (DMEM/F12 1:1, supplemented with 1% Triple Antibiotic Solution and 10% FBS), filtered through a 70-μm strainer, and plated in T75 flask. Cultures were maintained in an incubator at 37°C with 5% CO₂. The medium was changed every 3-4 days. To separate astrocytes from microglia, the cultures were shaken at 240 rpm for 6 hours, and the detached cells were removed.

### Myelin debris preparation and fluorescent labeling

Myelin debris were isolated from the brains of 2-month-old C57BL/6J mice as previously described ^34^. Endotoxin contamination was below the detection limit as determined by the Limulus amebocyte lysate assay. For fluorescence labeling, purified myelin debris were incubated with 50 nM carboxyfluorescein succinimidyl ester (CFSE), a membrane-permeable amine-reactive fluorescent dye, for 30 min at 37°C. Excess dye was quenched with 100 mM glycine in PBS, followed by centrifugation at 14,000 rpm for 10 min. The labeled myelin debris were washed three times with PBS, with centrifugation after each wash, and finally resuspended in PBS for subsequent experiments. CFSE fluorescence colocalized with myelin basic protein (MBP) immunostaining in myelin-laden perivascular cells (Fig. S16B), confirming the fidelity of CFSE labeling for tracking myelin debris. MBP-deficient myelin debris were prepared from *Mbp*^−^/^−^ (*Mbp^shi^*) mice using the same isolation and labeling procedures. Unless otherwise indicated, myelin debris were used at a final concentration of 1 mg/mL for all in vitro experiments.

### *In vitro* phagocytosis of myelin debris

For uptake of myelin debris, perivascular cells were co-cultured with 1 mg/mL of CFSE-labeled myelin for indicated time periods in 24-well gelatin-precoated plates. Non-ingested myelin debris was eliminated by EDTA for 30 seconds and citric acid for 1 minute. The internalized CFSE-labeled myelin debris was analyzed by confocal imaging or flow cytometry detection. For confocal imaging, myelin debris-laden perivascular cells were fixed with 4% paraformaldehyde (PFA), stained with α-SMA to label perivascular cell identity and imaged with Nikon A1 (Nikon, Japan) or Leica TCS SP8 (Leica, Germany) laser scanning confocal microscope.

To test the role of actin and dynamin in myelin debris uptake, 0.5 μM of cytochalasin D (inhibitor of actin assembly) or 80 μM dynasore (dynamin inhibitor) were added in the culture medium. Gilteritinib was used at a final concentration of 1 μM for primary perivascular cells and astrocytes and 250 nM for bone marrow-derived macrophages. Cells were pretreated with the drug for 1 hr before the addition of myelin debris, and the drug was maintained in the culture medium throughout the phagocytosis assay. No detectable cytotoxicity was observed in any cell type at the concentrations used.

For tests with perivascular cells lacking specific receptors, cells were isolated from corresponding knockout mice (*Axl^−/−^*, *Fcgr1^−/−^*, and *CR3^−/−^*). Cells were treated with CFSE-labeled normal myelin debris or CFSE-labeled MBP-deficient myelin debris for 72 hr and assayed for CFSE fluorescence within perivascular cells. For serum-dependent phagocytosis of myelin debris, wild-type perivascular cells were cocultured with CFSE-labeled myelin debris for 72 hr in the presence of 0% (serum free), 5% and 10% of FBS, respectively.

### *In vitro* uptake of necrotic neuronal bodies and zymosan

Mouse neuroblastoma cell line, Neuro-2a (N2A, CCL-131) was purchased from American Type Culture Collection (Manassas, VA). Neuro-2a were cultured in DMEM containing 5% or 10% FBS and 1% penicillin/streptomycin. To induce differentiation to neurons, N2A cells were cultured in DMEM medium containing 1% FBS and 20 µM retinoic acid for 3 days as previously described ^94^. All cells tested negative for mycoplasma contamination. Neuronal bodies and debris were generated from N2A-differentiated neuronal cells by repeated freeze-thaw cycles (3×). CFSE-labelled neuronal cell bodies or debris were added to perivascular cells at 10:1 ratio and cultured for indicated time points. For zymosan uptake, CFSE-conjugated zymosan was incubated with perivascular cells or BMDMs at 1mg/mL for indicated time points.

### Myelin debris uptake by perivascular cells on Matrigel

To generate microvessel-like tubular structures, primary perivascular cells were seeded onto polymerized Matrigel-coated coverslips in 24-well plates and cultured for 24 hr at 37°C. Cells were then incubated with 1 mg/mL CFSE-labeled myelin debris. Following incubation, non-internalized myelin debris was carefully removed by gentle washing to preserve the integrity of the tubular structures. Cells were fixed with 2% PFA and subjected to immunostaining. Confocal images were acquired using a Nikon A1 laser scanning confocal microscope (Nikon, Japan). Orthogonal (XY and YZ) views of the tubular structures were analyzed using Nikon NIS-Elements software, and 3D reconstructions were generated with Imaris software (Bitplane) to visualize the spatial distribution of internalized myelin debris within the microvessel-like structures.

### Recombinant Annexin V blocking assay

To mask phosphatidylserine (PtdSer) on myelin debris, purified myelin debris were incubated with 8 μL recombinant Annexin V (0.5 mg/mL; BD Pharmingen, Cat#556416) in 50 μL Annexin V binding buffer at room temperature for 3 hr. Control myelin debris was incubated in Annexin V binding buffer alone under identical conditions. Following pretreatment, Annexin V-coated or control myelin debris was added to primary perivascular cell cultures in 24-well plates and cocultured for 24 hr. Cells were then processed for immunocytochemistry and subsequent quantitative analyses.

### Myelin debris degradation assay

Primary perivascular cells (2×10^4^ cells per well) were seeded in 24-well plates and incubated with CFSE-labeled myelin debris (final concentration, 1 mg/mL) for 24 hr to allow cargo uptake. Cells were then washed thoroughly with PBS to remove non-internalized myelin debris and cultured for additional 24 hr in fresh medium containing the cathepsin B inhibitor CA-074Me (10 μM). Control cultures were treated with DMSO. At the end of the treatment, cells were fixed and processed for immunocytochemistry. Confocal images were acquired, and the intracellular CFSE signal was quantified to assess myelin debris degradation.

### Oil Red O (ORO) staining

For ORO staining, cells were fixed and dehydrated in 100% propylene glycol for 5 min, followed by incubation with 0.5% ORO solution at 60°C for 8 min. Samples were then processed in 85% propylene glycol for 5 min, rinsed three times with distilled water, and imaged using a confocal laser scanning microscope.

### Plasmid construction

For knockdown of human *AXL* and mouse *Lrp1* in primary perivascular cells, shRNA sequences were cloned into the PLKO.1 vector (Addgene #8453) encoding a U6 promoter-driven shRNA expression cassette and an hPGK promoter-driven puromycin resistance gene for selection. Briefly, shRNA oligonucleotides were annealed and ligated into AgeI/EcoRI digested PLKO.1 vector by T4 DNA ligase(Accurate #AG11801). Scramble shRNA as a control is non-targeting. The shRNA oligonucleotides used in this study are listed below:

Mouse *shLrp1#1*: 5-GGAAGTGATGGGAAGTCTTGT-3
Mouse *shLrp1#2*: 5-GCTCACACCGAGATATCTTTG-3
scramble: 5-CCTAAGGTTAAGTCGCCCTCG-3
Human *shAXL#1*: 5-GCGGTCTGCATGAAGGAATTT-3
Human *shAXL#2*: 5-CGAAAGAAGGAGACCCGTTAT-3

### Lentivirus production and transduction

HEK293T cells were seeded in 10-cm culture dishes and transfected at 70%–80% confluency. Before transfection, the culture medium was replaced with 4 mL serum-free DMEM (Gibco, Cat#C11995500BT). The lentiviral packaging plasmids psPAX2 (Addgene, Cat#12260), pMD2.G (Addgene, Cat#12259), and the PLKO.1 target transfer plasmid were mixed at a mass ratio of 1.2:0.6:2, diluted in 100 μL DMEM, and combined with 4 μL polyethyleneimine (PEI; LABLEAD, Cat#P4000). After incubation at room temperature for 15 min, the transfection mixture was added to the cells. After 8–12 hr, the medium was replaced with 6 mL DMEM containing 10% fetal bovine serum (FBS). Lentivirus-containing supernatants were collected at 24, 48, 72, and 96 hr after transfection, pooled, filtered, and concentrated before use.

Primary perivascular cells were transduced with concentrated lentivirus in the presence of polybrene (1:1000; Santa Cruz Biotechnology, Cat#sc-134220) for 24 hr, followed by puromycin selection (5 μg/mL) for 36 hr. Gene knockdown efficiency was confirmed by RT-qPCR.

### RNA sequencing and analysis

Total RNA was isolated from sorted cells or cultured cells with the miRNeasy Micro Kit (QIAGEN, Cat#217084) according to the manufacturer’s instructions. RNA integrity was verified on an Agilent 2100 Bioanalyzer (RIN ≥ 8.5). Strand-specific libraries were generated with the TruSeq Stranded mRNA Library Prep Kit, pooled by sample-specific indexes, and sequenced on Illumina HiSeq 4000 as 2 × 150 bp paired-end reads following the manufacturer’s protocols. Raw FASTQ reads were trimmed and filtered with Cutadapt v1.9.1 as follows: adapters and low-quality bases (Phred < 20) were removed, the maximum allowed error rate during adapter trimming was set to 0.1, reads shorter than 75 bp or containing >10 % ambiguous (N) bases were discarded. Clean reads were then aligned to the *Mus musculus* reference genome (GRCm39, Ensembl release 107) using HISAT2 v2.2.1 with default settings. Gene-level counts were obtained with HTSeq-count (HTseq v0.6.1) using Ensembl gene annotations (release 107) and the read counts were expressed as fragments per kilobase of transcript per million (FPKM). Differential expression analysis was performed in R with DESeq2 v1.34.0. Genes with |log2Foldchange| > 0.5 or 1 (see figure legend) and FDR adjusted *P*-value (Benjamini–Hochberg) < 0.05 were considered differentially expressed. Volcano plots, heatmaps and GSEA were visualized by using ggplot2 4.0.1, pheatmap 1.0.13, and GSEABase 1.68.0, respectively.

### Transmission electron microscopy

Primary perivascular cells, with or without myelin debris treatment (72 hr), were cultured in 10-cm gelatin-coated culture dishes and collected for transmission electron microscopy (TEM). Cells were fixed in 2.5% glutaraldehyde for 5 min at room temperature and pelleted by centrifugation at 3,000 rpm for 5 min. The cell pellets were resuspended in fresh 2.5% glutaraldehyde and further processed by the Core Facility of Biomedical Sciences at Xiamen University. Briefly, samples were post-fixed in 1% osmium tetroxide, en bloc stained with uranyl acetate, dehydrated through a graded acetone series, and embedded in SPI-Pon 812 resin (SPI Supplies). Ultrathin sections were cut, counterstained with uranyl acetate and lead citrate, and examined using a Hitachi HT-7800 transmission electron microscope.

### CLEM analysis

In-house letter markers were generated on glass-bottom confocal dishes (NEST, Cat#801001) using an ion sputter coater (Leica ACE600) to facilitate ROI localization during correlative imaging. Briefly, a mold with letter marker was positioned onto the confocal dish, followed by the sputtering of platinum target at 35 mA for 70s to acquire letter-marked dish for confocal microscopy. 5×10^3^ perivascular cells were seeded in the letter-marked dish, subjected to an uptake assay using CFSE-labeled myelin debris, and fixed with 2.5% glutaraldehyde. Fluorescence images were immediately acquired at specific positions using a confocal microscope (Leica SP8). For subsequent transmission electron microscopy, samples were washed three times with 0.1 M PBS (15 min per wash at 4°C). They were then postfixed with 1% OsO4-1.5% tetrapotassium ferrocyanide in 0.1 M PBS for 1 h at 4℃. After brief rinses with ddH₂O, the samples were treated with 1% thiocarbohydrazide for 20 min at room temperature, and then washed in ddH_2_O and incubated overnight in 2% aqueous uranyl acetate at 4°C. Following a final rinse with ddH₂O, the samples were then dehydrated stepwise in ethanol through ascending concentration gradient on ice. They were subsequently infiltrated with a graded series of ethanol and 812 resin mixtures (ethanol: resin ratios of 3:1 for 40 min, 1:1 for 2 h, and 1:3 for 2 h), followed by overnight infiltration with pure 812 resin and polymerization at 70°C for 18 h. The dishes were carefully removed. ROIs were trimmed from the resin based on the fluorescence images, and ultrathin (70-nm thickness) sections were cut and inspected with a transmission electron microscope (HT-7800; Hitachi).

Two-dimensional image correlation or overlay between fluorescence microscopy and EM images was performed using the BigWarp plugin in ImageJ (v.1.54p). A manual, landmark-based linear registration was carried out by selecting six pairs of corresponding, morphologically unambiguous landmarks in the fluorescence (moving) and EM (fixed) images. Based on these landmarks, a linear transformation was computed and applied to the fluorescence image. The transformed fluorescence image was resampled into the coordinate space of the EM image using bilinear interpolation.

### *In vitro* perivascular cell proliferation assay

Following myelin debris engulfment, the proliferation index of perivascular cells was assessed by Ki67 staining. 2×10^4^ perivascular cells were seeded and grown for 24 hr on coverslips in 24-well plates. After treatment with or without 1 mg/mL of CFSE-labeled myelin debris for 72 hr. The fixed cells on coverslips were stained with Ki67 antibody and DAPI to label proliferating cells and nuclei, respectively. Fixed cells were imaged using a laser scanning confocal microscope and Ki67 and DAPI positive cells were counted in Fiji. To quantify the total cell counts, a hemocytometer was used. Briefly, 1×10^4^ perivascular cells were seeded and grown for 24 hours on coverslips in 24-well plates, followed by 72 hr treatment with or without myelin debris. The perivascular cells in 24-well plates were trypsinized and counted using a hemocytometer.

### *In vivo* Matrigel plug angiogenesis assay

The in vivo Matrigel plug angiogenesis assay was performed as previously described with minor modifications ^34^. Briefly, primary wild-type or *Axl^−/−^* perivascular cells were cultured in the presence or absence of myelin debris (1 mg/mL) for 72 hr and resuspended in ice-cold Matrigel at 4°C. A total of 100 μL of the Matrigel-cell suspension was subcutaneously injected into the left and right abdominal flank of wild-type recipient mice using a 26-gauge needle fitted to a Hamilton syringe. Each mouse received two independent Matrigel plugs, one at each injection site. Seven days after implantation, the Matrigel plugs were harvested, fixed, and processed for routine histological and immunofluorescence analyses. Ki67 and PDGFRβ immunostaining was performed to evaluate the proliferation of perivascular cells within the Matrigel plugs.

### Myelin debris microinjection assay

CFSE-labeled myelin debris was microinjected into the spinal cords of uninjured wild-type or *Axl^−/−^* mice, with PBS injection serving as the control. Mice were anesthetized with isoflurane, and a laminectomy was performed at the T10 vertebral level to expose the spinal cord. A total volume of 2 μL of CFSE-labeled myelin debris or PBS was delivered into the T10 spinal cord using a 32-gauge needle attached to a Hamilton syringe. To minimize tissue damage and facilitate diffusion, the injection was performed in two sequential 1 μL injections separated by a 5 min interval. After each 1 μL injection, the needle was left in place for an additional 5 min before the second injection or needle withdrawal. Five days after microinjection, mice were anesthetized and transcardially perfused with 4% PFA. Spinal cords were collected, cryosectioned, and processed for immunostaining and imaging as described below.

### Sulfo-NHS-Biotin permeability assay

Sulfo-NHS-biotin (4 mg in 400 μL per mouse; APExBIO, Cat#A8001) was administered by intravenous injection. Mice were then anesthetized, and brains were rapidly dissected without transcardial PBS perfusion to preserve the intravascular distribution of the tracer. Brains were immersion-fixed in 4% PFA for 3 hr at room temperature, washed overnight in PBS, and cryoprotected in 30% sucrose at 4°C. Tissues were embedded in OCT, frozen at −20°C, and sectioned coronally at 25 μm using a cryostat (CM1950, Leica). Brain sections were incubated with Streptavidin-HyperFluor™ 555 (1:25; APExBIO, Cat#K4404) and DAPI for 1.5 hr at room temperature. Images were acquired using a Leica SP8 laser scanning confocal microscope.

### Histology and immunofluorescence staining

Mice were anesthetized with isoflurane and transcardially perfused with 0.9% saline followed by 4% PFA. Brains and spinal cords were dissected, post-fixed in 4% PFA, cryoprotected in 30% sucrose overnight at 4°C, embedded in OCT compound (Sakura, Cat#4583), and sectioned at 20 μm using a cryostat (CM1950, Leica). Sections were mounted onto glass slides and stored at −40°C or −80°C until use.

For immunohistochemical staining of tissue sections, samples were blocked for 1 hr at room temperature in PBS containing 0.3% Triton X-100 and 1% bovine serum albumin (BSA), followed by incubation with primary antibodies overnight at 4°C. After washing with PBS, sections were incubated with the appropriate fluorophore-conjugated secondary antibodies for 1.5 hr at room temperature.

For immunocytochemical staining of cultured cells, cells were fixed with 4% PFA for 20 min, permeabilized with 0.2% Triton X-100 for 8 min, and blocked with 5% BSA for 30 min at room temperature. Samples were incubated with primary antibodies overnight at 4°C, washed with PBS, and incubated with the appropriate secondary antibodies for 1.5 hr at room temperature.

Tissue sections and cultured cells were mounted using Fluoro-Gel mounting medium in Tris buffer (Electron Microscopy Sciences, Cat#17985-11) and imaged by confocal microscopy.

### Histology and immunofluorescence staining of human brain tissues

Paraffin-embedded human brain tissue samples were sectioned at 5 μm, mounted onto adhesive glass slides, and dried at 65°C for 2 hr. Sections were deparaffinized in xylene (2 × 5 min), rehydrated through a graded ethanol series (100%, 100%, 95%, 80%, and 70%; 5 min each), and rinsed in distilled water. Heat-induced antigen retrieval was performed in 10 mM sodium citrate buffer (pH 6.0) by microwave heating at medium power for 2 min followed by medium-low power for 8 min. Sections were then allowed to cool to room temperature, blocked with 5% bovine serum albumin (BSA) for 30 min, and incubated overnight at room temperature with primary antibodies against MBP (1:200; Abcam, Cat#ab40390) and PDGFRβ (1:100; R&D Systems, Cat#AF385) diluted in 2.5% BSA. After washing with PBS, sections were incubated with the appropriate fluorophore-conjugated secondary antibodies diluted in 5% BSA for 1 hr at room temperature. Sections were subsequently washed, mounted with Fluoro-Gel mounting medium (Electron Microscopy Sciences), and imaged by confocal microscopy.

### HCR-FISH with immunofluorescence staining in spinal cords

Detection of mRNA in spinal cords was performed using a modified hybridization chain reaction (HCR)-based fluorescent in situ hybridization (FISH) method ^95^. Mouse spinal cords were collected, OCT embedded, sectioned and fixed as described for immunohistochemical staining. Subsequently, the slides were immersed in 100% ethanol for 5 minutes at room temperature (RT), followed by a wash in 1× Dulbecco’s Phosphate-Buffered Saline (DPBS, Solarbio, P1004). The sections were air-dried, and a hydrophobic barrier was drawn around the tissue using an immunohistochemistry pen (Vectorlabs, H-400). A humidified chamber and probe hybridization buffer (PHB) were pre-heated to 37°C. Samples were pre-hybridized by applying 100 μL of PHB and incubating at 37°C for 10 min in the humidified chamber. A probe solution was prepared by diluting 2 μL of each target probe into 100 μL of pre-warmed PHB. Following removal of the pre-hybridization solution, samples were hybridized with 100 μL of the probe solution at 37°C for >8 h in the humidified chamber. Excess probes were removed by washing the samples four times (15 min per wash) with pre-warmed Probe Wash Buffer at 37°C, followed by two 5-min washes with 5× saline-sodium citrate with Tween 20 (SSCT) at RT. Samples were pre-amplified by applying 200 μL of Amplifier Buffer and incubating at RT for 30 min in a humidified chamber. Hairpins h1 and h2 (2 μL each) were separately snap-cooled by heating to 95°C for 90 sec and then transferred to a dark drawer to cool to RT for 30 min. An amplifier solution was prepared by adding the snap-cooled h1 and h2 hairpins to 100 μL of Amplifier Buffer at RT. After removing the pre-amplification solution, samples were incubated with 100 μL of the amplifier solution at RT for 8 hr in a dark humidified chamber. Unbound amplifiers were removed by washing the samples four times (15 min per wash) with 5× SSCT at RT in the dark.

Following FISH, samples were blocked (1% BSA, 0.3% Triton X-100 in 1× PBS) at RT for 30 min and incubated overnight at 4°C with primary antibodies (Goat anti-PDGFRβ, R&D systems, Cat#AF385) diluted in the same blocking buffer. After three 5-min washes with 1× PBS, samples were incubated with fluorophore-conjugated secondary antibodies (Donkey anti-Goat 555, Invitrogen, Cat#A32816) and DAPI (Bioss, Cat#S0001) diluted in blocking buffer at RT for 1.5 h in the dark. Finally, samples were washed three times with 1× PBS and mounted with an antifade mounting medium (Fluoro-Gel, EMS, Cat#17985-11).

### Image acquisition and processing

Fixed brain, spinal cord, and cultured cell samples were imaged using a Leica TCS SP8 (Leica, Germany) or Nikon A1 (Nikon, Japan) laser scanning confocal microscope. Low-magnification images of brain and spinal cord sections were acquired using a 10×/0.40 numerical aperture (NA) objective on the Leica TCS SP8 and stitched using Leica LAS X software to generate whole-section images. Regions of interest were imaged using either a 63×/1.40 NA oil-immersion objective on the Leica TCS SP8 or a 60×/1.49 NA oil-immersion objective on the Nikon A1. Confocal z-stacks were acquired at 0.5- or 1-μm intervals. Maximum-intensity projections and orthogonal (XY, XZ, and YZ) views were generated using Fiji. 3D reconstructions and volumetric analyses were performed using Imaris (Bitplane). For each experiment, all samples were imaged using identical acquisition settings, including laser power, detector gain, and pinhole size. Brightness and contrast were adjusted uniformly across images. Gamma correction and smoothing were applied only when necessary to improve visualization and did not alter data interpretation.

### 3D reconstruction of confocal images

The acquired images of perivascular cells and myelin staining in fixed samples were preprocessed with Fiji. Brightness and contrast were adjusted, and region of interest was duplicated and saved for downstream processing in Imaris (version 9.9.0, Bitplane). 3D reconstruction of perivascular cells and myelin signals were created by the ‘Surface’ and ‘Mask’ tools in Imaris. The ‘Surface’ tool was applied to reconstruct perivascular cells by adjusting ‘Smooth’, ‘Threshold’ and ‘Filter’ parameters following the creation wizard. The ‘Smooth’ parameter of perivascular cell surface was set to 0.3μm for ‘Surface Detail’. The ‘Threshold’ parameter was set by adjusting absolute intensity. ‘Filter Surface’ was applied to eliminate discontinuous signals of the perivascular cell surface. This cell surface was pseudo-colored in magenta. The recreated perivascular cell surface then served as the region of interest to generate a mask interface of myelin channel where MBP signals were only present within perivascular cells. The ‘Mask’ tool was used to create a mask interface for the myelin channel, which was merged with perivascular cell surface to generate 3D-reconstructed images. The ‘Surface Detail’ of MBP surface was adjusted to 0.1 and colored in green. The same settings were employed to create perivascular cell surfaces and ingested myelin interfaces. The 3D-constructed images were saved using Snapshot option. Videos showing volumetric view of perivascular cell phagocytosis of myelin debris were created using Animation feature in Imaris, with keyframes introduced at various time intervals. Videos were compressed via Microsoft Powerpoint and brief descriptions were added in the videos via CapCut video editor software (version 7.1.0) without modifying scientific contents.

### RT-qPCR

RNA was extracted using TRIzol (ThermoFisher, Cat#15596026). cDNA was reverse transcribed using the Toyobo ReverTra Ace qPCR RT Kit (TOYOBO, Cat#FSQ-101) according to manufacturer’s instructions. qPCRs were performed in 10μL final volumes containing 5μL of DNA master mix for SYBR Green Master Mix (Yeasen, Cat#11201ES08), 0.5 μL of each primer, and 1 μL of template DNA. All samples were amplified by Bio-Rad CFX96. The specificity of every reaction was determined using melting curve analysis. The expression level of target genes was normalized to *GAPDH* and calculated using the ΔΔ^Ct^ method. For each sample, independent repeats were performed in triplicate.

### Western blot

Cultured perivascular cells were lysed in cold RIPA buffer containing 1% protease inhibitor cocktail. Protein concentrations in the cleared lysates were measured by bicinchoninic acid (BCA) test. 40 μg of total protein were resolved by 12% SDS-PAGE gel, which was then transferred to 0.22 μm PVDF membranes using a wet transfer system (Bio-Rad). After blocking with 5% BSA in 1× TBS containing 0.1% Tween for 30 min, the membranes were incubated overnight at 4 °C diluted in TBST. After washing with TBST three times for 5 min each, the membranes were incubated with HRP-conjugated secondary antibodies for 30 min at room temperature. The membranes were developed using an ECL kit, and the resulting images were processed with Fiji.

### Quantitative and statistical analysis

For quantitative analysis of myelin debris phagocytosis in vivo, confocal z-stacks were reconstructed in three dimensions using Imaris (v9.9.0). The total volume of reconstructed PDGFRβ^+^ perivascular structures and the volume of intracellular MBP signal within the corresponding perivascular mask (PDGFRβ^+^or GLAST^+^) were quantified using the Imaris statistics module. Phagocytic volume was expressed as the percentage of the engulfed MBP volume (MBP^+^ PDGFRβ^+^volume or MBP^+^ GLAST^+^ volume) relative to the total perivascular volume (PDGFRβ^+^ or GLAST^+^ volume).

Because perivascular cells undergo extensive proliferation and form densely interconnected networks after injury, reliable segmentation of individual cells was not feasible. Therefore, the frequency of phagocytic perivascular cells was estimated by quantifying individual vascular unit (PDGFRβ^+^ or GLAST^+^) on single optical sections, where adjacent vascular unit could be readily distinguished. A PDGFRβ^+^ or GLAST^+^ vascular unit was scored as phagocytic when it contained clearly identifiable intracellular MBP puncta. The percentage of phagocytic events was calculated as the number of MBP^+^ PDGFRβ^+^ (or MBP^+^ GLAST^+^) vascular units divided by the total number of PDGFRβ^+^ (or GLAST^+^) vascular units analyzed.

For quantification of *in vitro* perivascular cell phagocytosis of CFSE-labeled cargos (myelin debris, neuronal bodies and zymosan), the area of fluorescent CFSE signals in 20× or 63× confocal images was measured in Fiji. Each image was first converted to 811bit RGB format and inverted to grayscale. A threshold was then applied to establish identical fluorescence settings across all groups. The fluorescent area was determined by measuring the area of CFSE signal within CFSE⁺ αSMA⁺ cells and then normalized to the total number of αSMA⁺ cells (based on DAPI staining). The percentage of cargo-laden cells were determined by calculating CFSE^+^ αSMA^+^ cells over total αSMA^+^ cells. For area measurements of other markers (fibronectin, collagen I, ORO, Ctsb), the fluorescent areas of interested signals in perivascular cells were measured. The area of interested signal within αSMA⁺ cells was divided by the total αSMA⁺ cell area.

The following markers were quantified as the area of fluorescent signals in spinal cord or brain sections: perivascular cells (PDGFRβ-tdTomato or PDGFRβ), myelin (MBP), fibrosis (fibronectin and collagen I), axons (NF-H and 5-HT), astrocytes (GFAP), and myeloid cells (IBA1 or F4/80). Stitched low-magnification confocal images (10× objective) encompassing the lesion were analyzed using Fiji (ImageJ). To ensure consistent sampling across animals, a fixed square region of interest (ROI) centered on the lesion was applied to all sections. At the analyzed time points, the lesion area was smaller than the ROI, allowing the entire lesion to be included within the same measurement window for all samples. Images were converted to 8-bit format, and a uniform fluorescence threshold was applied identically to all images within each experiment. The area occupied by the fluorescence signal above threshold within the ROI was then measured and expressed as the fluorescent area. This analysis quantified the area occupied by marker-positive signals.

The size of lysosomes in perivascular cells was quantified by measuring the diameter of LAMP1^+^ puncta. In myelin debris-treated perivascular cells, the sizes of lysosomes with or without myelin debris were separately quantified. The percentage of LC3^+^ phagosomes were determined by calculating the number of LC3^+^ LAMP1^+^ puncta over total LAMP1^+^ puncta in fixed perivascular cells with or without cargos. The number of single-membrane vesicles that contain multiple membrane structures (myelin) was manually counted in every cell from TEM images.

For co-localization analysis, confocal images were split into single-channel images based on antibody combination (e.g., CD31 and ZO-1 or Mfsd2a; myelin debris and Ctsb) and converted to RGB format using Fiji software. Subsequently, images were inverted to grayscale. The same fluorescence threshold was applied to both channels. The Image Calculator function was used to generate colocalized images from the two channels, followed by measuring the colocalization area using the same thresholding criteria.

For the quantification of BBB leakage, confocal images acquired at 10× magnification were converted to RGB format and inverted to grayscale in Fiji. After thresholding, the total fluorescence area of the entire field was quantified, and the fluorescence area within the vascular ROIs was subtracted from the total area to calculate the extravascular Sulfo-NHS-Biotin leakage signal.

To determine the cellular distribution of Axl, confocal images acquired at 20× magnification were used to identify Axl-positive cells, marker-positive populations and double-positive cells (Axl-positive within each cell type-specific marker population). The proportion of Axl-expressing cells within each cell type was calculated as the number of Axl+/marker+ double-positive cells normalized to the total number of Axl+ cells.

To evaluate the labeling efficiency and specificity of PDGFRβ-Cre::Ai14 reporter mice in the spinal cord, tissue sections from mice at P21, P45, and P90 were stained for mCherry, PDGFRβ, CD13, IBA1, GFAP, and NeuN, followed by analysis of confocal images acquired at 20× magnification. For labeling efficiency, the number of mCherry+/CD13+ double positive cells was normalized to the total number of CD13+ cells. Labeling specificity was quantified by normalizing the number of cells double-positive for mCherry and a given lineage marker to the total number of mCherry+ cells.

*In vitro* Ki67^+^ proliferating perivascular cells were counted, and the proliferation index was calculated by the proportion of Ki67^+^ cells to the total number of cells (indicated by DAPI staining) and normalized by fold change. For quantification of *in vivo* Ki67^+^ proliferative perivascular cell in Matrigel plug assay, myelin microinjection assay and SCI model, the number of K67^+^ PDGFRβ^+^ cells was counted and the area of PDGFRβ^+^ was measured. A proliferation index was calculating the density of Ki67^+^ PDGFRβ^+^ cells in area covered by total PDGFRβ^+^cells.

For *in vivo* proliferation analyses in the Matrigel plug assay, myelin microinjection assay, and SCI model, Ki67^+^ PDGFRβ^+^ cells were counted and normalized to the total PDGFRβ^+^ area. The proliferation index was calculated as the density of Ki67^+^ PDGFRβ^+^ cells per unit PDGFRβ^+^ area.

For quantification of p-Smad3 activation, the density of p-Smad3^+^ PDGFRβ^+^ cells in tissue sections was calculated as the number of double-positive cells per PDGFRβ^+^ area. In cultured perivascular cells, p-Smad3 activation was quantified as the percentage of cells displaying nuclear p-Smad3 staining among the total number of cells identified by DAPI staining.

For quantification of in situ mRNA hybridization, Axl mRNA signals were thresholded in Fiji, and the area of Axl mRNA puncta within PDGFRβ^+^ perivascular cells, SOX9^+^ astrocytes, IBA1^+^ microglia, NeuN^+^ neurons, and CC1^+^ oligodendrocytes was measured. For comparisons among different cell types, Axl mRNA signal was normalized to the corresponding cell number. For comparisons of perivascular cells before and after SCI, Axl mRNA signal was normalized to the total PDGFRβ^+^ area.

Statistical analyses were performed using GraphPad Prism 9.5.1 (GraphPad Software, San Diego, CA). The statistical test used for each experiment is specified in the corresponding figure legend. Depending on the experimental design, comparisons were performed using unpaired two-tailed Student’s *t* test, one-way ANOVA, or two-way ANOVA followed by appropriate post hoc multiple-comparison tests. Data are presented as mean ± SEM from at least three independent biological replicates or independent experiments. Differences were considered statistically significant at *P* < 0.05.

## Author contributions

TZ, LW, YR and YZ conceived and designed the research study. TZ and YZ performed experiments and analyzed data. ZZ, FZ, HS, ZX, ZH, LY, WW, TZ, XD, KL, ZX, YS, BR, BF, SQ, YL, YH, MF, YC, NZ, MA contributed to the experiments and data analysis. NH, HS, LS, KH, QQ, QL, FP, XC, CSZ, KM, BW, ZJ, GB, FM and TM provided essential reagents, provided scientific advice, and contributed to the experiments/anlaysis. TZ, YR and YZ wrote the manuscript with the inputs from all authors. TM and FP critically edited the manuscript.

## Declaration of generative AI and AI-assisted technologies in the writing process

During the preparation of this work, the authors used ChatGPT in order to refine language. After using this tool, the authors reviewed and edited the content as needed and take full responsibility for the content of the published article.

## Acknowledgements

We thank Prof. Daijun Han (Peking University), Prof. Xiaojun Xu (China Pharmaceutical University), Prof. Linjuan Zhang (Xiamen University) and Prof. Jie Zhang (Xiamen University) for kindly providing *Axl* KO mice, Prof. Xingqun Liang (Tongji University) for kindly providing PDGFRβ-Cre mice, Prof. Bayasi Guleng (Xiamen University) for kindly providing *CR3* KO mice, Prof. Chengyong He (Xiamen University) for kindly providing p-Smad3 antibody, Prof. Tianzhi Huang (Xiamen University) for kindly sharing plasmids. We thank Prof. Chao Ma and Ms. Xue Wang (National Human Brain Bank for Development and Function), Dr. Huachen Huang (Tianjin Medical University General Hospital) for assistance with human stroke samples. We thank Prof. Zengqiang Yuan (Institute of Biophysics, Chinese Academy of Sciences) for feedback on the work. We thank Dr. Marisa Tillery for figure illustration. We thank Caiming Wu, Lei Huang and Qingfen Liu at Core Facility of Biomedical Sciences of Xiamen University for technical support, Yunmei Wu and Bingbing Fan at Research Service Center of School of Life Sciences in Xiamen University for assisting with laboratory management, Jianfeng Wu, Xuejuan He and Ying He in Xiamen University Laboratory Animal Center for assisting with animal care. We thank Shuji Li, Ting Guo, Yingying Fang and Wen Yao at Core Facility of Southern Medical University for technical support. This work was supported by Shenzhen Medical Research Funds (A2303068 to TZ), the National Natural Science Foundation of China (32370731 to YZ, 32471046 to TZ, 82171082 to ZX, 32170902 to ZJ), Shenzhen Science and Technology Program (JCYJ20230807091308018 to YZ, JCYJ20220531100204010 and GJHZ20220913142807015 to TZ, JCYJ20210324101603009 and JCYJ20240813155017022 to ZX), Science and Technology Program of Guangzhou (2025A04J4473 to TZ), Major Non-consensus Program of Guangdong Provincial Basic and Applied Basic Research Program (2026B0303050006 to TZ). Guang Dong Basic and Applied Basic Research Foundation (2024A1515010691 to ZX), The Fundamental Research Funds for the Central Universities (20720250100 to YZ), National Science Foundation (DMS-2054014 to YR), The Shenzhen Medical Research Fund (C2406001 to LW), Research Grants Council of Hong Kong (the Theme-Based Research Scheme T12-611/25-N to GB).

---

**Figure S1 – related to Figure 1.**
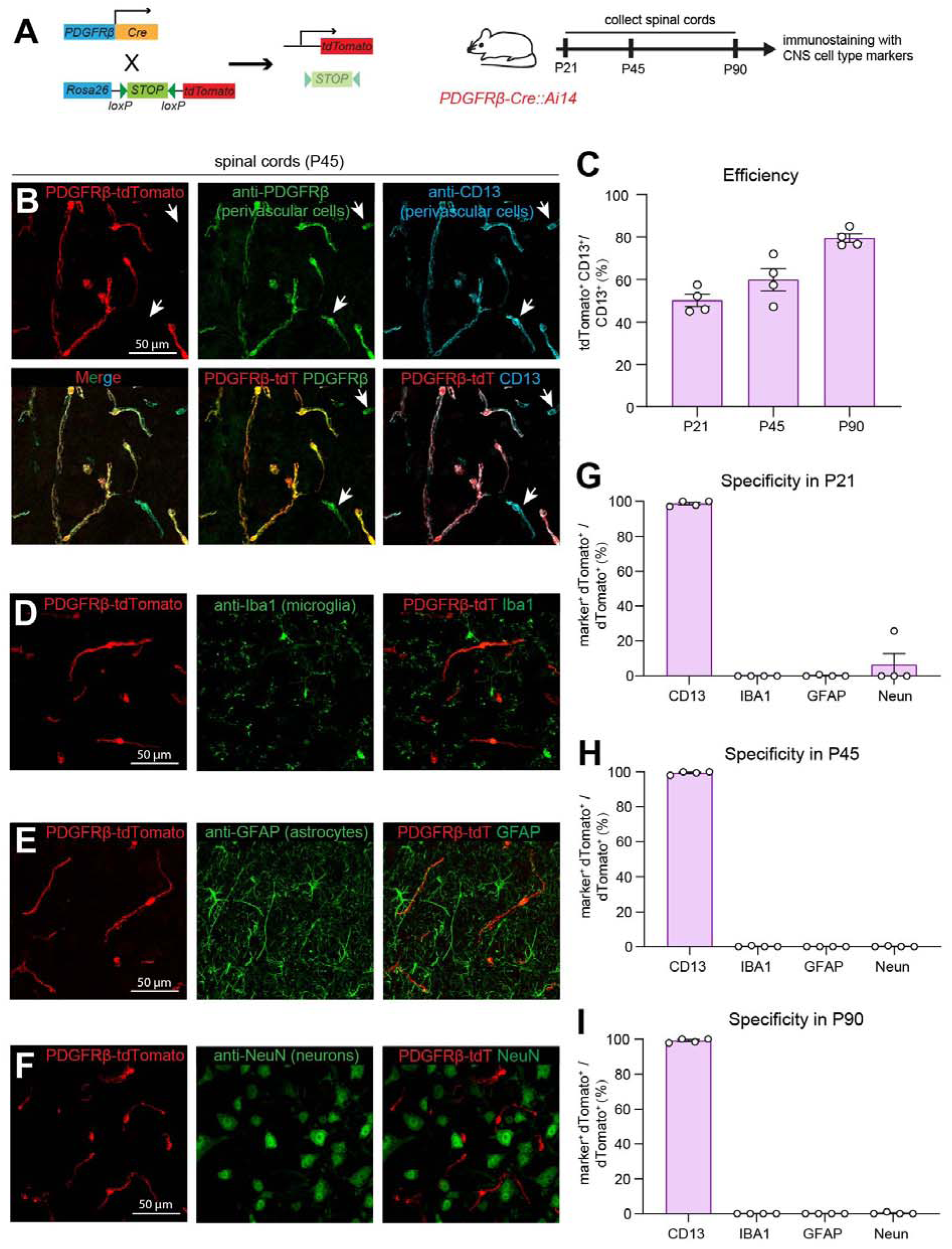
Validation of specificity and efficiency of the PDGFRβ-Cre strain in labeling perivascular cells of normal spinal cords across time points. **(A)** Experimental approaches to genetically label perivascular cells in the spinal cords using PDGFRβ-Cre-mediated recombination of R26R-loxP-STOP-loxP-tdTomato (Ai14) reporter strain (left) and to characterize labeling efficacy and specificity at postnatal days 21, 45, and 90 using immunohistochemical staining (right). **(B)** Representative images of P45 PDGFRβ-Cre::Ai14 spinal cords immunostained for the perivascular cell markers PDGFRβ (green) and CD13 (white). tdTomato (red) marks PDGFRβ-Cre-labeled cells. Arrows indicate PDGFRβ^⁺^CD13^⁺^ perivascular cells that were not labeled by the PDGFRβ-Cre::Ai14 reporter. Scale bar, 50 μm. **(C)** Quantification of the labeling efficiency of PDGFRβ-Cre::Ai14 in spinal cords at P21, P45, and P90, calculated as the percentage of tdTomato^⁺^CD13^⁺^ cells among total CD13^⁺^ perivascular cells. Data are presented as mean ± SEM (n = 4 mice). The total number of cells analyzed at each time point is indicated. **(D–F)** Representative confocal images showing the labeling specificity of PDGFRβ-Cre::Ai14 in P45 spinal cords by co-immunostaining with markers for microglia (Iba1, **D**), astrocytes (GFAP, **E**), and neurons (NeuN, **F**). Scale bar, 50 μm. **(G–I)** Quantification of the labeling specificity of PDGFRβ-Cre::Ai14 in spinal cords at P21, P45, and P90, calculated as the percentage of tdTomato⁺ cells co-expressing the indicated cell-type marker among all tdTomato^⁺^ cells. Data are presented as mean ± SEM (n = 4 mice). The total number of cells analyzed for each cell type at each time point is indicated.

**Figure S2 – Related to Figure 1.**
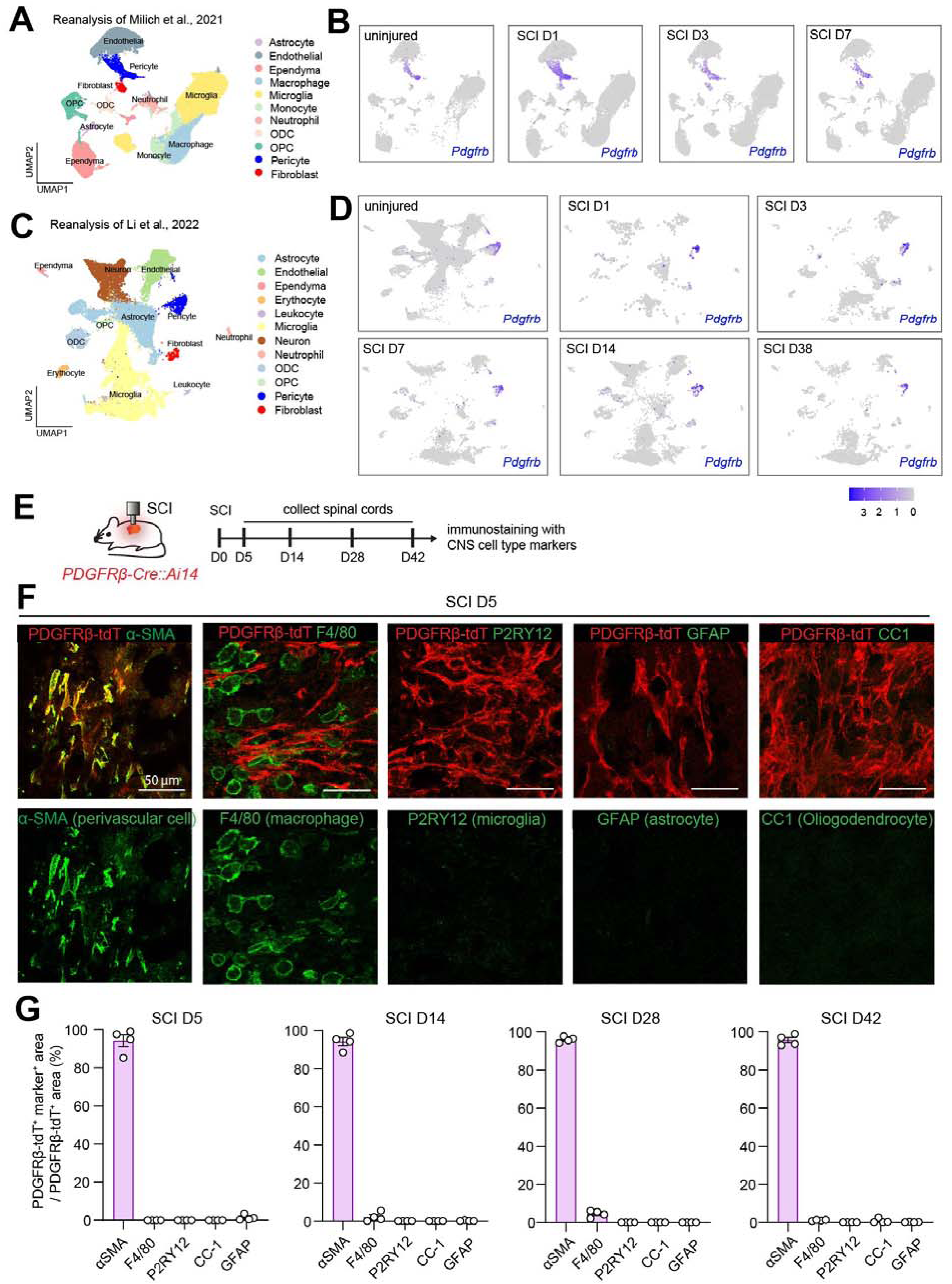
Validation of PDGFRβ-Cre specificity for labeling perivascular cells after spinal cord injury. **(A, B)** Reanalysis of the single-cell RNA-sequencing dataset from Milich *et al.* (2021) showing the cellular distribution of *Pdgfrb* expression in spinal cords at 1, 3, and 7 days after SCI. *Pdgfrb* expression remains largely restricted to perivascular cell clusters (pericytes and fibroblasts), with minimal expression in other CNS cell types. **(C, D)** Reanalysis of the single-cell RNA-sequencing dataset from Li *et al.* (2022) showing the cellular distribution of *Pdgfrb* expression in spinal cords at 4 hr, 1, 3, 7, 14, and 38 days after SCI. *Pdgfrb* expression remains predominantly restricted to perivascular cell clusters throughout SCI progression. **(E)** Experimental design for evaluating the specificity of PDGFRβ-Cre::Ai14-mediated labeling in spinal cords collected at the indicated time points after SCI. **(F)** Representative confocal images of spinal cord sections from PDGFRβ-Cre::Ai14 mice at 5 days after SCI showing tdTomato-labeled cells (red) together with immunostaining for α-SMA (perivascular cells), F4/80 (macrophages), P2RY12 (microglia), GFAP (astrocytes), and CC1 (oligodendrocytes). Scale bar, 50 μm. **(G)** Quantification of the labeling specificity of PDGFRβ-Cre::Ai14 at the indicated time points after SCI, expressed as the percentage of tdTomato^⁺^ cells co-expressing the indicated cell-type markers. Data are presented as mean ± SEM (n = 4 mice per time point).

**Figure S3 – Related to Figure 1.**
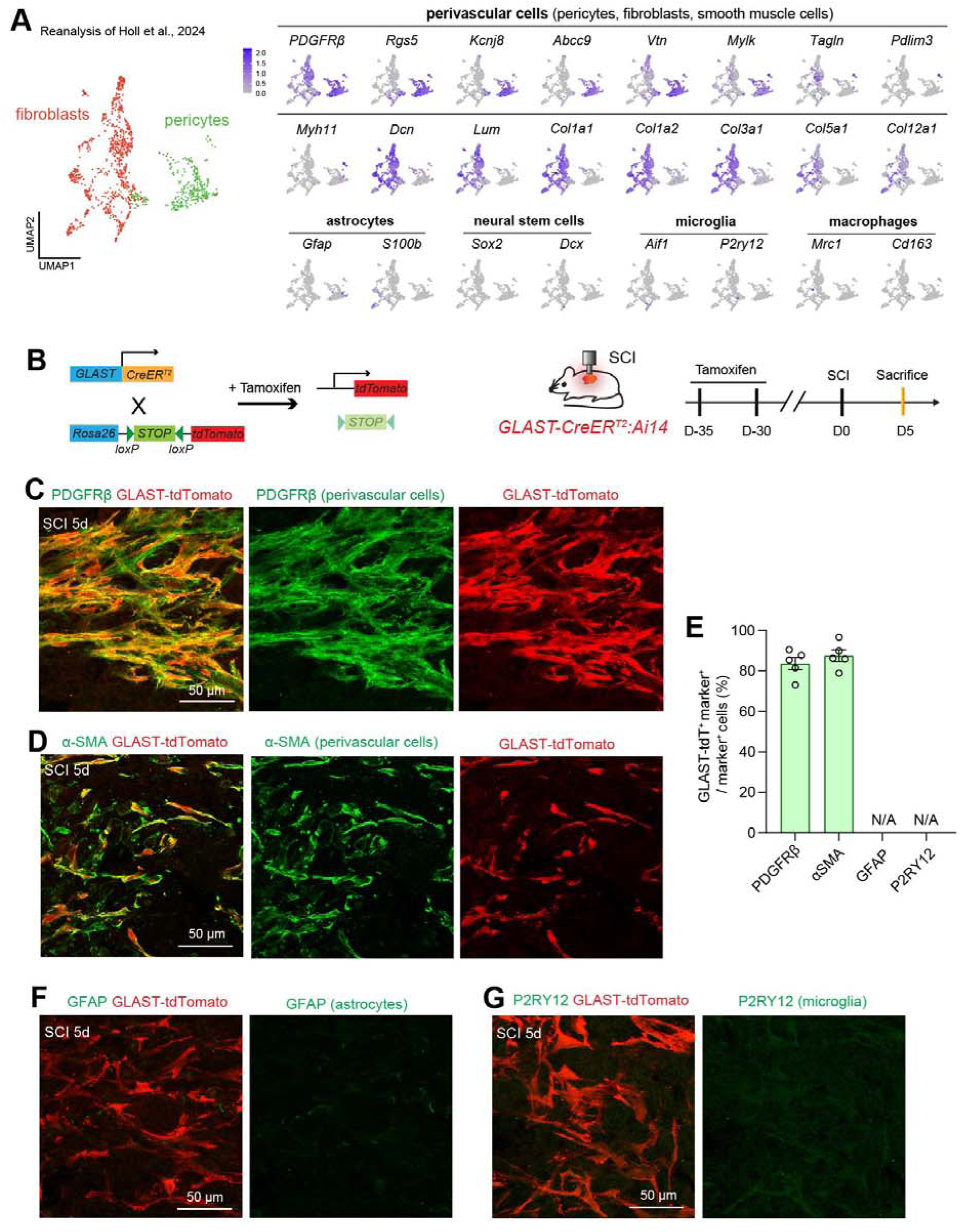
Validation of the GLAST-CreERT2 line for labeling perivascular cells after spinal cord injury. **(A)** Reanalysis of the single-cell RNA sequencing dataset from Holl *et al.*, 2024, showing the molecular identity of GLAST-expressing cells after SCI. Following injury, GLAST-expressing cells are enriched in perivascular cell populations, including pericytes, perivascular fibroblasts, and vascular smooth muscle cells, with minimal expression of markers for other CNS cell types. **(B)** Schematic illustrating genetic labeling of perivascular cells in GLAST-CreERT2::Ai14 mice. Tamoxifen-induced Cre recombination activates tdTomato expression from the Ai14 reporter. The timeline indicates tamoxifen administration, SCI, and spinal cord collection for immunohistochemical analysis. **(C, D)** Representative confocal images showing colocalization of tdTomato-labeled cells with the perivascular cell markers PDGFRβ **(C)** and α-SMA **(D)** in spinal cord sections at 5 days after SCI. Scale bar, 50 μm. **(E)** Quantification of the percentage of PDGFRβ^⁺^ or α-SMA^⁺^ perivascular cells labeled by tdTomato in GLAST-CreERT2::Ai14 mice at 5 days after SCI. Data are presented as mean ± SEM (n = 5 mice). **(F, G)** Representative confocal images showing minimal colocalization of tdTomato-labeled cells with the astrocyte marker GFAP **(F)** and the microglia marker P2RY12 **(G)** in spinal cord sections at 5 days after SCI. Scale bar, 50 μm.

**Figure S4 – Related to Figure 1.**
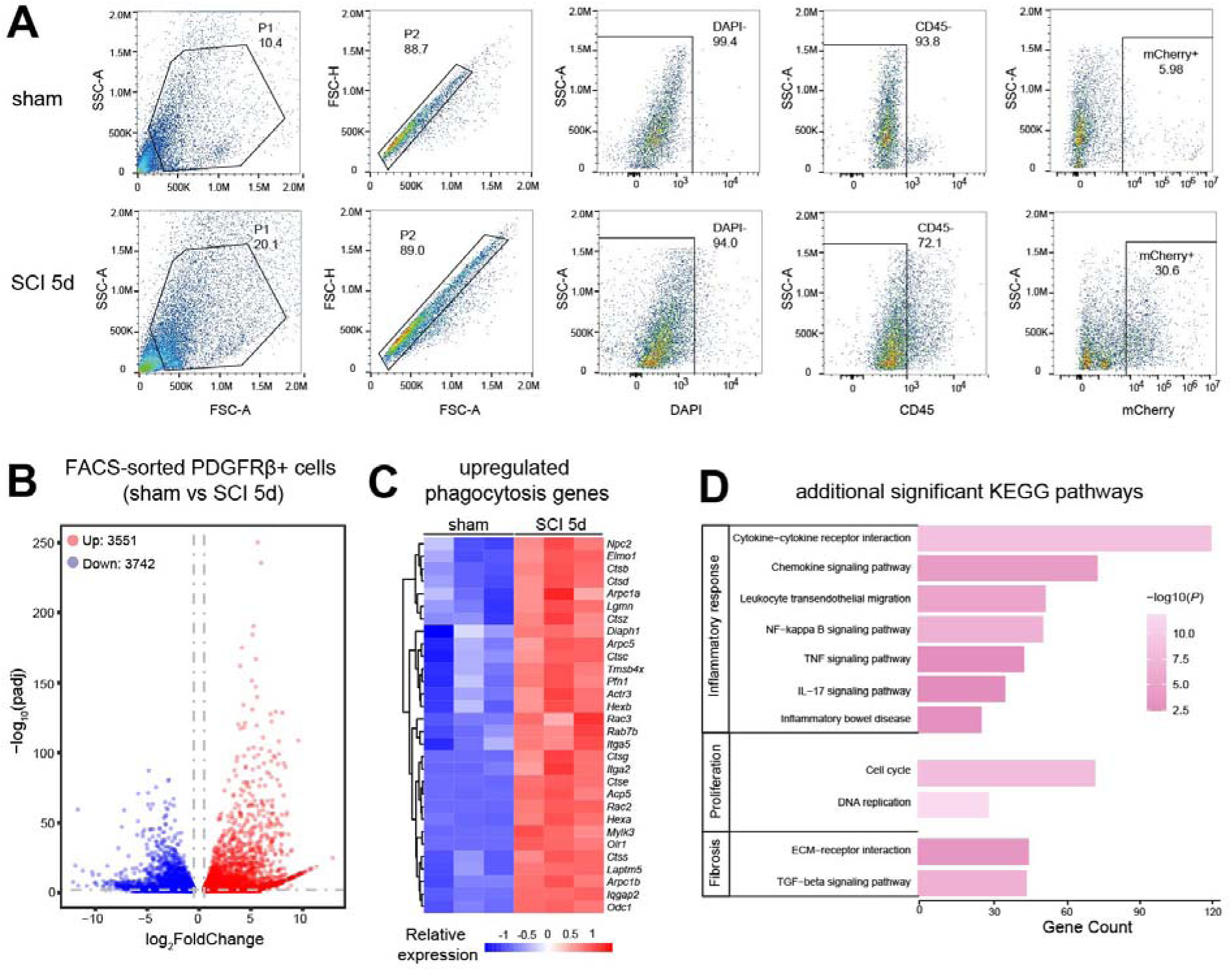
Transcriptomic profiling of purified perivascular cells following spinal cord injury. **(A)** Gating strategy for fluorescence-activated cell sorting (FACS) of perivascular cells (PDGFRβ-tdTomato^⁺^ CD45^⁻^) from sham-operated and SCI spinal cords. **(B)** Volcano plot showing differentially expressed genes (DEGs) in purified perivascular cells from sham-operated and SCI spinal cords. Genes with |log_2_FoldChange | > 0.5 and a false discovery rate (FDR)-adjusted *P* < 0.05 were considered significantly differentially expressed. **(C)** Heatmap showing the relative expression of phagocytosis-related genes that were significantly upregulated in purified perivascular cells following SCI compared with sham controls. **(D)** Additional KEGG pathways significantly enriched in purified perivascular cells following SCI.

**Figure S5 – Related to Figure 1.**
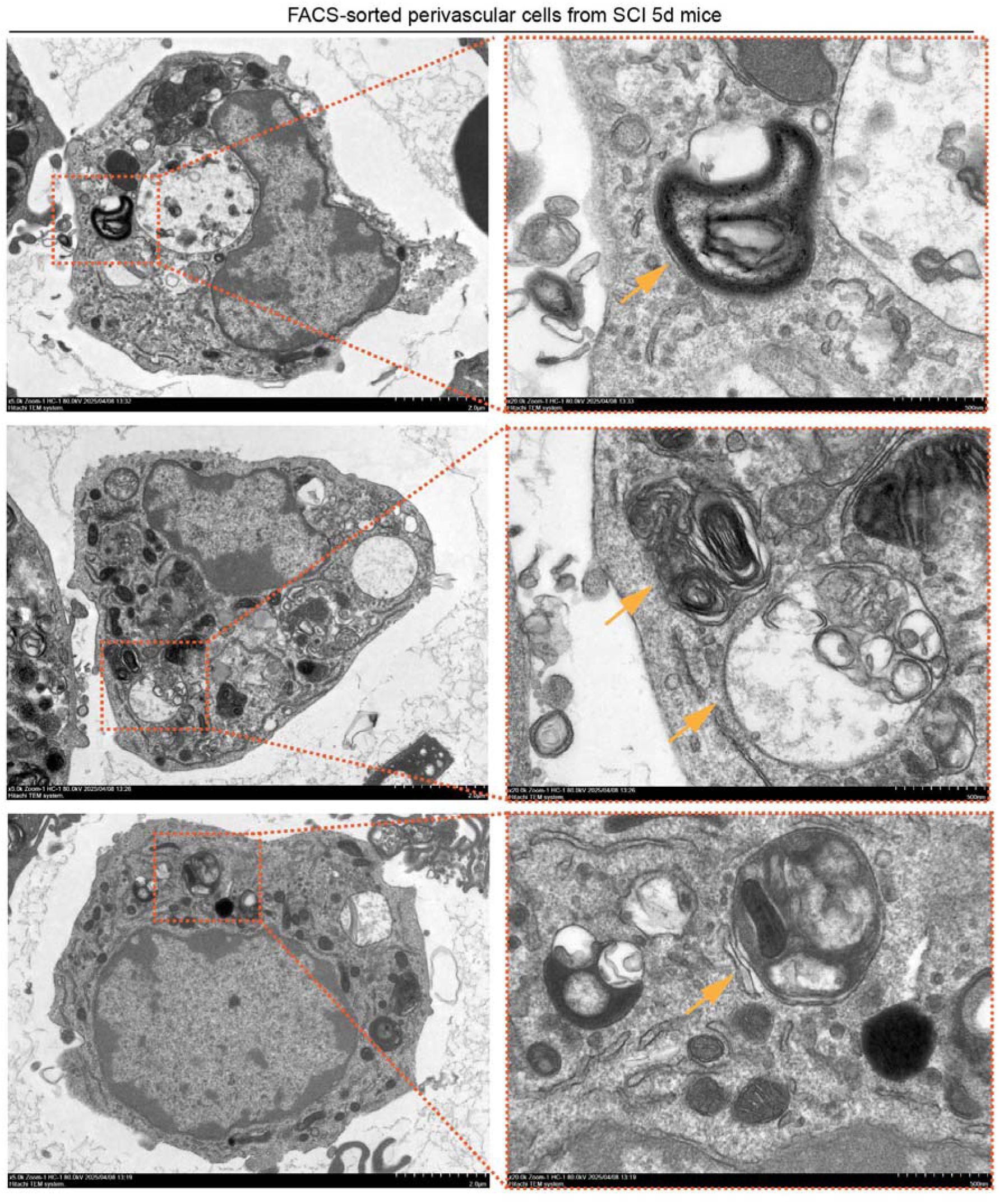
Ultrastructural evidence of myelin debris within acutely isolated perivascular cells after spinal cord injury. Transmission electron micrographs showing multilamellar myelin structures (arrows) within FACS-purified perivascular cells isolated from spinal cords at 5 days after SCI. Scale bars, 2 µm and 500 nm (enlarged view).

**Figure S6 – Related to Figure 1.**
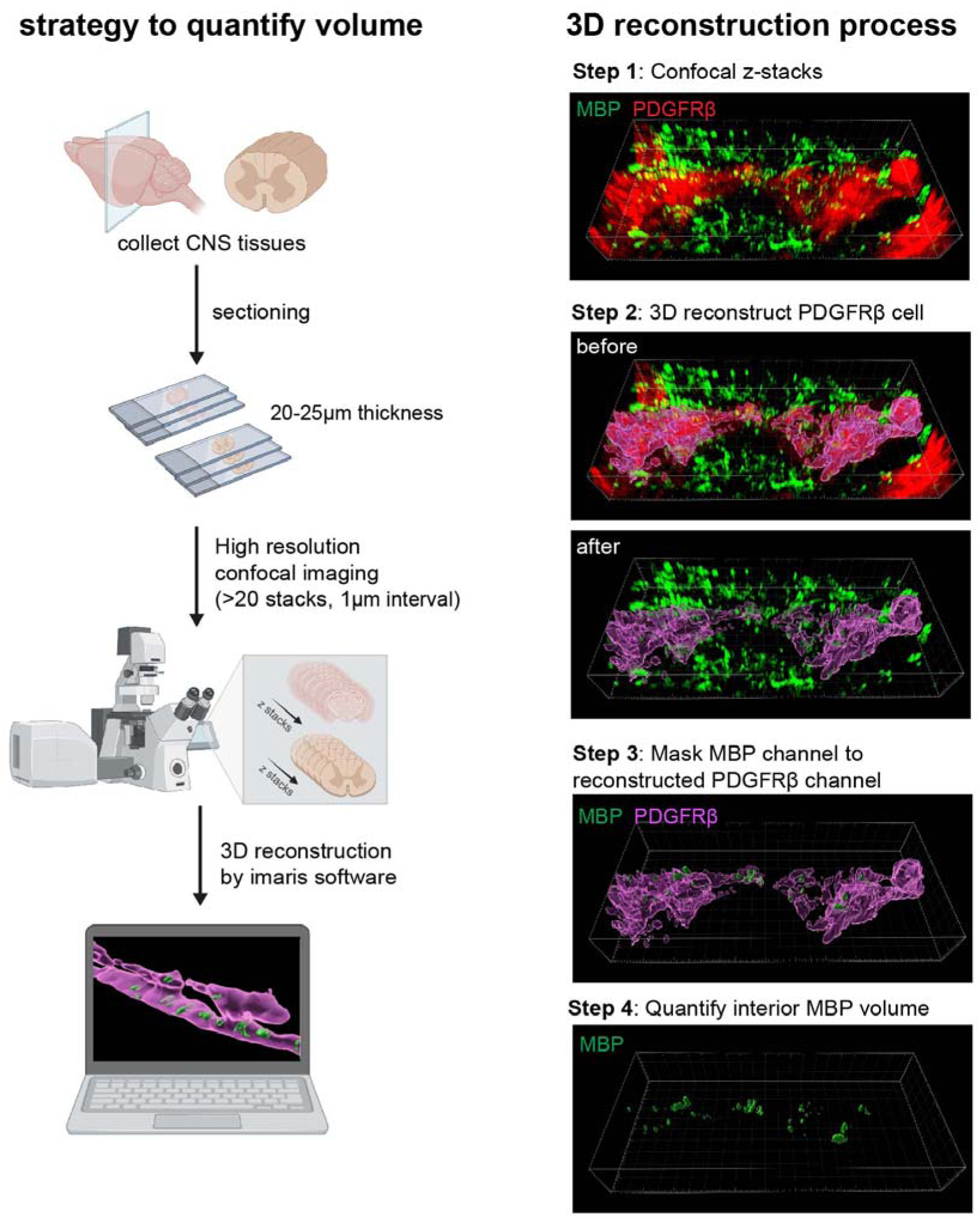
Strategy for quantifying the volume of engulfed cargos by 3D reconstruction. Left, schematic illustrating the workflow for volumetric quantification of engulfed cargos. Mouse spinal cords or brains were sectioned, immunostained, and imaged by confocal microscopy to acquire high-resolution z-stacks for 3D reconstruction in Imaris. Right, representative images illustrating the stepwise reconstruction of engulfed cargos (e.g., myelin debris) within PDGFRβ^⁺^ perivascular cells from confocal z-stacks for volumetric analysis.

**Figure S7 – Related to Figure 2.**
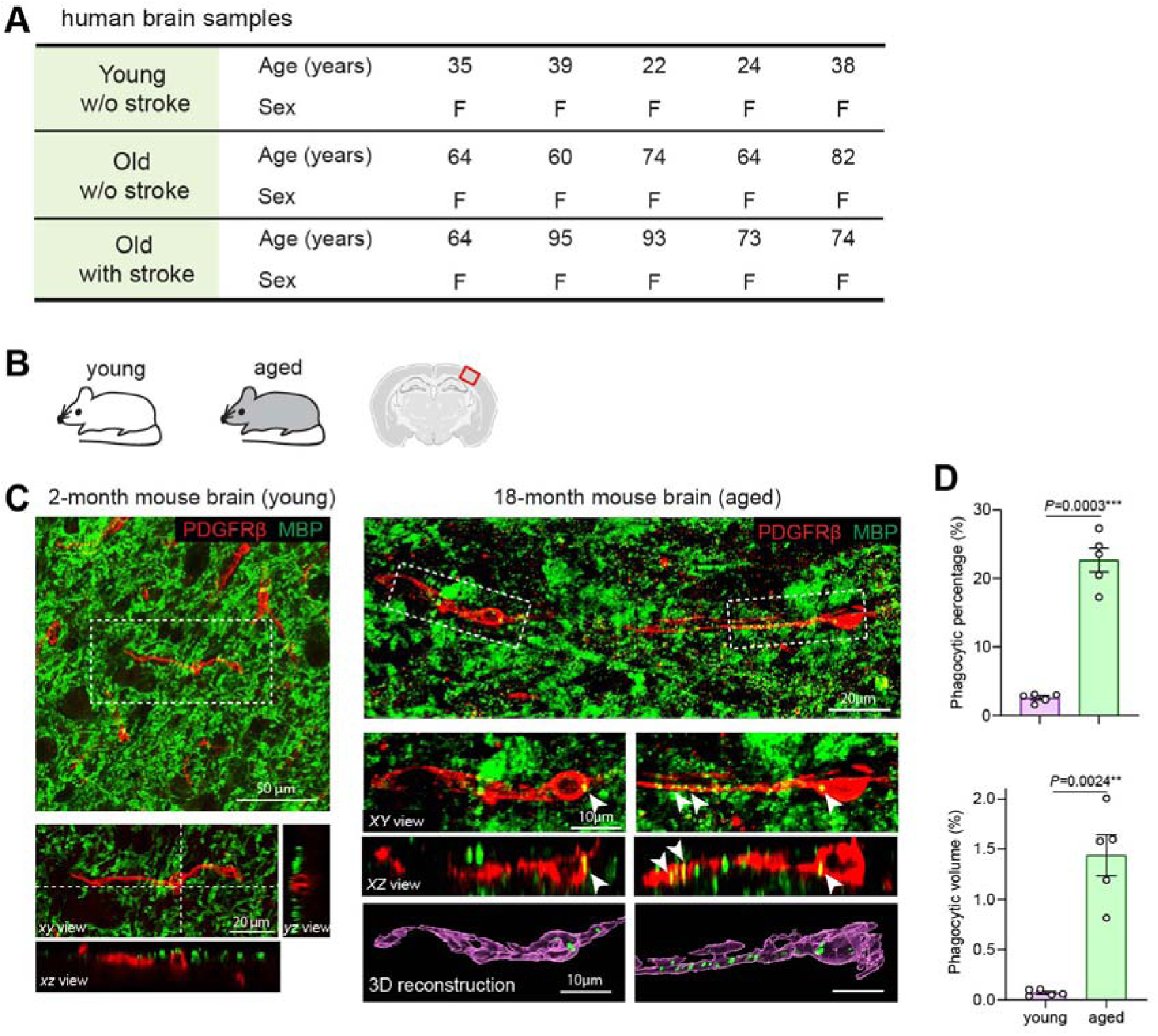
Perivascular cells engulf myelin debris during aging in humans and mice. **(A)** Table summarizing the clinical information of human brain samples. **(B)** Schematic illustrating the cortical region analyzed in young and aged mouse brains. **(C)** Representative confocal images of PDGFRβ^⁺^ perivascular cells (red) containing MBP^⁺^ myelin debris (green) in the brains of aged (18-month-old) mice. Higher-magnification images show orthogonal XZ views and 3D reconstructions demonstrating intracellular localization of MBP^⁺^ myelin debris. Scale bars: 50 μm (left, top), 20 μm (left, XY view), 20 μm (right, top), and 10 μm (right, XY view and 3D reconstruction). **(D)** Quantification of the phagocytic percentage (percentage of PDGFRβ^⁺^ perivascular cells containing MBP^⁺^ myelin debris) and phagocytic volume (percentage of intracellular MBP^⁺^ volume relative to total PDGFRβ^⁺^ cell volume). Data are presented as mean ± SEM (n = 5 mice). Statistical significance was determined by unpaired two-tailed *t* test with Welch’s correction.

**Figure S8 – Related to Figure 2.**
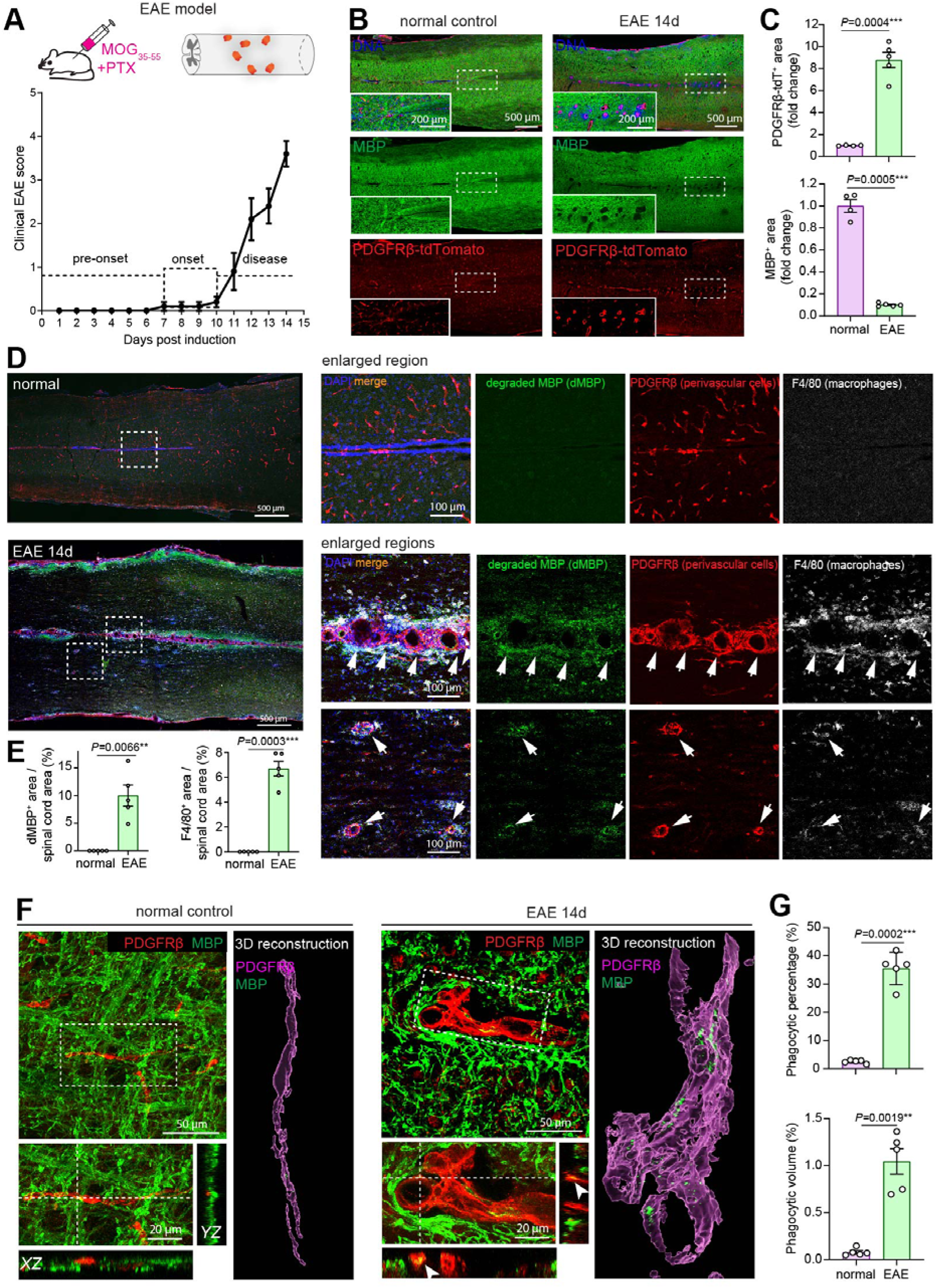
Perivascular cells expand in demyelinating lesions and function as phagocytes in the mouse EAE model of multiple sclerosis. **(A)** Experimental design of the experimental autoimmune encephalomyelitis (EAE) model induced by immunization with myelin oligodendrocyte glycoprotein peptide (MOG_35-55_). Demyelinating lesions in the spinal cord are indicated in orange. Clinical EAE scores showing disease progression through the three disease stages are presented as mean ± SEM. **(B)** Representative confocal images of spinal cord sections from control and EAE mice at 14 days after immunization, showing PDGFRβ-tdTomato^⁺^ perivascular cells (red) and MBP^⁺^ myelin (green). Insets show higher-magnification views of the boxed regions, illustrating expansion of PDGFRβ-tdTomato^⁺^ perivascular cells and loss of MBP immunoreactivity within demyelinating lesions. Scale bars: 500 μm; 200 μm (insets). **(C)** Quantification of PDGFRβ-tdTomato⁺ perivascular cell area and MBP immunoreactive area in control and EAE spinal cords. Data are presented as mean ± SEM (n = 5 mice). Statistical significance was determined by unpaired two-tailed *t* test with Welch’s correction. **(D)** Representative confocal images showing demyelinated lesions (degraded MBP staining), accumulation of PDGFRβ^⁺^ perivascular cells, and infiltration of F4/80⁺ macrophages in spinal cords 14 days after EAE induction. Higher-magnification images show PDGFRβ^⁺^ perivascular cells and F4/80^⁺^ macrophages within demyelinated lesions (arrowheads). Scale bars: 500 μm; 100 μm (enlarged views). **(E)** Quantification of the relative area of degraded MBP (dMBP) immunoreactivity and F4/80^⁺^ macrophage area in control and EAE spinal cords. Data are presented as mean ± SEM (n = 5 mice). Statistical significance was determined by unpaired two-tailed *t* test with Welch’s correction. **(F)** Representative confocal images of PDGFRβ^⁺^ perivascular cells (red) and MBP^⁺^ myelin (green) in spinal cords 14 days after EAE induction. Higher-magnification XY, XZ, and YZ orthogonal views together with 3D reconstructions demonstrate intracellular MBP^⁺^ myelin debris within PDGFRβ^⁺^ perivascular cells (arrowheads). Scale bars: 50 μm; 20 μm (enlarged views). **(G)** Quantification of the phagocytic percentage (percentage of PDGFRβ^⁺^ perivascular cells containing MBP^⁺^ myelin debris) and phagocytic volume (percentage of intracellular MBP^⁺^ volume relative to total PDGFRβ^⁺^ cell volume). Data are presented as mean ± SEM (n = 5 mice). Statistical significance was determined by unpaired two-tailed *t* test with Welch’s correction.

**Figure S9 – Related to Figure 2.**
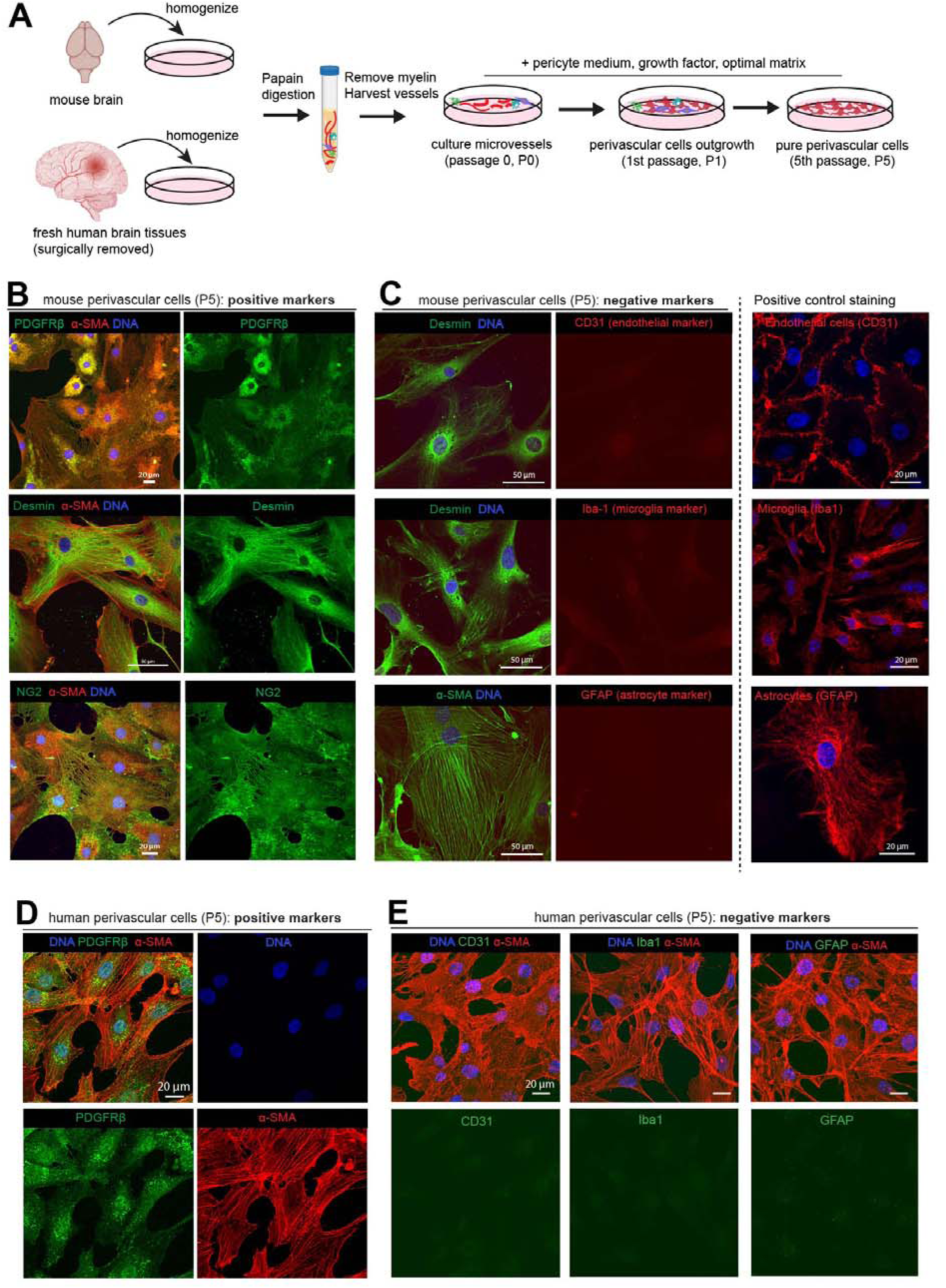
Isolation and characterization of primary mouse and human brain perivascular cells *in vitro*. **(A)** Workflow for isolation and purification of primary mouse and human brain perivascular cells. **(B)** Representative immunocytochemical images of mouse brain perivascular cells at passage 5 (P5) showing expression of the perivascular cell markers PDGFRβ, α-SMA, Desmin, and NG2. Scale bars: 20 μm (PDGFRβ, NG2); 50 μm (α-SMA, Desmin). **(C)** Representative immunocytochemical images of mouse brain perivascular cells at P5 showing absence of the endothelial marker CD31, microglial marker Iba-1, and astrocyte marker GFAP. Positive staining in the corresponding cell types is shown as controls (right panels). Scale bars: 50 μm (left panels); 20 μm (right panels). **(D)** Representative immunocytochemical images of human brain perivascular cells at P5 showing expression of the perivascular cell markers PDGFRβ and α-SMA. Scale bar: 20 μm. **(E)** Representative immunocytochemical images of human brain perivascular cells at P5 showing absence of the endothelial marker CD31, microglial marker Iba-1, and astrocyte marker GFAP. Scale bar: 20 μm.

**Figure S10 – Related to Figure 2.**
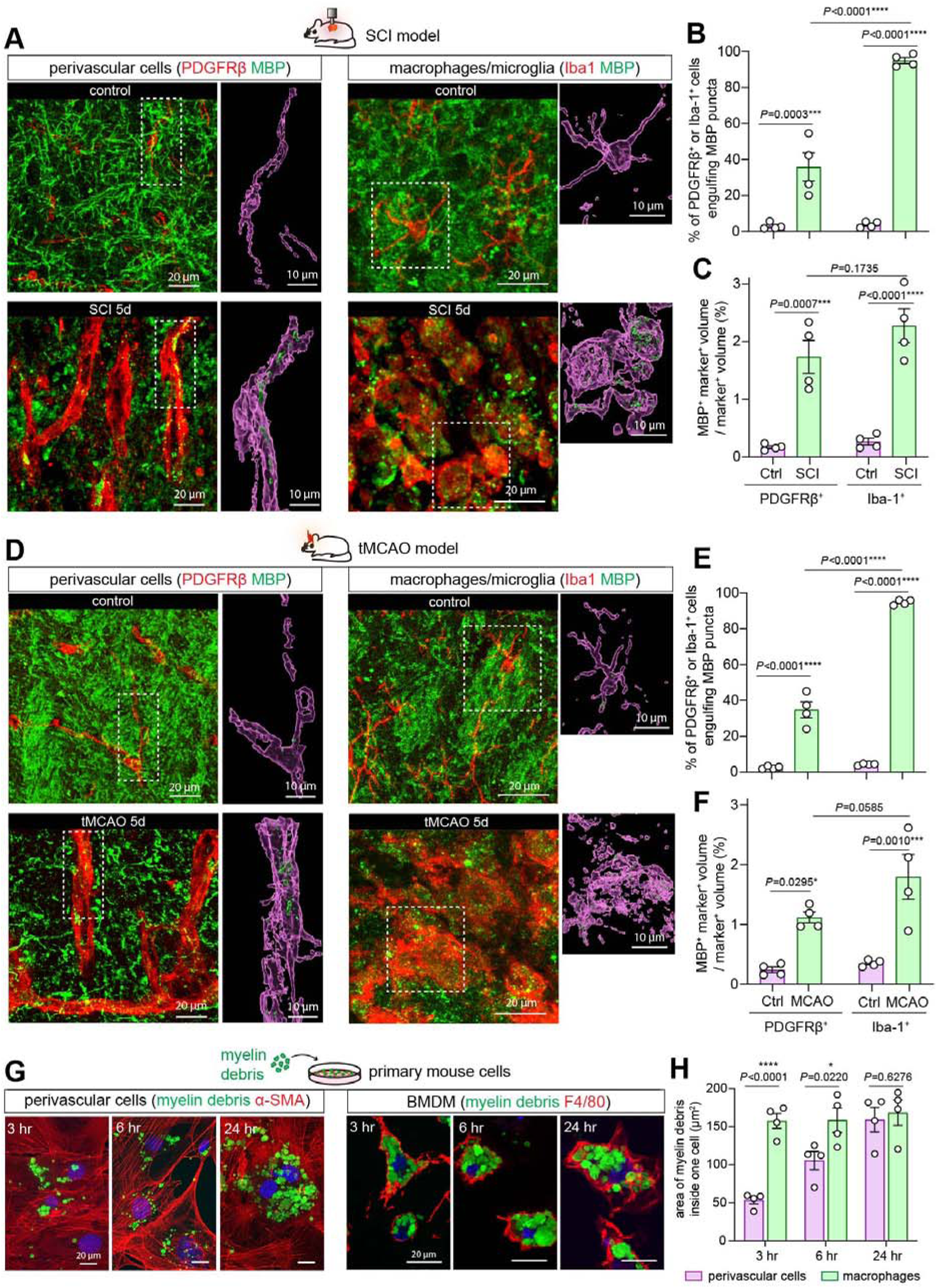
Comparison of myelin debris phagocytosis by perivascular cells and macrophages/microglia *in vivo* and *in vitro*. **(A)** Representative confocal images of spinal cord sections from control and SCI mice at 5 days after injury showing myelin debris engulfment by PDGFRβ^⁺^ perivascular cells and Iba-1^⁺^ macrophages/microglia. Higher-magnification images show 3D reconstructions of intracellular myelin debris. Scale bars: 20 μm; 10 μm (3D reconstructions). **(B, C)** Quantification of the phagocytic percentage (percentage of PDGFRβ^⁺^ or Iba-1^⁺^ cells containing MBP^⁺^ myelin debris) **(B)** and phagocytic volume (percentage of intracellular MBP^⁺^ volume relative to total cell volume) **(C)** in control and SCI mice. Data are presented as mean ± SEM (n = 4 mice). Statistical significance was determined by two-way ANOVA with Holm–Šídák’s multiple-comparisons test. **(D)** Representative confocal images of brain sections from control and transient middle cerebral artery occlusion (tMCAO) mice at 5 days after injury showing myelin debris engulfment by PDGFRβ^⁺^ perivascular cells and Iba-1^⁺^ macrophages/microglia. Higher-magnification images show 3D reconstructions of intracellular myelin debris. Scale bars: 20 μm; 10 μm (3D reconstructions). **(E, F)** Quantification of the phagocytic percentage (percentage of PDGFRβ^⁺^ or Iba-1^⁺^ cells containing MBP^⁺^ myelin debris) **(E)** and phagocytic volume (percentage of intracellular MBP^⁺^ volume relative to total cell volume) **(F)** in control and tMCAO mice. Data are presented as mean ± SEM (n = 4 mice). Statistical significance was determined by two-way ANOVA with Holm–Šídák’s multiple-comparisons test. **(G)** Representative immunocytochemical images comparing uptake of CFSE-labeled myelin debris by primary mouse brain perivascular cells (α-SMA^⁺^) and bone marrow-derived macrophages (BMDMs; F4/80^⁺^) at 3 hr, 6 hr and 24 hr after coculture. Scale bar: 20 μm. **(H)** Quantification of intracellular myelin debris area (CFSE-positive area normalized to cell number) in primary perivascular cells and BMDMs. Data are presented as mean ± SEM (n = 4 biological replicates). Statistical significance was determined by two-way ANOVA followed by Holm–Šídák’s multiple-comparisons test.

**Figure S11 – Related to Figure 2.**
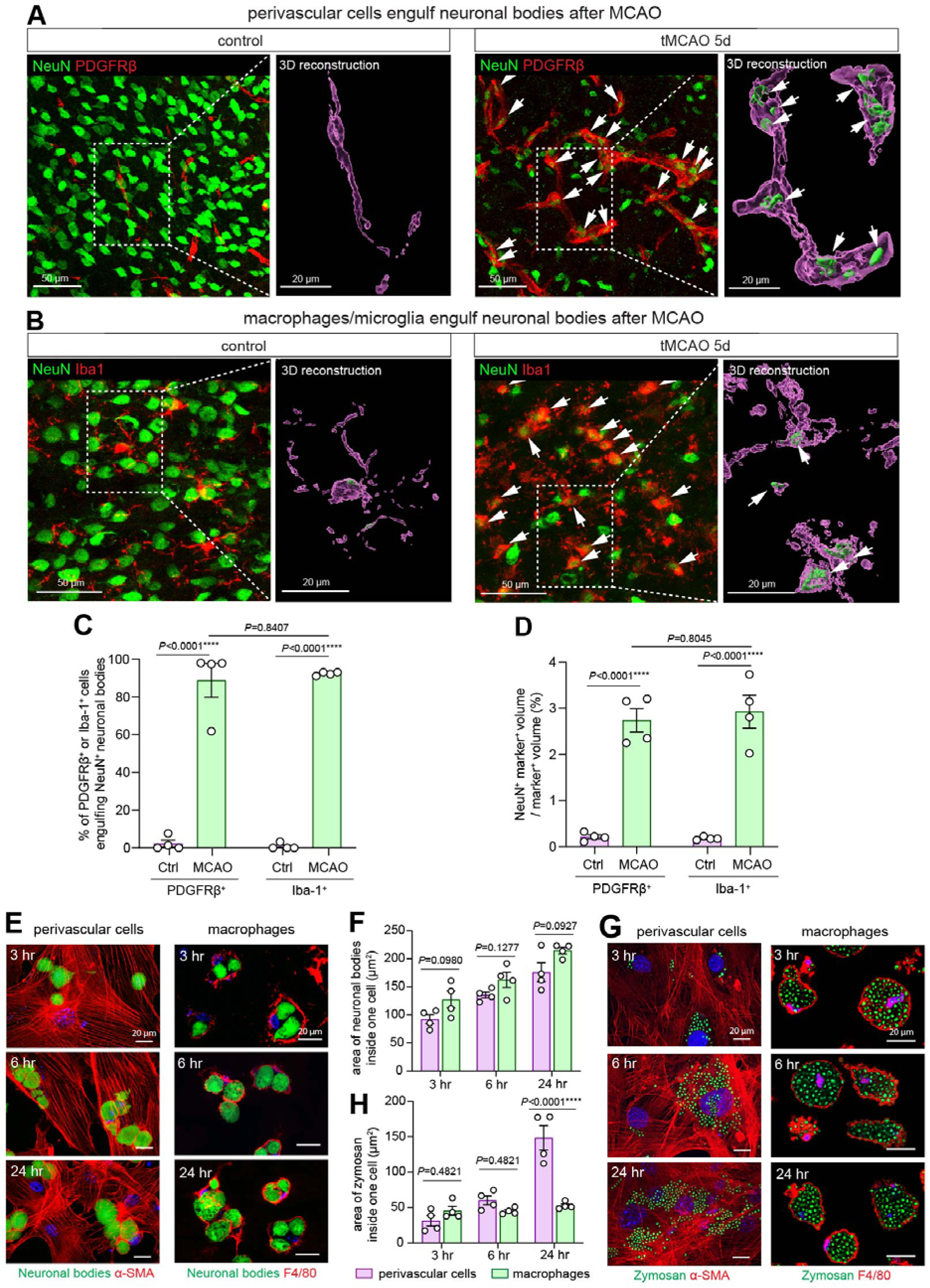
Perivascular cells exhibit broad phagocytic capacity comparable to macrophages *in vivo* and *in vitro*. **(A)** Representative confocal images of brain sections from mice at 5 days after transient middle cerebral artery occlusion (tMCAO) showing engulfment of NeuN^⁺^ neuronal bodies by PDGFRβ^⁺^ perivascular cells. Higher-magnification images show orthogonal views and 3D reconstructions. Arrows indicate engulfed neuronal bodies. Scale bars: 50 μm; 20 μm (3D reconstructions). **(B)** Representative confocal images of brain sections from mice at 5 days after tMCAO showing engulfment of NeuN^⁺^ neuronal bodies by Iba-1^⁺^ microglia/macrophages. Higher-magnification images include orthogonal views and 3D reconstructions. Arrows indicate engulfed neuronal bodies. Scale bars: 50 μm; 20 μm (3D reconstructions). **(C, D)** Quantification of the phagocytic percentage (percentage of PDGFRβ^⁺^ or Iba-1^⁺^ cells containing NeuN^⁺^ neuronal bodies) **(C)** and phagocytic volume (percentage of intracellular NeuN^⁺^ volume relative to total cell volume) **(D)** in control and tMCAO mice. Data are presented as mean ± SEM (n = 4 mice). Statistical significance was determined by two-way ANOVA followed by Holm–Šídák’s multiple-comparisons test. **(E)** Representative immunocytochemical images comparing uptake of CFSE-labeled neuronal bodies by primary mouse brain perivascular cells (α-SMA^⁺^) and bone marrow-derived macrophages (BMDMs; F4/80^⁺^) at 3, 6, and 24 hr after coculture. Scale bar: 20 μm. **(F)** Quantification of intracellular neuronal bodies area (CFSE-positive area normalized to total cell number) in primary perivascular cells and BMDMs. Data are presented as mean ± SEM (n = 4 biological replicates). Statistical significance was determined by two-way ANOVA followed by Holm–Šídák’s multiple-comparisons test. **(G)** Representative immunocytochemical images comparing uptake of CFSE-labeled zymosan particles by primary mouse brain perivascular cells (α-SMA^⁺^) and bone marrow-derived macrophages (BMDMs; F4/80^⁺^) at 3, 6, and 24 hr after coculture. Scale bar: 20 μm. **(H)** Quantification of intracellular zymosan area (CFSE-positive area normalized to total cell number) in primary perivascular cells and BMDMs. Data are presented as mean ± SEM (n = 4 biological replicates). Statistical significance was determined by two-way ANOVA followed by Holm–Šídák’s multiple-comparisons test.

**Figure S12 – Related to Figure 2.**
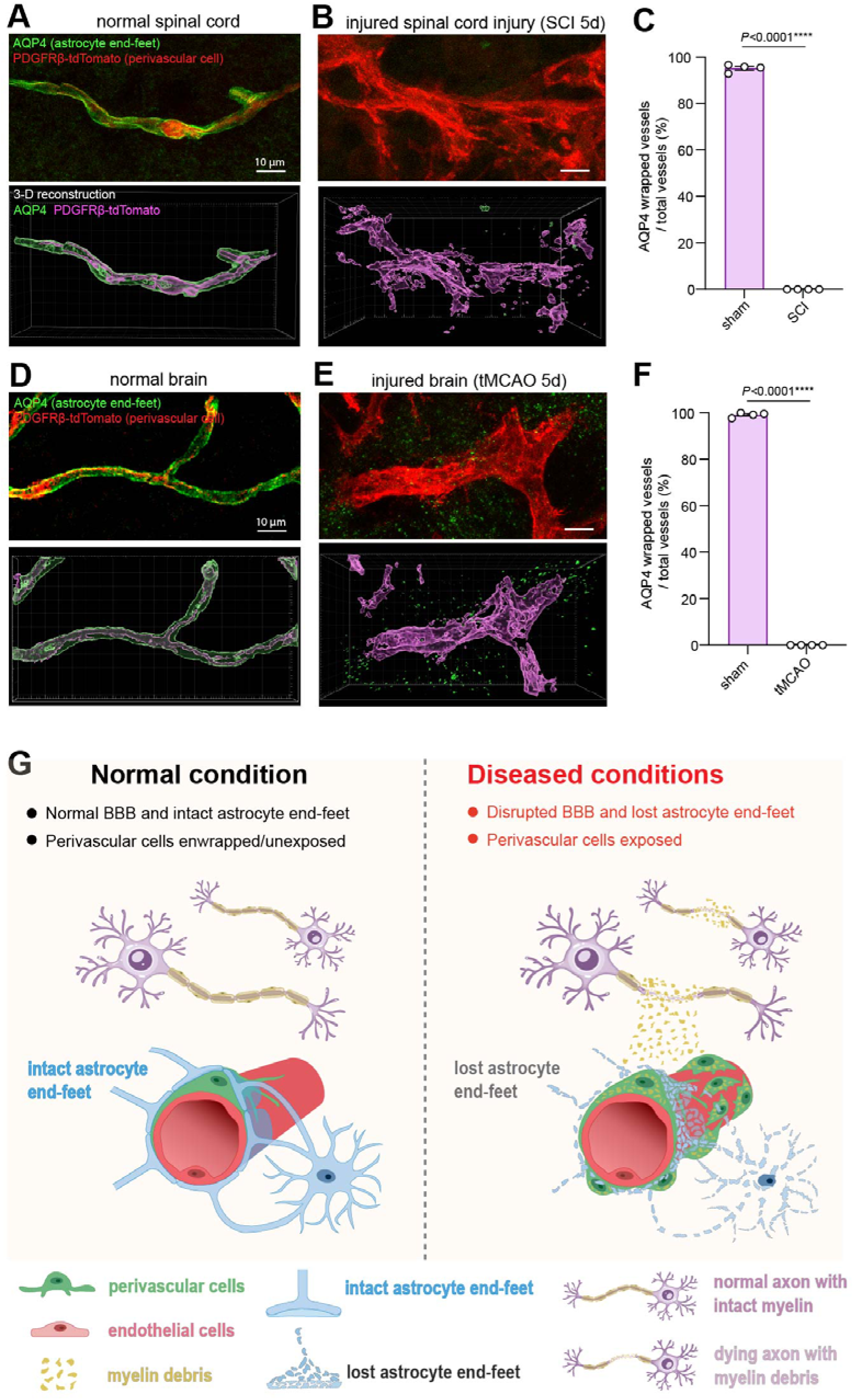
Loss of astrocytic end-feet coverage exposes perivascular cells to the injured CNS parenchyma. **(A, B)** Representative confocal images of spinal cord sections from sham **(A)** and SCI mice at 5 days after injury **(B)** showing astrocytic end-feet (AQP4, green) surrounding PDGFRβ-tdTomato^⁺^ perivascular cells (red). Lower panels show 3D reconstructions. Scale bar: 10 μm. **(C)** Quantification of the percentage of blood vessels covered by AQP4^⁺^ astrocytic end-feet in sham and SCI mice. Data are presented as mean ± SEM (n = 4 mice). Statistical significance was determined by an unpaired two-tailed *t* test with Welch’s correction. **(D, E)** Representative confocal images of brain sections from control **(D)** and transient middle cerebral artery occlusion (tMCAO) mice at 5 days after injury **(E)** showing astrocytic end-feet (AQP4, green) surrounding PDGFRβ-tdTomato^⁺^ perivascular cells (red). Lower panels show 3D reconstructions. Scale bar: 10 μm. **(F)** Quantification of the percentage of blood vessels covered by AQP4^⁺^ astrocytic end-feet in control and tMCAO mice. Data are presented as mean ± SEM (n = 4 mice). Statistical significance was determined by an unpaired two-tailed *t* test with Welch’s correction. **(G)** Proposed model illustrating how disruption of astrocytic end-feet following CNS injury exposes perivascular cells to parenchymal myelin debris. Under physiological conditions, perivascular cells are enclosed by astrocytic end-feet as part of the blood-brain/blood-spinal cord barrier and are largely isolated from parenchymal components. Following CNS injury, loss of astrocytic end-foot coverage disrupts the barrier, allowing perivascular cells to directly contact and engulf myelin debris.

**Figure S13 – Related to Figure 3.**
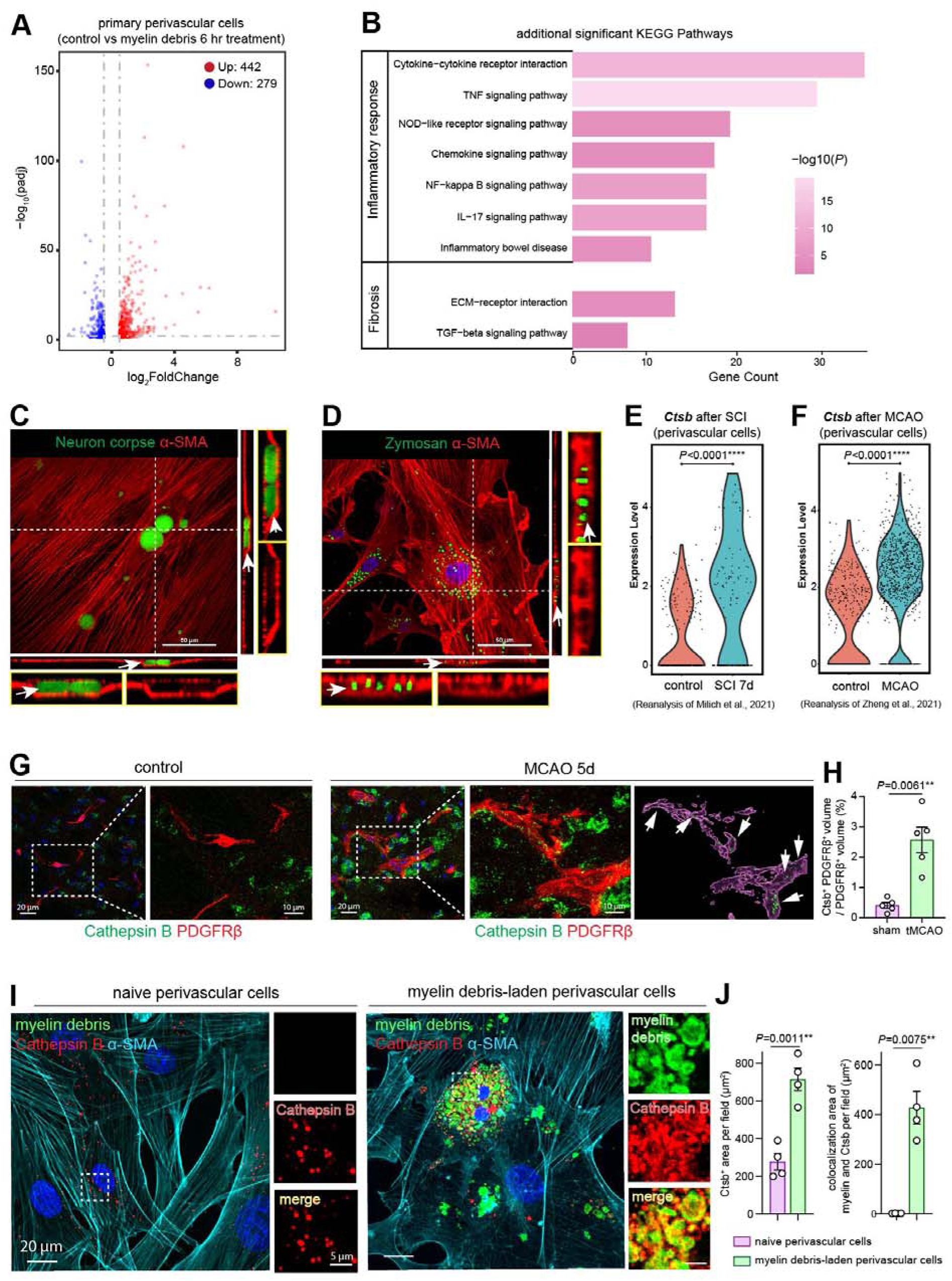
Molecular and cellular machinery underlying perivascular cell phagocytosis and lysosomal degradation. **(A)** Volcano plot showing differentially expressed genes (DEGs) in primary perivascular cells following 6 h exposure to myelin debris. Genes with |log_2_FoldChange| > 0.5 and FDR-adjusted *P* < 0.05 were considered differentially expressed. **(B)** KEGG pathway analysis of RNA sequencing data showing enrichment of pathways associated with inflammation and fibrosis in primary perivascular cells following myelin debris exposure. **(C, D)** Representative immunocytochemical images showing the formation of α-SMA^⁺^ actin-rich phagocytic cups (arrows) surrounding CFSE-labeled neuronal bodies **(C)** or zymosan particles **(D)** in primary perivascular cells after 6 hr of coculture. Scale bar: 50 μm. **(E, F)** Reanalysis of published single-cell RNA sequencing datasets showing Cathepsin B (Ctsb) expression in perivascular cells following SCI **(E)** and transient middle cerebral artery occlusion (tMCAO) **(F)**. **(G)** Representative confocal images of brain sections from control and tMCAO mice showing Cathepsin B (Ctsb) expression in PDGFRβ^⁺^ perivascular cells. Higher-magnification images show orthogonal views and 3D reconstructions illustrating increased Ctsb expression after tMCAO. Scale bars: 20 μm; 10 μm (higher magnification). **(H)** Quantification of the percentage of intracellular Ctsb volume occupied in the total volume of PDGFRβ^⁺^ perivascular cells. Data are presented as mean ± SEM (n = 5 mice). Statistical significance was determined by an unpaired two-tailed *t* test with Welch’s correction. **(I)** Representative immunocytochemical images showing Ctsb expression in primary perivascular cells under basal conditions or following myelin debris uptake. Scale bars: 20 μm; 5 μm (higher magnification). **(J)** Quantification of the intracellular Ctsb-positive area per imaging field and the colocalization area between Ctsb and myelin debris per imaging field. Data are presented as mean ± SEM (n = 4 biological replicates). Statistical significance was determined by an unpaired two-tailed *t* test with Welch’s correction.

**Figure S14 – Related to Figure 3.**
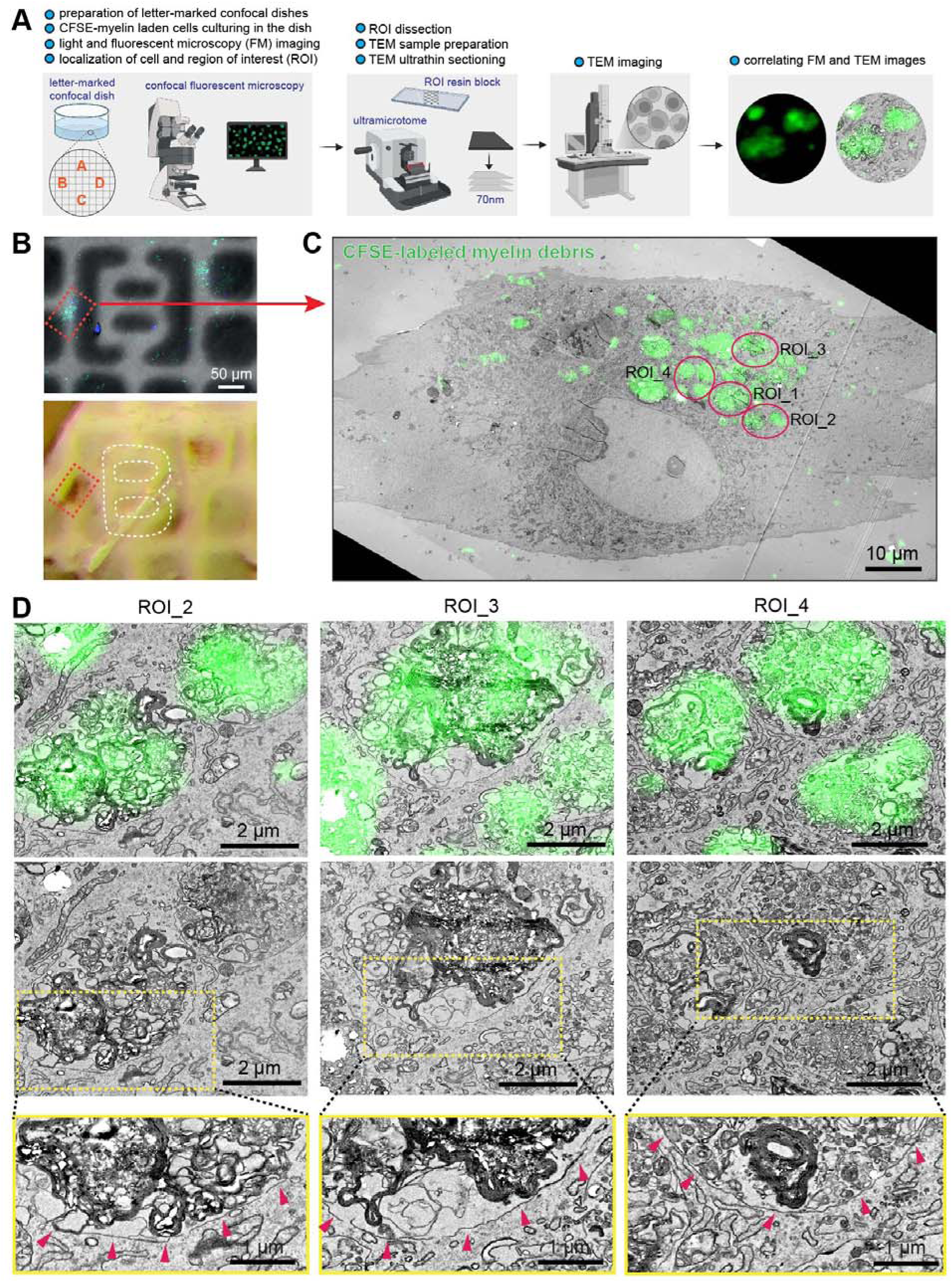
Correlative light and electron microscopy (CLEM) reveals the subcellular localization of myelin debris and the intracellular vesicular structures in perivascular cells. **(A)** Workflow illustrating the CLEM procedure used to examine the intracellular localization of CFSE-labeled myelin debris in primary perivascular cells following *in vitro* phagocytosis. **(B)** Brightfield and fluorescence images of a perivascular cell adjacent to a fiducial landmark used for CLEM registration (top), and the corresponding resin-embedded sample before ultrathin sectioning (bottom). **(C)** Low-magnification correlated fluorescence and transmission electron microscopy (TEM) image of a myelin debris-laden perivascular cell. Circles indicate regions of interest (ROI_1–4) selected for higher-magnification analysis. Scale bar: 10 μm. **(D)** High-magnification CLEM images showing CFSE-labeled myelin debris within single-membrane intracellular vesicles of perivascular cells. Arrows indicate single-membrane vesicles enclosing myelin debris. Scale bars: 2 μm; 1 μm (higher magnification).

**Figure S15 – Related to Figure 4.**
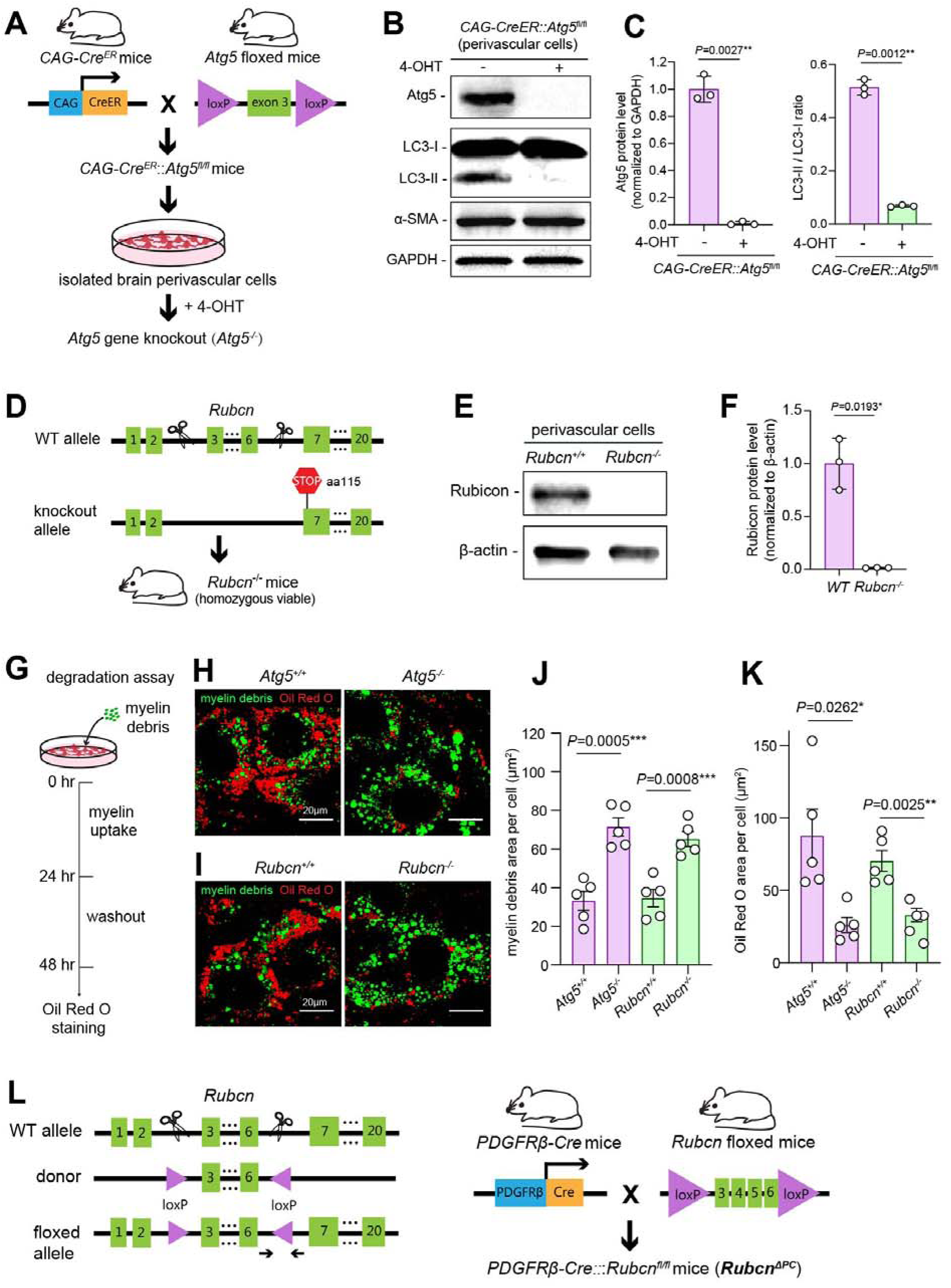
Generation and validation of *Atg5* and *Rubcn* knockout perivascular cells. **(A)** Diagram illustrating the generation of *CAG-Cre^ER^*::*Atg5^fl/fl^* mice and *in vitro* deletion of *Atg5* in primary perivascular cells. Primary perivascular cells were isolated from *CAG-Cre^ER^*::*Atg5^fl/fl^* mice and treated with 4-hydroxytamoxifen (4-OHT; 200 nM) for 48 hr to induce Cre-mediated excision of the floxed *Atg5* allele. Untreated cells served as controls. **(B)** Representative immunoblots validating deletion of *Atg5* in primary perivascular cells isolated from *CAG-Cre^ER^*::*Atg5^fl/fl^* mice. Loss of Atg5 protein and impaired LC3-I to LC3-II conversion following 4-OHT treatment confirmed functional disruption of *Atg5* gene. α-SMA served as a perivascular cell marker, and GAPDH as the loading control. **(C)** Quantification of Atg5 protein levels and LC3-II/LC3-I ratios shown in **(B)**. Data are presented as mean ± SEM (n = 3 biological replicates). Statistical significance was determined by an unpaired two-tailed *t* test with Welch’s correction. **(D)** Scheme illustrating generation of global *Rubcn* knockout (*Rubcn^−/−^*) mice and isolation of primary *Rubcn^−/−^* perivascular cells. CRISPR-Cas9-mediated deletion of a 6.1-kb genomic fragment encompassing exons 3–6 generated a premature stop codon at amino acid 115. **(E)** Representative immunoblots showing loss of Rubicon protein in primary *Rubcn^−/−^* perivascular cells. β-actin served as the loading control. **(F)** Quantification of Rubicon protein levels shown in **(E).** Data are presented as mean ± SEM (n = 3 biological replicates). Statistical significance was determined by an unpaired two-tailed *t* test with Welch’s correction. **(G)** Diagram illustrating the *in vitro* myelin degradation assay. Primary perivascular cells were loaded with CFSE-labeled myelin debris, followed by washout and subsequent analysis of intracellular myelin degradation and neutral lipid accumulation by Oil Red O staining. **(H, I)** Representative images of CFSE-labeled myelin debris and Oil Red O staining in myelin-laden *Atg5^+/+^* and *Atg5^−/−^* perivascular cells **(H),** and *Rubcn^+/+^* and *Rubcn^−/−^*perivascular cells **(I).** Compared with control cells, *Atg5*- or *Rubcn*-deficient perivascular cells retained more intracellular myelin debris and accumulated fewer neutral lipids, indicating impaired myelin degradation. Scale bar: 20 μm. **(J)** Quantification of the intracellular area occupied by CFSE-labeled myelin debris in control, *Atg5^−/−^*, and *Rubcn^−/−^* perivascular cells. Data are presented as mean ± SEM (n = 5 biological replicates). Statistical significance was determined by an unpaired two-tailed *t* test. **(K)** Quantification of Oil Red O^+^ area in myelin-laden control, *Atg5^−/−^*, and *Rubcn^−/−^* perivascular cells. Data are presented as mean ± SEM (n = 5 biological replicates). Statistical significance was determined by an unpaired two-tailed *t* test with or without Welch’s correction, as appropriate. **(L)** Diagram illustrating the generation of *PDGFR*β*-Cre*::*Rubcn ^fl/fl^* mice (*Rubcn*^Δ*PC*^) for perivascular cell-specific deletion of *Rubcn*. Two loxP sites were inserted by CRISPR/Cas9 to flank exons 3–6 of the endogenous *Rubcn* locus. Arrows indicate the primer pair used for PCR genotyping.

**Figure S16 – related to Figure 5.**
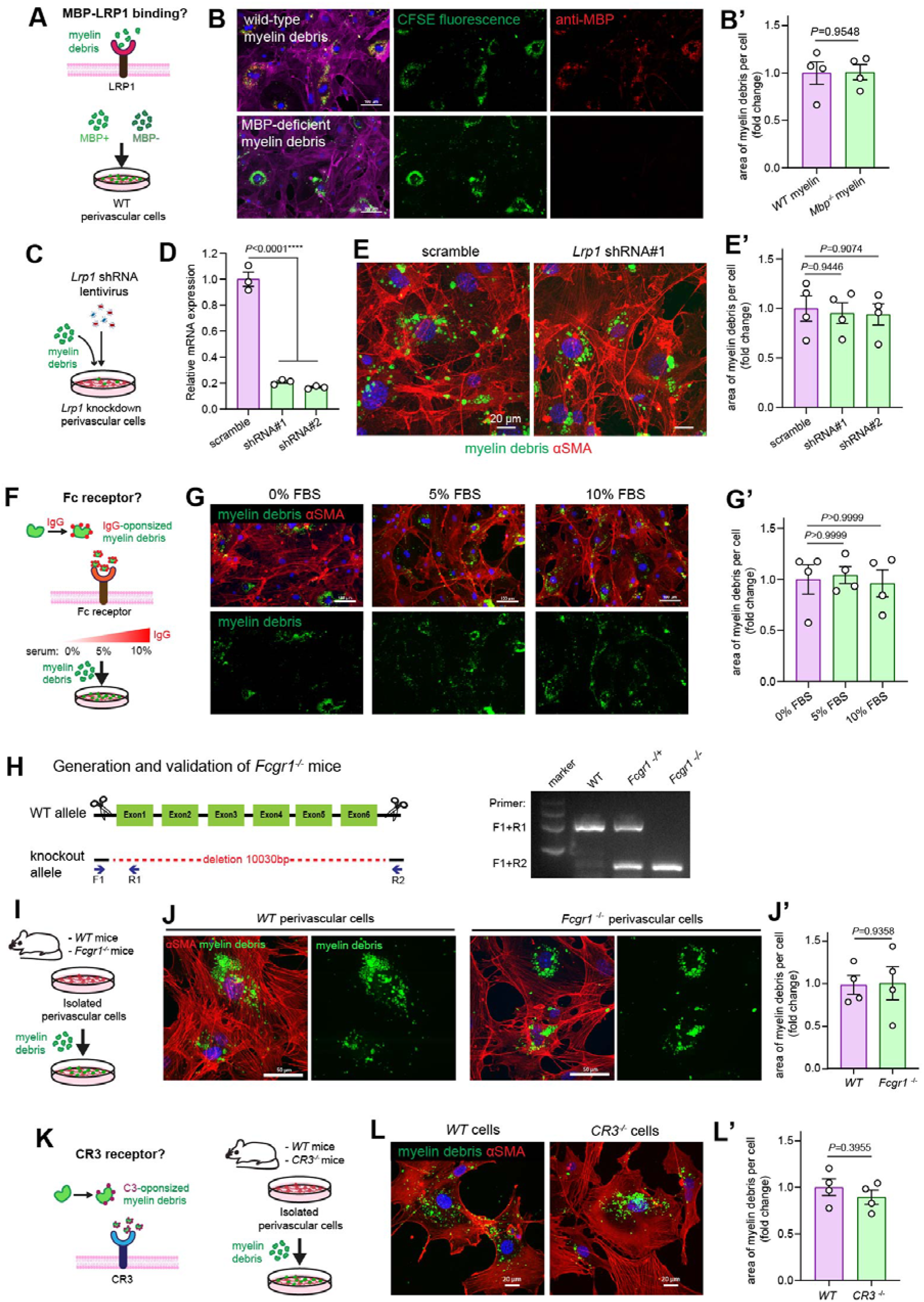
Canonical LRP1-, Fc receptor-, and CR3-mediated phagocytic pathways are dispensable for myelin debris uptake by perivascular cells. **(A)** Experimental strategy to evaluate the role of the LRP1 receptor in perivascular cell phagocytosis of myelin debris. CFSE-labeled myelin debris was prepared from wild-type mice or *Mbp*^−/−^ mice lacking MBP and incubated with primary wild-type perivascular cells in vitro. **(B, B**′) Representative confocal images and quantification showing that MBP deficiency in myelin debris does not affect its uptake by perivascular cells. MBP immunostaining validates the absence of MBP in myelin debris derived from *Mbp*^−/−^ mice. CFSE labels myelin debris and α-SMA labels perivascular cells. Scale bar: 100 μm. Data are presented as mean ± SEM (*n* = 4 biological replicates). Statistical significance was determined by an unpaired two-tailed *t* test. **(C)** Experimental design for lentiviral knockdown of *Lrp1* in primary perivascular cells followed by analysis of CFSE-labeled myelin debris uptake. **(D)** RT-qPCR validation of *Lrp1* knockdown using two independent shRNAs. Scramble shRNA served as the control. Data are presented as mean ± SEM (*n* = 3 biological replicates). Statistical significance was determined by one-way ANOVA with Dunnett’s multiple-comparisons test. **(E, E**′) Representative confocal images and quantification showing that *Lrp1* knockdown does not alter myelin debris phagocytosis by primary perivascular cells. CFSE labels myelin debris and α-SMA labels perivascular cells. Scale bar: 20 μm. Data are presented as mean ± SEM (*n* = 4 biological replicates). Statistical significance was determined by one-way ANOVA with Dunnett’s multiple-comparisons test. **(F)** Experimental strategy to evaluate the role of Fc receptor-mediated IgG opsonization in perivascular cell phagocytosis. Primary wild-type perivascular cells were cultured in medium containing 0%, 5%, or 10% fetal bovine serum together with CFSE-labeled myelin debris. **(G, G**′) Representative confocal images and quantification showing comparable uptake of CFSE-labeled myelin debris by perivascular cells cultured under different serum concentrations. CFSE labels myelin debris and α-SMA labels perivascular cells. Scale bar: 100 μm. Data are presented as mean ± SEM (*n* = 4 biological replicates). Statistical significance was determined by the Kruskal–Wallis test with Dunn’s multiple-comparisons test. **(H)** Generation and validation of *Fcgr1^−/−^* mice. Left, CRISPR/Cas9 strategy used to generate a null *Fcgr1* allele by deleting the entire coding region. Right, PCR genotyping of wild-type, heterozygous, and homozygous knockout mice. **(I)** Experimental design for assessing myelin debris phagocytosis by primary perivascular cells isolated from wild-type and *Fcgr1^−/−^* mice. **(J, J**′) Representative confocal images and quantification showing that *Fcgr1* deficiency does not impair perivascular cell phagocytosis of CFSE-labeled myelin debris. Data are presented as mean ± SEM (*n* = 4 biological replicates). Statistical significance was determined by an unpaired two-tailed *t* test. **(K)** Experimental strategy to evaluate the role of CR3-mediated complement-dependent phagocytosis. Primary perivascular cells isolated from wild-type and *CR3*^−/−^ mice were incubated with CFSE-labeled myelin debris and analyzed for phagocytic uptake. **(L, L**′) Representative confocal images and quantification showing comparable uptake of CFSE-labeled myelin debris by wild-type and *CR3*^−/−^ perivascular cells. Data are presented as mean ± SEM (*n* = 4 biological replicates). Statistical significance was determined by an unpaired two-tailed *t* test.

**Figure S17 – related to Figure 5.**
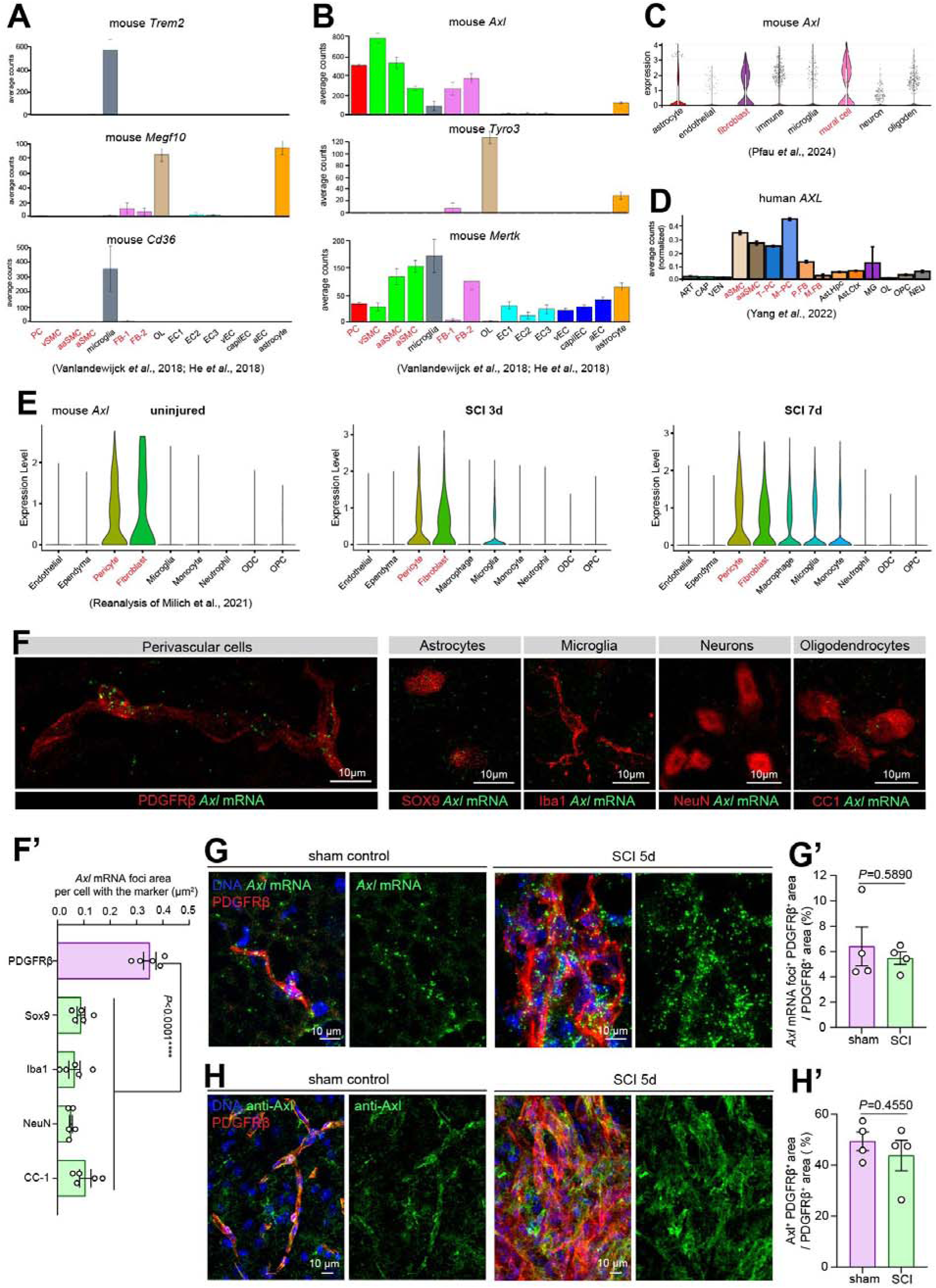
Identification of Axl as the phagocytic receptor in perivascular cells. **(A, B)** Reanalysis of mouse vascular single-cell RNA-sequencing datasets from Vanlandewijck *et al.* (2018) and He *et al.* (2018). Expression of phagocytic receptors **(A)** *Trem2, Megf10*, and *Cd36*, and TAM family receptors **(B)** *Axl, Tyro3*, and *Mertk* across various cell populations. PDGFRβ-expressing perivascular cell populations are highlighted in red and include pericytes (PC), venous smooth muscle cells (vSMC), arterial smooth muscle cells (aSMC), arteriolar smooth muscle cells (aaSMC), and fibroblast-like cells (FB-1 and FB-2). Note the selective enrichment of Axl in perivascular cell populations. **(C)** Reanalysis of an independent mouse vascular single-cell RNA-sequencing dataset from Pfau *et al.* (2024) showing Axl expression in PDGFRβ-expressing mural cells and fibroblasts. **(D)** Reanalysis of a human vascular single-cell RNA-sequencing dataset from Yang *et al.* (2022). Axl expression is enriched in PDGFRβ-expressing perivascular cell populations, which are highlighted in red and include including arterial smooth muscle cells (aSMC), arteriolar smooth muscle cells (aaSMC), transport-associated pericytes (T-PC), ECM-regulating pericytes (M-PC), perivascular fibroblasts (P.FB), and meningeal fibroblasts (M.FB). **(E)** Reanalysis of published single-cell RNA-sequencing data from spinal cords before and after SCI (Milich *et al.*, 2021). Axl expression is shown across spinal cord cell populations, with PDGFRβ-expressing pericytes and fibroblasts highlighted in red. **(F, F**′) Representative confocal images of Axl mRNA detected by HCR-FISH together with immunostaining for the indicated CNS cell-type markers in normal spinal cords **(F).** Scale bar, 10 μm. **(F**′) Quantification of Axl mRNA expression, measured as the area of Axl mRNA puncta normalized to the number of the corresponding cell marker. Data are presented as mean ± SEM (n = 5 mice). Statistical significance was determined by one-way ANOVA with Holm–Šídák’s multiple-comparisons test. **(G, G**′) Representative confocal images of Axl mRNA (HCR-FISH) in PDGFRβ^+^perivascular cells from sham-operated and 5-day SCI spinal cords **(G).** Scale bar, 10 μm. **(G**′) Quantification of Axl mRNA expression, measured as the area of Axl mRNA puncta normalized to the PDGFRβ^+^ area. Data are presented as mean ± SEM (n = 5 mice). Statistical significance was determined by an unpaired two-tailed *t* test. **(H, H**′) Representative confocal images of Axl protein immunostaining in PDGFRβ^+^ perivascular cells from sham-operated and 5-day SCI spinal cords **(H).** Scale bar, 10 μm. **(H**′) Quantification of Axl protein expression, measured as the Axl^+^ area normalized to the total PDGFRβ^+^ area. Data are presented as mean ± SEM (n = 4 mice). Statistical significance was determined by an unpaired two-tailed *t* test.

**Figure S18 – related to Figure 6.**
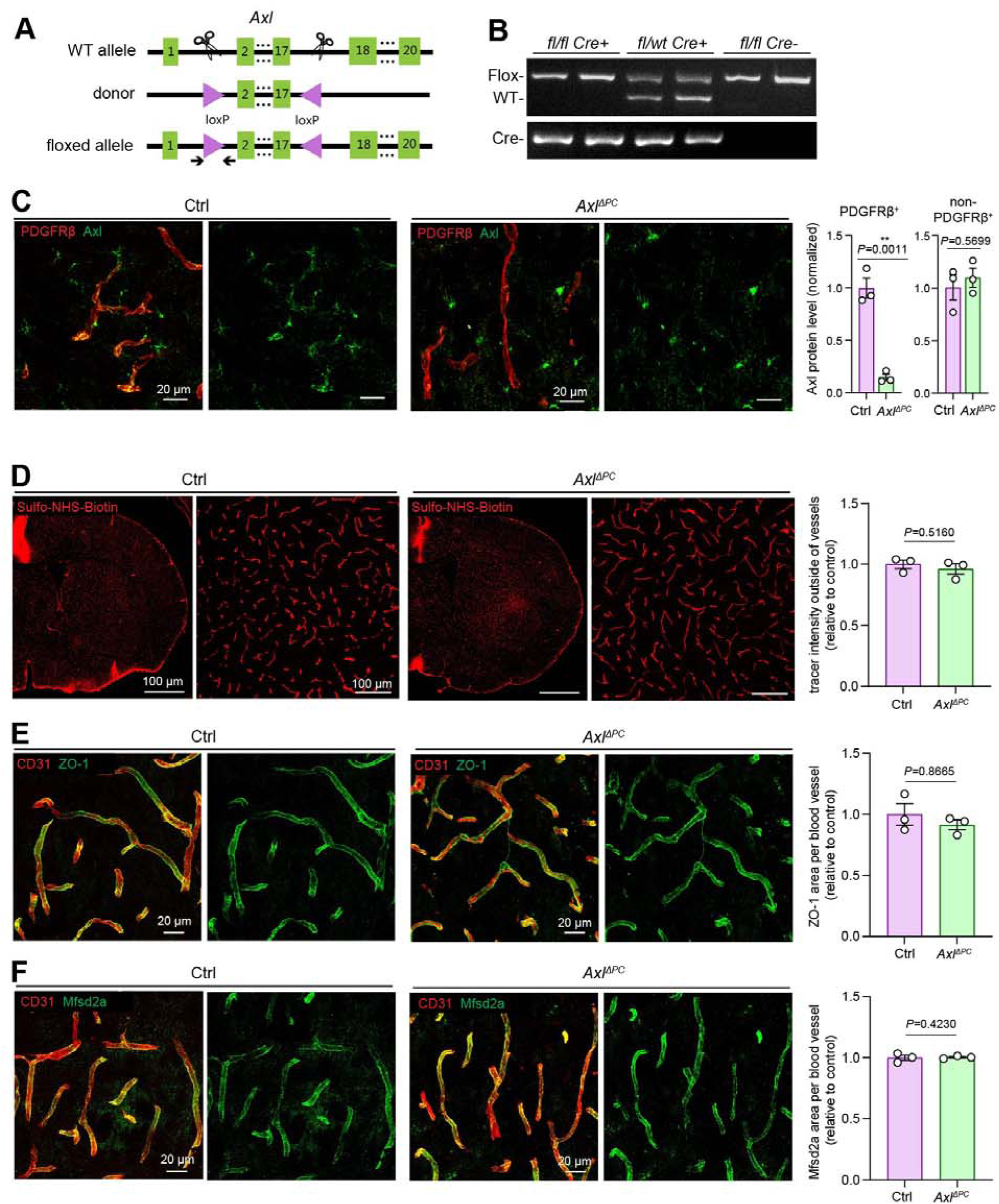
Perivascular cell-specific deletion of *Axl* does not affect blood–brain barrier integrity. **(A)** Schematic illustrating the generation of the *Axl* floxed allele for Cre-dependent deletion of *Axl*. The floxed allele was generated by CRISPR–Cas9-mediated insertion of loxP sites flanking exons 2–17 of the endogenous *Axl* locus. Arrows indicate the primer pair used for PCR genotyping. **(B)** Representative PCR genotyping of *Axl*^Δ*PC*^ mice. **(C)** Representative confocal images showing Axl immunostaining in brain sections from control and *Axl*^Δ*PC*^ mice, validating selective loss of Axl protein in PDGFRβ-positive perivascular cells while preserving Axl expression in non-perivascular cells. Scale bar, 20 μm. Right, quantification of Axl immunoreactivity in PDGFRβ-positive and PDGFRβ-negative cells. Data are presented as mean ± SEM (n = 3 mice). Statistical significance was determined by an unpaired two-tailed *t* test. **(D)** Representative confocal images showing blood–brain barrier permeability assessed by extravasation of the low-molecular-weight tracer Sulfo-NHS-biotin (443 Da) in control and *Axl*^Δ*PC*^ mouse brains. Scale bar, 100 μm. Right, quantification of extravasated Sulfo-NHS-biotin-positive area, normalized to the control group. Data are presented as mean ± SEM (n = 3 mice). Statistical significance was determined by an unpaired two-tailed *t* test. **(E)** Representative confocal images of ZO-1 immunostaining showing intact endothelial tight junctions in brain blood vessels of control and *Axl*^Δ*PC*^ mice. Scale bar, 20 μm. Right, quantification of ZO-1-positive area per blood vessel, normalized to control group. Data are presented as mean ± SEM (n = 3 mice). Statistical significance was determined by an unpaired two-tailed *t* test. **(F)** Representative confocal images of Mfsd2a immunostaining in brain blood vessels from control and *Axl*^Δ*PC*^ mice. Scale bar, 20 μm. Right, quantification of Mfsd2a-positive area per blood vessel, normalized to control group. Data are presented as mean ± SEM (n = 3 mice). Statistical significance was determined by an unpaired two-tailed *t* test.

**Figure S19 – related to Figure 6.**
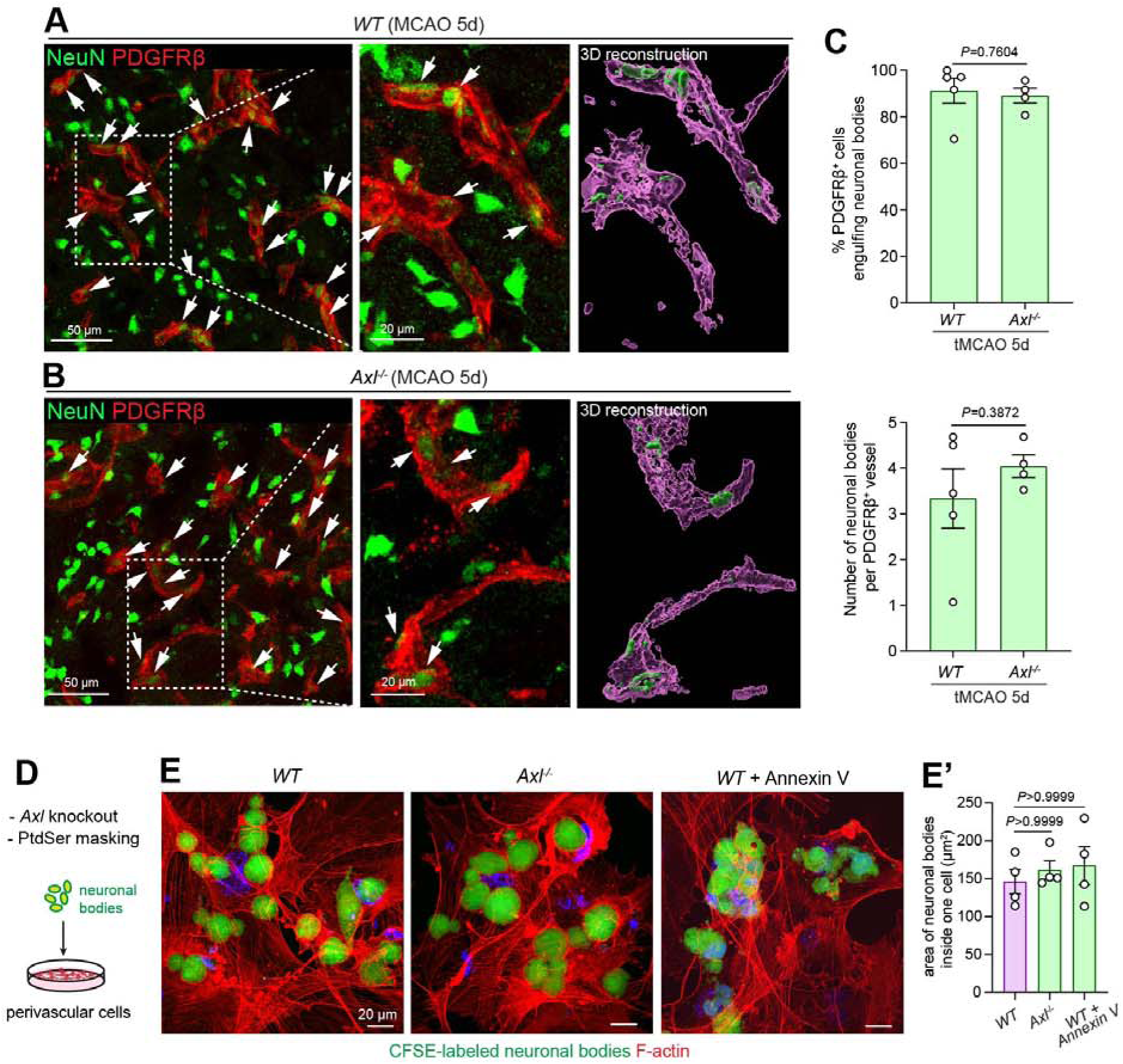
Perivascular cell engulfment of neuronal bodies is independent of Axl receptor. **(A, B)** Representative confocal images of brain sections from wild-type and *Axl^−/−^* mice showing PDGFRβ-positive perivascular cells engulfing NeuN-positive neuronal bodies 5 days after tMCAO. Enlarged views show 3D reconstructions of intracellular neuronal bodies. Arrows indicate engulfed neuronal bodies. Scale bar, 50 μm; 20 μm (3D views). **(C)** Quantification of the percentage of PDGFRβ-positive perivascular cells containing neuronal bodies (top) and the number of intracellular neuronal bodies per PDGFRβ-positive vessel (bottom). Data are presented as mean ± SEM (wild-type, *n* = 5 mice; *Axl^−/−^*, *n* = 4 mice). Statistical significance was determined by an unpaired two-tailed *t* test. **(D)** Schematic illustrating the experimental design for assessing *in vitro* phagocytosis of CFSE-labeled neuronal bodies by primary perivascular cells following Axl deletion or phosphatidylserine (PtdSer) masking with recombinant Annexin V (4 μg). **(E, E**′) Representative confocal images **(E)** and quantification **(E**′) of CFSE-labeled neuronal bodies uptake by primary mouse brain perivascular cells following Axl deletion or Annexin V treatment. F-actin was labeled with phalloidin. Quantification shows the intracellular area of CFSE-positive neuronal bodies per cell. Scale bar, 20 μm. Data are presented as mean ± SEM (*n* = 4 biological replicates). Statistical significance was determined by the Kruskal–Wallis test with Dunn’s multiple-comparisons test.

**Figure S20 – related to Figure 6.**
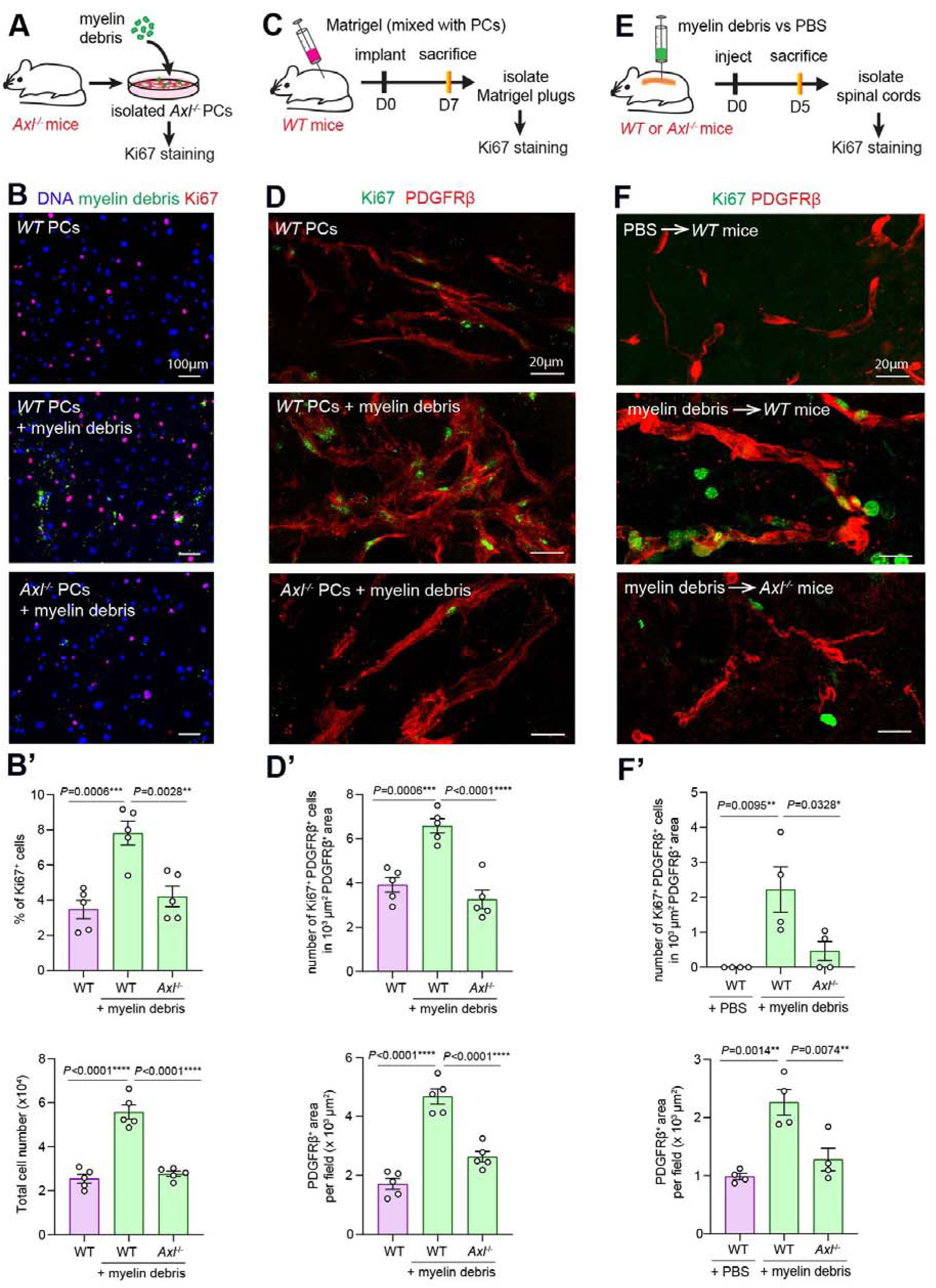
Axl-dependent engulfment of myelin debris promotes perivascular cell proliferation. **(A)** Schematic illustrating the experimental design for isolating primary WT and *Axl^−/−^*perivascular cells and assessing their proliferative response to myelin debris by Ki67 immunostaining. **(B, B**′) Representative confocal images **(B)** and quantification **(B**′) of WT and *Axl^−/−^*primary perivascular cells cultured in the presence or absence of myelin debris for 72 hr. Ki67 immunostaining was used to identify proliferating cells, and DAPI was used to label nuclei. Scale bar, 100 μm. Quantification shows the percentage of Ki67-positive cells (top) and total cell number (bottom). Data are presented as mean ± SEM (*n* = 5 biological replicates). Statistical significance was determined by one-way ANOVA followed by Tukey’s multiple-comparisons test. **(C)** Schematic illustrating the Matrigel plug assay. WT and *Axl^−/−^* perivascular cells with or without prior exposure to myelin debris were mixed with Matrigel, implanted subcutaneously into WT mice, and harvested 7 days later for analysis of perivascular cell proliferation by Ki67 immunostaining. **(D, D**′) Representative confocal images **(D)** and quantification **(D**′) of Matrigel plugs containing WT or *Axl^−/−^* perivascular cells with or without myelin debris pretreatment. Ki67 immunostaining was used to assess cell proliferation and PDGFRβ immunostaining to identify perivascular cells. Scale bar, 20 μm. Quantification shows Ki67-positive cells normalized to PDGFRβ-positive area (top) and total PDGFRβ-positive area (bottom). Data are presented as mean ± SEM (*n* = 5 mice). Statistical significance was determined by one-way ANOVA followed by Tukey’s multiple-comparisons test. **(E)** Schematic illustrating the spinal cord myelin debris microinjection assay. CFSE-labeled myelin debris or PBS (control) was injected into the spinal cords of WT or *Axl^−/−^*mice, and spinal cords were collected 5 days later to assess the proliferative response of perivascular cells by Ki67 immunostaining. **(F, F**′) Representative confocal images **(F)** and quantification **(F**′) of perivascular cell proliferation following spinal cord injection of myelin debris or PBS. Ki67 immunostaining was used to identify proliferating cells and PDGFRβ immunostaining to identify perivascular cells. Scale bar, 20 μm. Quantification shows Ki67-positive cells normalized to PDGFRβ-positive area (top) and total PDGFRβ-positive area (bottom). Data are presented as mean ± SEM (*n* = 4 mice). Statistical significance was determined by one-way ANOVA followed by Tukey’s multiple-comparisons test.

**Figure S21 – related to Figure 6.**
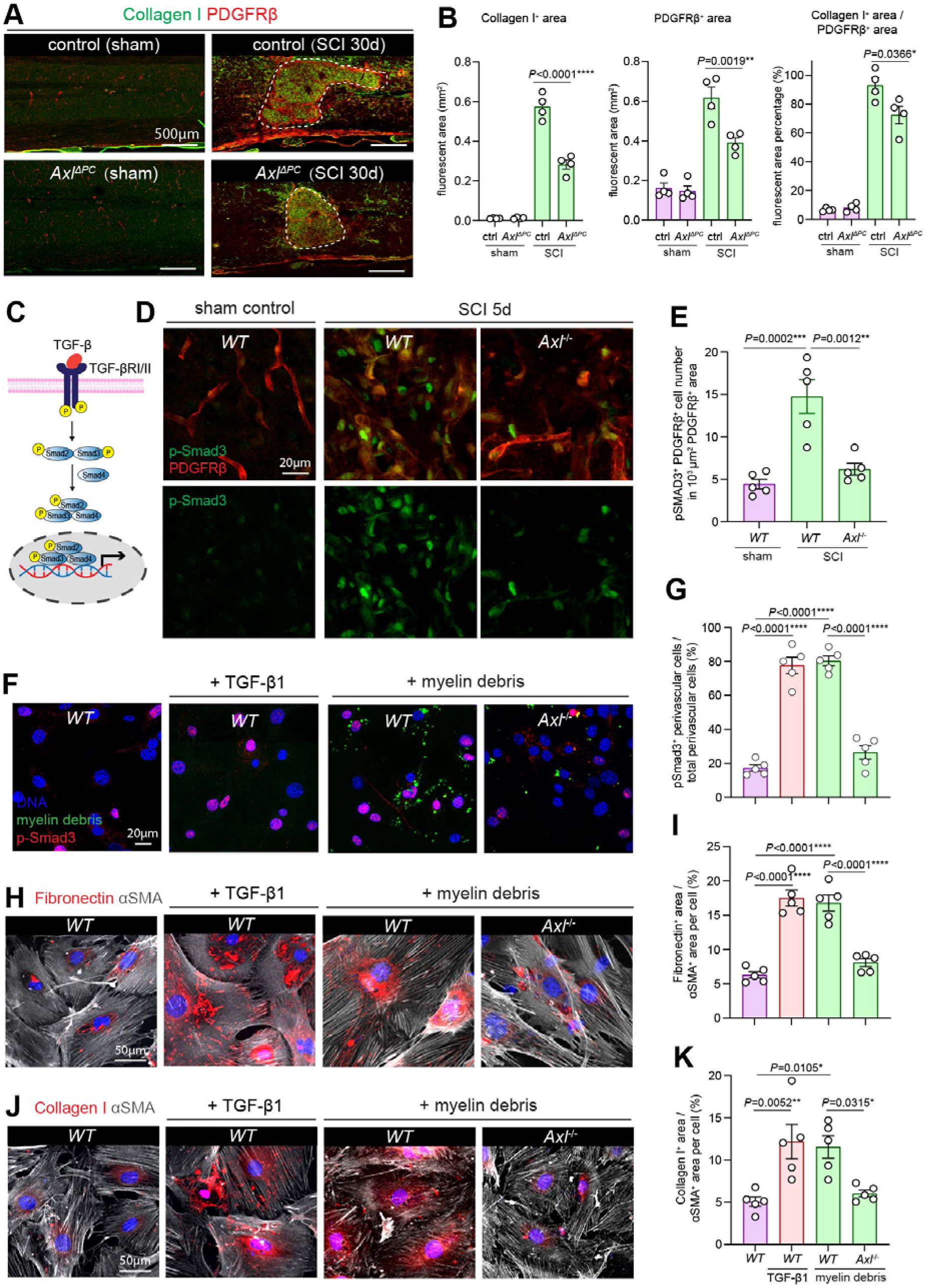
Axl-dependent engulfment of myelin debris activates TGF-β signaling and promotes production of pro-fibrotic extracellular matrix proteins. **(A)** Representative confocal images of collagen I immunostaining in spinal cords from control and *Axl*^Δ*PC*^ mice 30 days after SCI. Dashed outlines indicate the lesion area. Scale bar, 500 μm. **(B)** Quantification of collagen I-positive area (left), PDGFRβ-positive area (middle), and collagen I-positive area normalized to PDGFRβ-positive area (right) in spinal cords from the indicated mice. Data are presented as mean ± SEM (*n* = 4 mice). Statistical significance was determined by two-way ANOVA followed by Holm–Šídák’s multiple-comparisons test. **(C)** Schematic illustrating canonical TGF-β signaling. Binding of TGF-β to its receptors induces Smad3 phosphorylation, followed by nuclear translocation of phosphorylated Smad3 (p-Smad3) to regulate transcription of pro-fibrotic genes. **(D, E)** Representative confocal images **(D)** and quantification **(E)** of nuclear p-Smad3 in PDGFRβ-positive perivascular cells from WT and *Axl^−/−^* mice 5 days after SCI. p-Smad3 and PDGFRβ were detected by immunostaining. Scale bar, 20 μm. Quantification shows the percentage of perivascular cells with nuclear p-Smad3. Data are presented as mean ± SEM (*n* = 5 mice). Statistical significance was determined by one-way ANOVA followed by Tukey’s multiple-comparisons test. **(F, G)** Representative confocal images **(F)** and quantification **(G)** of nuclear p-Smad3 in primary WT and *Axl^−/−^* perivascular cells cultured in the presence or absence of myelin debris. Recombinant TGF-β1 served as a positive control. p-Smad3 was detected by immunostaining, CFSE labeled myelin debris, and DAPI labeled nuclei. Scale bar, 20 μm. Quantification shows the percentage of cells with nuclear p-Smad3. Data are presented as mean ± SEM (*n* = 5 biological replicates). Statistical significance was determined by one-way ANOVA followed by Tukey’s multiple-comparisons test. **(H–K)** Representative confocal images and quantification of fibronectin **(H, I)** and collagen I **(J, K)** production by primary WT and *Axl^−/−^* perivascular cells cultured in the presence or absence of myelin debris. Recombinant TGF-β1 served as a positive control. Fibronectin, collagen I, and α-SMA were detected by immunostaining. Scale bar, 50 μm. Quantification shows fibronectin-positive area **(I)** and collagen I-positive area **(K)** normalized to α-SMA-positive area. Data are presented as mean ± SEM (*n* = 5 biological replicates). Statistical significance was determined by one-way ANOVA followed by Tukey’s multiple-comparisons test.

**Figure S22 – related to Figure 6.**
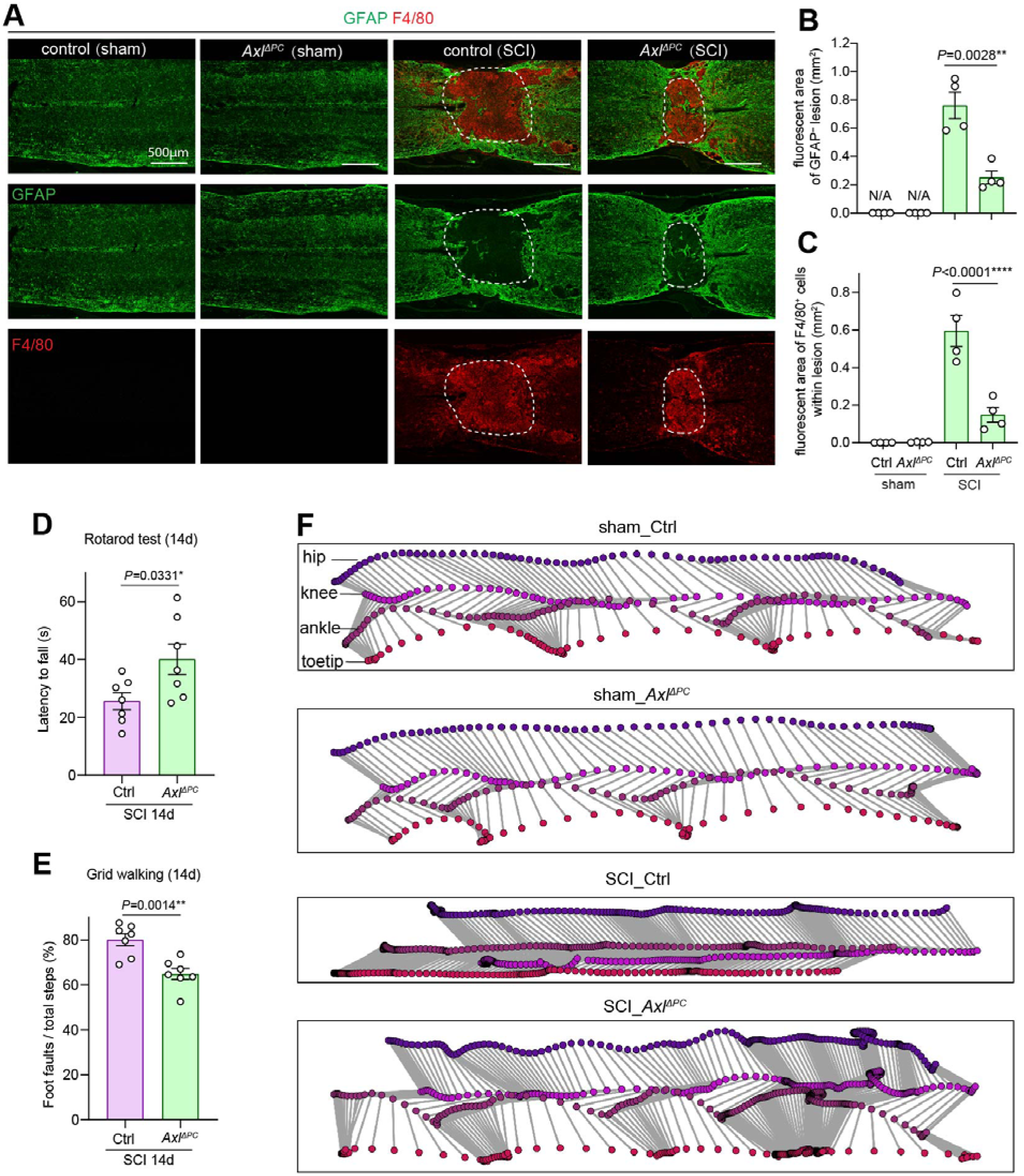
Perivascular cell-specific deletion of Axl attenuates neuroinflammation and improves functional recovery after spinal cord injury. **(A)** Representative confocal images of GFAP (green) and F4/80 (red) immunostaining in spinal cords from control and *Axl*^Δ*PC*^ mice 42 days after sham or SCI. GFAP labels the astroglial scar and F4/80 labels infiltrated macrophages. Dashed boxes indicate the regions of interest used for quantification. Scale bar, 500 μm. **(B, C)** Quantification of the GFAP-negative lesion area **(B)** and F4/80-positive area **(C)** in spinal cords from control and *Axl*^Δ*PC*^ mice 42 days after sham or SCI. Data are presented as mean ± SEM (*n* = 4 mice). Statistical significance was determined by an unpaired two-tailed *t* test **(B)** and two-way ANOVA followed by Holm–Šídák’s multiple-comparisons test **(C)**. **(D)** Quantification of rotarod performance, measured as latency to fall, in control and *Axl*^Δ*PC*^ mice 14 days after SCI. Data are presented as mean ± SEM (*n* = 7 mice). Statistical significance was determined by an unpaired two-tailed *t* test. **(E)** Quantification of grid-walking performance, measured as the percentage of faulty foot placements, in control and *Axl*^Δ*PC*^ mice 14 days after SCI. Data are presented as mean ± SEM (*n* = 7 mice). Statistical significance was determined by an unpaired two-tailed *t* test. **(F)** Representative images showing markerless tracking and hindlimb kinematic analysis in control and *Axl*^Δ*PC*^ mice 21 days after sham or SCI. Colored markers indicate the hip, knee, ankle, and toe joints used for motion tracking.

**Figure S23 – related to Figure 7.**
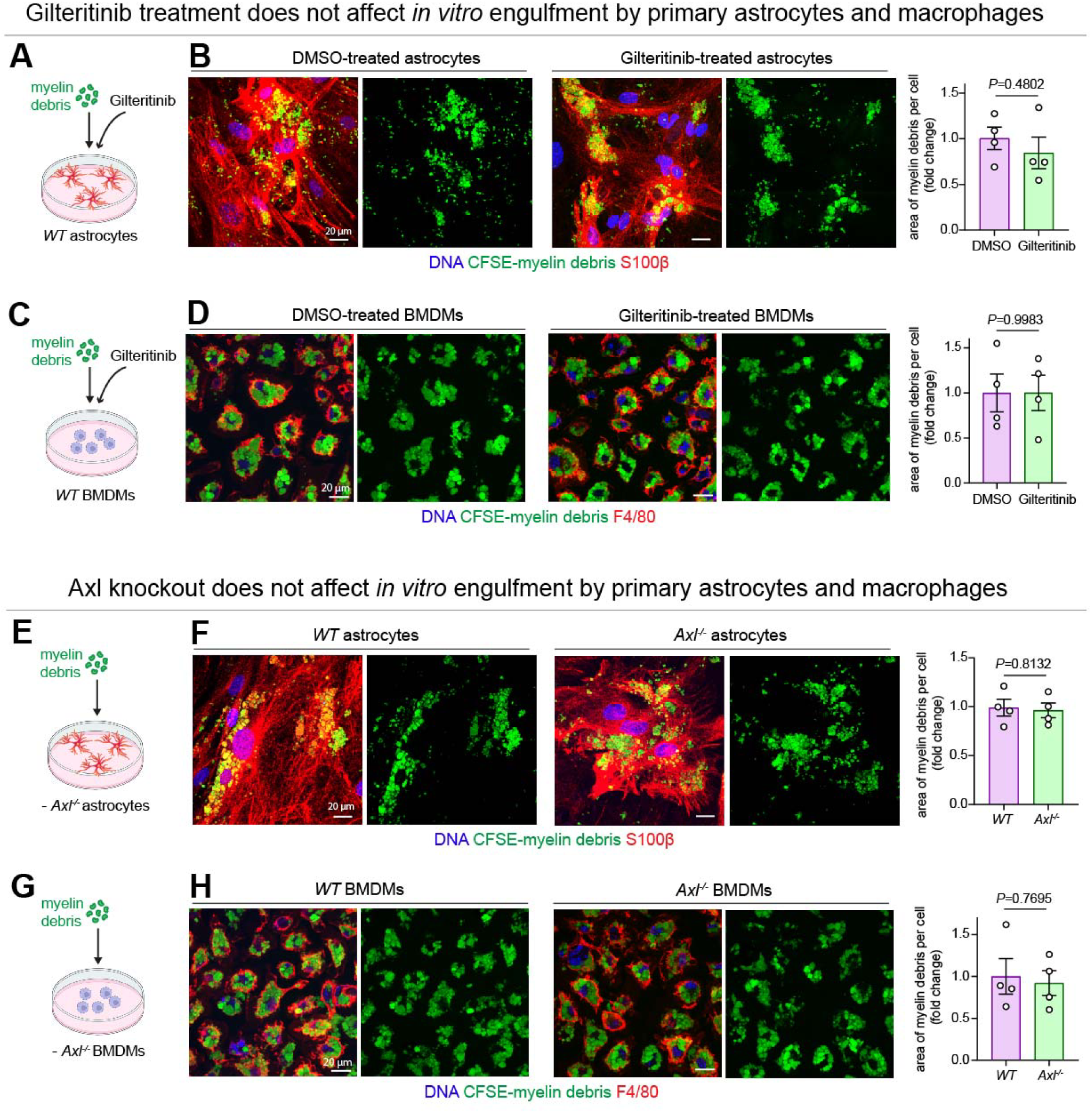
Pharmacological inhibition or genetic deletion of Axl does not affect engulfment of myelin debris by astrocytes or macrophages *in vitro*. **(A)** Schematic illustrating the experimental design for assessing the effect of Gilteritinib on myelin debris uptake by primary astrocytes. **(B)** Representative confocal images and quantification of CFSE-labeled myelin debris uptake by primary astrocytes treated with vehicle or Gilteritinib (1 μM) for 24 h. Astrocytes were identified by S100β immunostaining. Scale bar, 20 μm. Quantification shows the intracellular area of CFSE-positive myelin debris per cell. Data are presented as mean ± SEM (*n* = 4 biological replicates). Statistical significance was determined by an unpaired two-tailed *t* test. **(C)** Schematic illustrating the experimental design for assessing the effect of Gilteritinib on myelin debris uptake by bone marrow-derived macrophages (BMDMs). **(D)** Representative confocal images and quantification of CFSE-labeled myelin debris uptake by BMDMs treated with vehicle or Gilteritinib (250 nM) for 24 h. Macrophages were identified by F4/80 immunostaining. Scale bar, 20 μm. Quantification shows the intracellular area of CFSE-positive myelin debris per cell. Data are presented as mean ± SEM (*n* = 4 biological replicates). Statistical significance was determined by an unpaired two-tailed *t* test. **(E)** Schematic illustrating the experimental design for comparing myelin debris uptake by primary WT and *Axl^−/−^* astrocytes. **(F)** Representative confocal images and quantification of CFSE-labeled myelin debris uptake by primary WT and *Axl^−/−^* astrocytes. Astrocytes were identified by S100β immunostaining. Scale bar, 20 μm. Quantification shows the intracellular area of CFSE-positive myelin debris per cell. Data are presented as mean ± SEM (*n* = 4 biological replicates). Statistical significance was determined by an unpaired two-tailed *t* test. **(G)** Schematic illustrating the experimental design for comparing myelin debris uptake by WT and *Axl^−/−^*BMDMs. **(H)** Representative confocal images and quantification of CFSE-labeled myelin debris uptake by WT and *Axl^−/−^* BMDMs. Macrophages were identified by F4/80 immunostaining. Scale bar, 20 μm. Quantification shows the intracellular area of CFSE-positive myelin debris per cell. Data are presented as mean ± SEM (*n* = 4 biological replicates). Statistical significance was determined by an unpaired two-tailed *t* test.

**Figure S24 – related to Figure 7.**
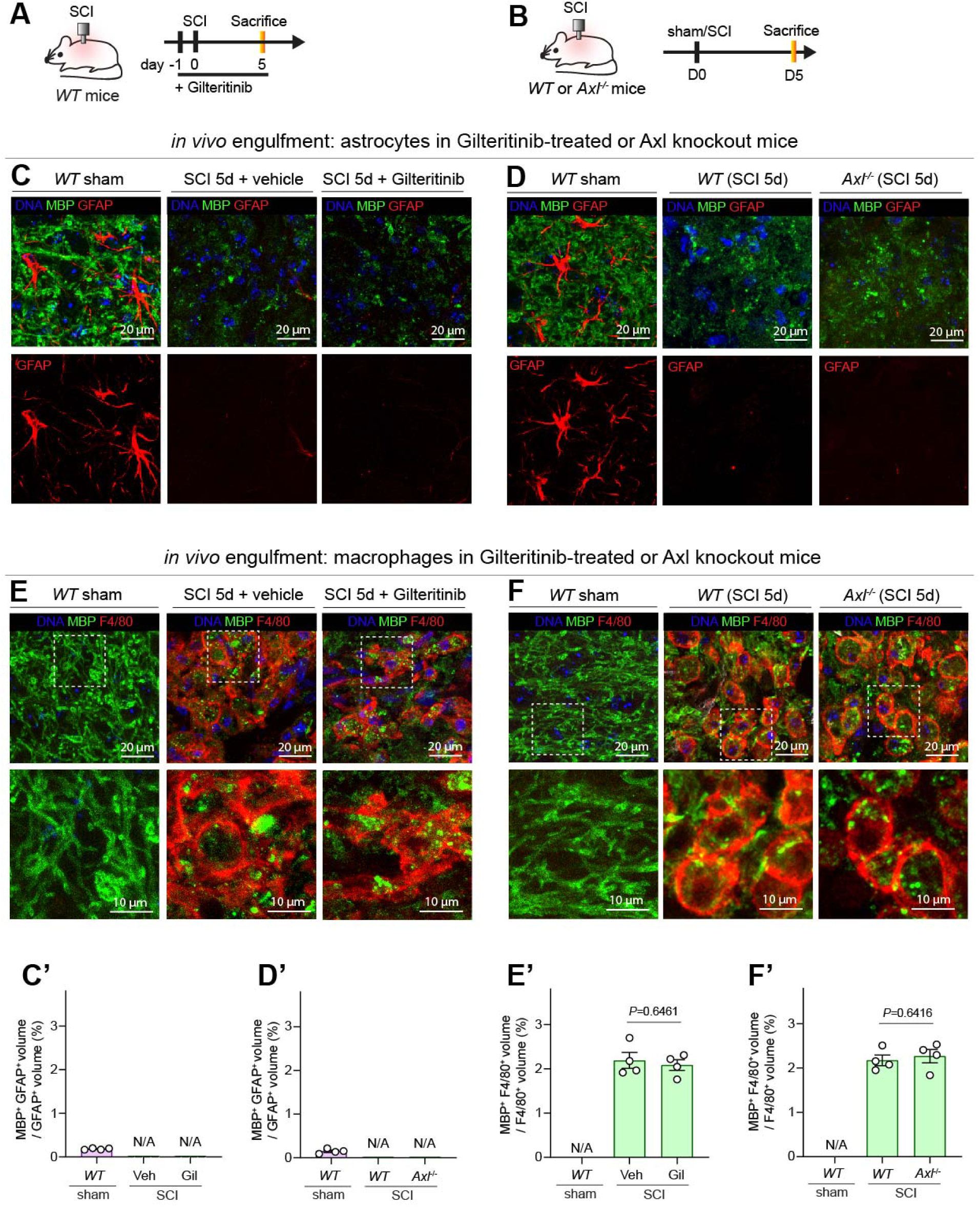
Pharmacological inhibition or genetic knockout of Axl does not affect engulfment of myelin debris by astrocytes or macrophages *in vivo*. **(A, B)** Experimental schemes for assessing the effects of Gilteritinib treatment **(A)** or Axl knockout **(B)** on myelin debris phagocytosis following spinal cord injury (SCI). **(C–F)** Representative confocal images showing myelin debris uptake (MBP staining) by GFAP-positive astrocytes **(C, D)** and F4/80-positive macrophages **(E, F)** in spinal cords from sham-operated mice and mice at 5 days after SCI following Gilteritinib treatment **(C, E)** or Axl knockout **(D, F).** Boxed regions show enlarged views in (E) and (F). Note that astrocytes are largely absent from the SCI lesion core. Scale bar: 20 µm; 10 µm (enlarged views). **(C**′**–F**′) Quantification of the relative intracellular volume of engulfed myelin debris in astrocytes **(C**′**, D**′) and macrophages **(E**′**, F**′) from sham-operated and SCI mice with the indicated treatments or genotypes. Data are presented as mean ± SEM (n = 4 mice). Statistical significance was determined by an unpaired two-tailed *t* test.

**Figure S25 – related to Figure 7.**
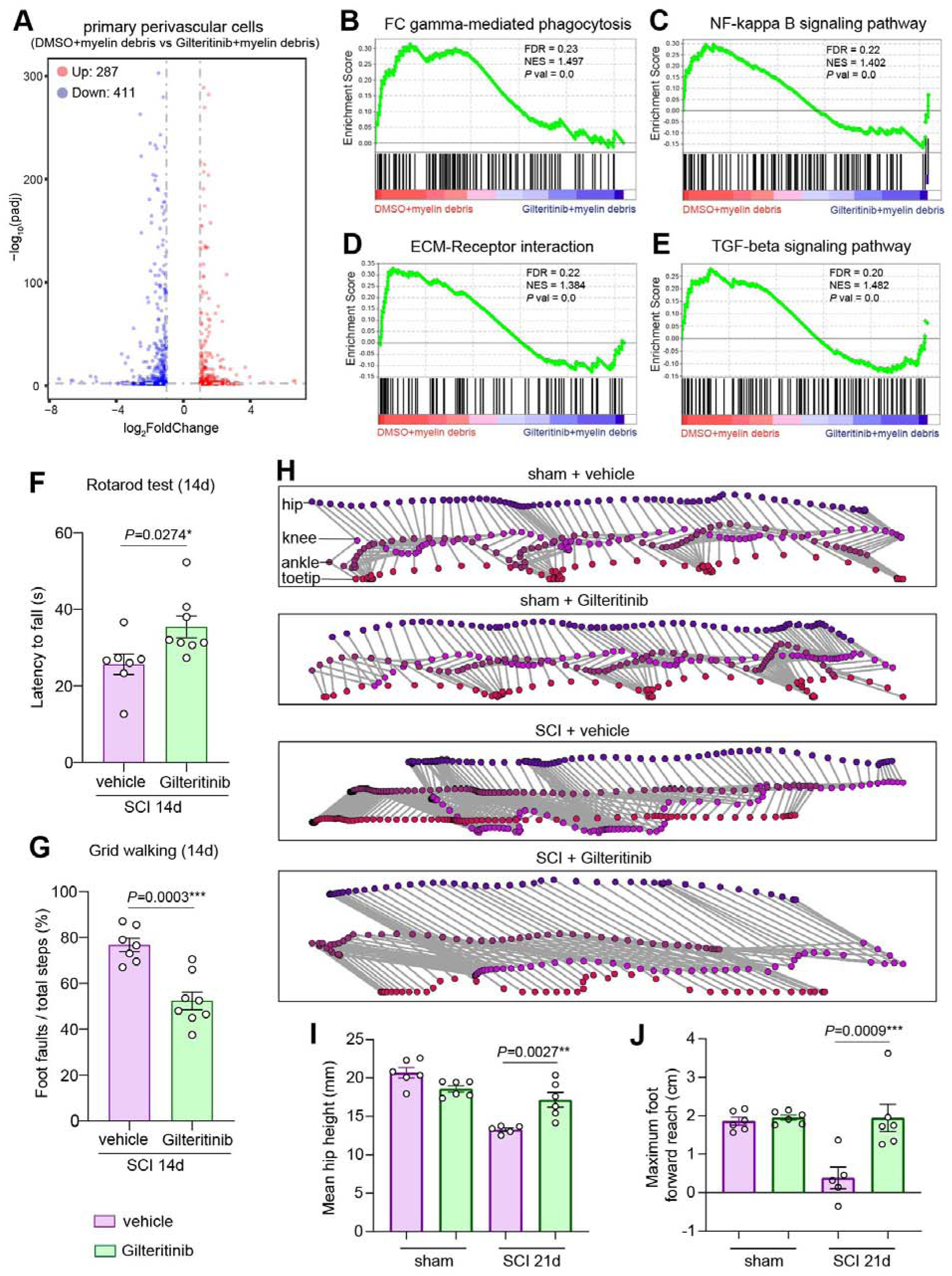
Gilteritinib treatment alters the transcriptomic profile of myelin debris-laden perivascular cells and improves functional recovery after SCI. **(A)** Volcano plot showing differentially expressed genes (DEGs) in myelin debris-laden primary perivascular cells treated with Gilteritinib or vehicle. Genes with |log_2_FoldChange| > 1 and FDR-adjusted *P* < 0.05 were considered differentially expressed. **(B–E)** Gene Set Enrichment Analysis (GSEA) of the indicated pathways in myelin debris-laden primary perivascular cells treated with Gilteritinib or vehicle. Normalized enrichment score (NES), false discovery rate (FDR), and *P* values are indicated. **(F)** Quantification showing latency to fall in the Rotarod test in sham-operated and SCI mice 14 days after treatment with vehicle or Gilteritinib. Data are presented as mean ± SEM (*n* = 7 mice for vehicle, *n* = 8 mice for Gilteritinib). Statistical significance was determined by an unpaired two-tailed *t* test. **(G)** Quantification showing the percentage of faulty foot placements in the grid-walking test in sham-operated and SCI mice 14 days after treatment with vehicle or Gilteritinib. Data are presented as mean ± SEM (*n* = 7 mice for vehicle, *n* = 8 mice for Gilteritinib). Statistical significance was determined by an unpaired two-tailed *t* test. **(H)** Representative images showing markerless tracking and hindlimb kinematic reconstruction in sham-operated and SCI mice 21 days after treatment with vehicle or Gilteritinib. Colored traces indicate the positions of the hip, knee, ankle, and toe joints. **(I, J)** Quantification showing hindlimb kinematic parameters, including mean hip height **(I)** and maximum forward foot reach **(J)**, in sham-operated and SCI mice 21 days after treatment with vehicle or Gilteritinib. Data are presented as mean ± SEM (*n* = 5–6 mice). Statistical significance was determined by two-way ANOVA followed by Holm–Šídák’s post hoc test.

**Figure S26.**
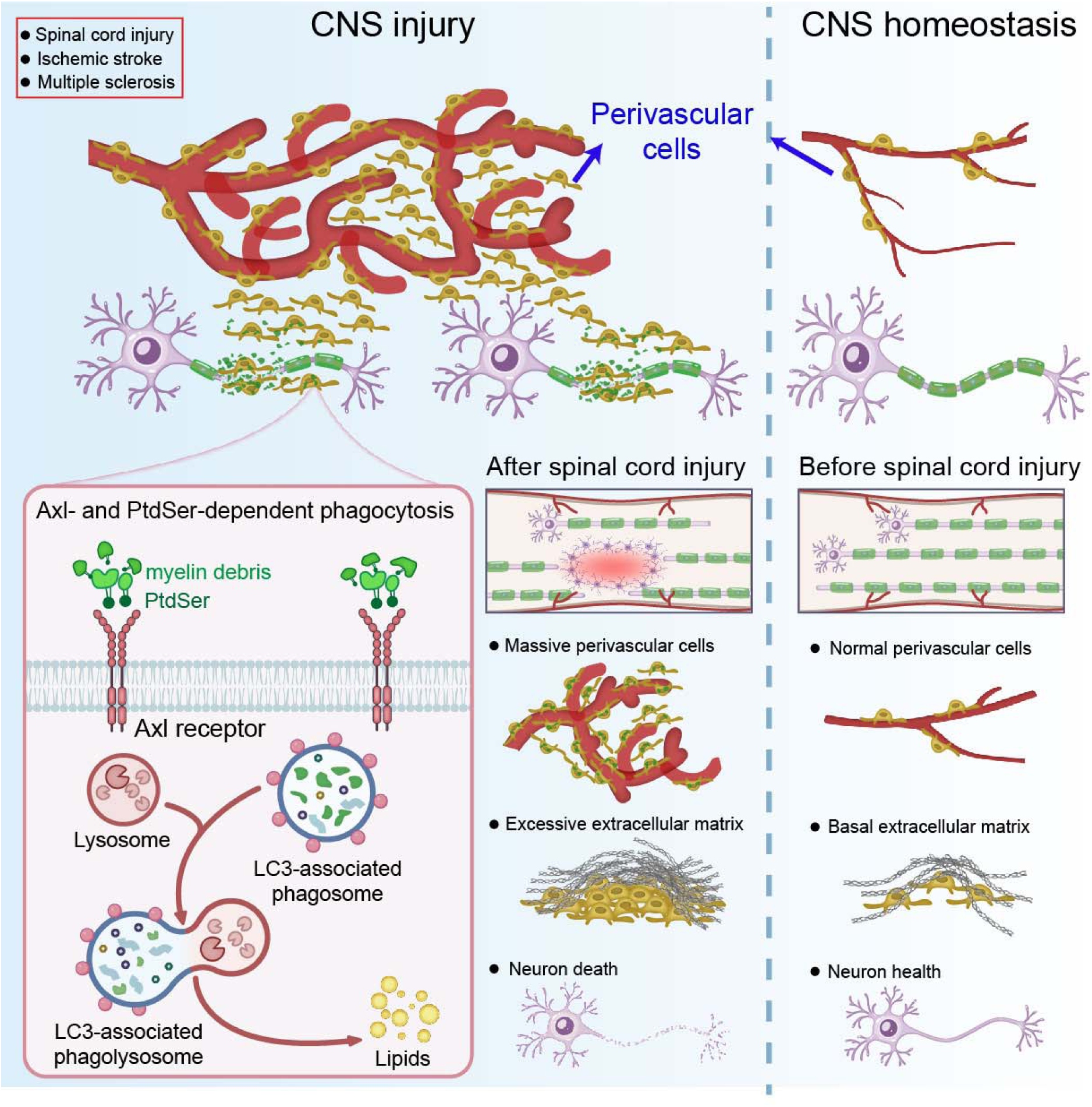
Summary and proposed working model. We identify perivascular cells, a vasculature-associated cell type traditionally regarded as structural components of the neurovascular unit, as previously unrecognized phagocytes that actively sense CNS injury. In multiple mouse models of CNS demyelinating pathology, these disease-associated perivascular cells rapidly expand and engulf tissue debris. Their phagocytic activity is initiated through recognition of phosphatidylserine (PtdSer) “eat-me” signals via the TAM receptor Axl. It is coupled to efficient cargo degradation mediated by LC3-associated degradative pathway characterized by single-membrane phagolysosomal structures, independent of canonical autophagy. Engulfment of lipid-rich myelin debris subsequently fuels perivascular cell proliferation and drives extracellular matrix production, thereby promoting lesion expansion and fibrotic scar formation. Together, these findings define perivascular cells as active participants in CNS injury and highlight Axl-dependent phagocytosis as a central pathway linking debris clearance to pathological tissue remodeling.

